# Circuit inhibition promotes the dynamic reorganization of prefrontal task encoding to support cognitive flexibility

**DOI:** 10.1101/2025.08.09.669414

**Authors:** Carlos A. Johnson-Cruz, Kathleen K.A. Cho, Vikaas S. Sohal

## Abstract

The mammalian prefrontal cortex encodes variables related to goal-directed behavior, and enables flexibility during environmental changes, making it critical to understand how the dynamic updating vs. stable maintenance of different encodings contribute to behavioral adaptation. We addressed this by comparing prefrontal encoding during successful adaptation vs. maladaptive perseveration. Specifically, we studied mutant (*Dlx5/6^+/-^*) mice, which have dysfunctional parvalbumin-expressing inhibitory interneurons and perseverate in a rule shifting task. We measured mPFC activity patterns using microendoscopic calcium imaging, then used linear classifiers and neural networks to compare representational geometries in wild-type mice and *Dlx5/6^+/-^* mutants before, during, and after benzodiazepine treatment, which persistently rescues their rule shift learning. The encoding of correct vs. incorrect trial outcomes rapidly shifts as mice successfully learn new cue-reward associations, but becomes more stable when mutant mice perseverate. We also find activity patterns that normally distinguish learning of the initial association vs. rule shift, but become diminished during perseveration. Finally, during perseveration, outdated representations are inappropriately reinstated, not just passively maintained. These results reveal prefrontal contributions to flexible behavior driven by the dynamic reorganization of abstract rule representations, rather than stable reinforcement signals.

## INTRODUCTION

The ability to update behavioral strategies to adapt to environmental changes is essential for survival in a complex world, but is impaired in many neuropsychiatric conditions. In particular, cognitive flexibility, as measured by tasks such as the Wisconsin Card Sorting Task (WCST), is one of the cognitive domains most impaired in schizophrenia (Green 2006). Tasks like the WCST require individuals to make decisions by selectively attending to specific cues. Individuals must learn rules that guide these decisions, identify when these rules change, and update their behavior accordingly.

The prefrontal cortex plays a key role in this type of cognitive flexibility, but the exact nature of its computational contribution remains unclear. Specifically, the prefrontal cortex could encode rules, outcomes, levels of uncertainty, actions (e.g., go vs. no-go), the value associated with actions or cues, or many other behavioral variables. Different types of encoding are hypothesized to support various potential functional roles for the prefrontal cortex. For example, robust encoding of rules might focus attention on specific cues, whereas robust encoding of outcomes could provide feedback that triggers learning. One recent study found that individual prefrontal neurons stably encode responses and outcomes (Spellman et al., 2021). Other studies have emphasized that prefrontal neurons form stable representations of task rules, which become unstable during behavioral transitions (Durstewitz et al., 2010; Karlsson et al., 2012; Malagon-Vina et al., 2018).

The mechanisms through which neural encoding – either stable encoding of specific information related to choices, outcomes, or value, or dynamic representations that correspond to updating internal models – support learning are broadly relevant to many brain regions and aspects of natural intelligence. For example, a recent study highlighted that over the course of learning, hippocampal representations for common elements (i.e., shared locations) associated with two different rules (predicting two different reward locations) become orthogonalized (Sun et al., 2025), illustrating how the manner in which neural representations are updated can inform and constrain the underlying mechanisms for learning.

Several challenges complicate analyses of prefrontal encoding, especially during behavioral transitions. First, prefrontal neurons often exhibit mixed selectivity for many variables (Tye et al., 2024). During behavioral transitions, these variables are inherently confounded; for example, a given trial may be associated with a change in the rule, but also with a particular reward location, level of uncertainty, outcome, and configuration of task-irrelevant cues. As a result, when animals learn new rules rapidly (e.g., in a few trials), it can be challenging to disentangle the contributions of these different factors to single-neuron encoding.

Population-level decoding techniques can aid in illuminating the content of representations associated with prefrontal neural encoding. Using the activity of multiple neurons as input to a classifier and then testing how well trials associated with different combinations of task variables can be separated based on the underlying neural activity provides a valuable test of whether a neuronal population meaningfully encodes specific variables. However, this introduces a second challenge, related to the functional significance of such encoding. Functional significance is typically assessed using causal manipulations such as opto- or chemogenetics to activate or silence specific cell types. However, given that encoding is often distributed across sparse neuronal ensembles, this approach has its limitations, especially in freely-moving animals where precise control of many constituent neurons within a dynamically formed ensemble is not currently feasible.

Here we address these challenges by leveraging a mutant mouse model in which deficits in prefrontal-dependent cognitive flexibility can be persistently reversed. We previously showed that during a prefrontal-dependent rule shifting task, mice heterozygous for the transcription factors *Dlx5* and *Dlx6* perseverate and have difficulty learning new rules based on previously task-irrelevant cues. We linked this behavioral impairment to the dysfunction of parvalbumin (PV) interneurons, specifically their inability to synchronize at gamma-frequencies (∼40 Hz) (Cho et. al., 2015; Cho et al., 2020). In particular, disrupting this gamma-synchrony by delivering 40Hz optogenetic stimulation to prefrontal PV interneurons *out-of-phase* across the hemispheres was sufficient to induce perseverative phenotypes in previously normal mice (Cho et al., 2020). (Disrupting synchrony by inhibiting callosal PV+ projections, which are necessary for gamma synchrony, produces a similar effect (Cho et al., 2023)). By contrast, delivering the same 40Hz pattern of stimulation *in-phase* across the hemispheres to restore gamma-synchrony was sufficient to rescue perseveration in *Dlx5/6^+/-^* mutants (Cho et al., 2020). Remarkably, this pro-cognitive effect of in-phase 40Hz stimulation is persistent, and can be reproduced pharmacologically, using low (sub-sedative and sub-anxiolytic) doses of the benzodiazepine clonazepam to enhance PV interneuron gamma synchrony.

Here, we confirm that clonazepam leads to a persistent rescue of behavior and gamma synchrony in *Dlx5/6^+/-^* mutant mice. We record activity from populations of prefrontal neurons using microendoscopic Ca2+ imaging, then examine this high-dimensional population activity using linear classifiers and neural network autoencoders to understand how representations evolve across different task stages. We compare classifier performance in wild-type mice, and *Dlx5/6^+/-^* mutants before, during and after treatment with clonazepam. This allows us to identify how specific variables are normally encoded during behavioral transitions, and cases in which this encoding is disrupted in *Dlx5/6^+/-^* mutants but restored when cognitive flexibility is normalized by clonazepam treatment. In this way, we test ideas about what types of information are robustly encoded, and highlight specific aspects of frontal encoding that are linked to successful cognitive flexibility based on a three-fold validation: they are significantly perturbed in mutants relative to wild-type, rescued by clonazepam acutely, and demonstrate a persistent rescue that outlasts the acute treatment with clonazepam.

## RESULTS

### Low-dose clonazepam persistently rescues cognitive impairments and gamma synchrony in *Dlx5/6^+/-^* mutant mice

We studied the behavior of mice in a rule-shifting task that measures specific aspects of cognitive flexibility (Cho et al., 2015, 2020, 2023). In the task, mice learn associations (a rule) in which one of 4 sensory cues across 2 sensory dimensions (i.e., 2 odors and 2 textured digging media) signals the location of a hidden food reward. On each trial, mice are presented with 2 bowls, each containing 1 texture and 1 odor cue – there are thus 4 unique combinations formed by the 2 texture and 2 odor cues (**Figure 1A**). Once mice learn the *initial association* (IA) between one cue and the hidden reward, the cue-reward association undergoes a *rule shift* (RS) in which a new cue from the other sensory dimension now indicates the location of the hidden food reward. Mice must recognize that this change has occurred, ignore the previously relevant sensory cues, and adapt their strategy to obtain food rewards by paying attention to the previously irrelevant sensory cues. (We refer to the trials after the new rule begins as the rule shift period). Prior work from our lab and others has demonstrated that this type of extra-dimensional rule shifting task depends on the medial prefrontal cortex (mPFC) (Bissonnette et al., 2008), evokes increases in PV IN activity and gamma-frequency (∼40 Hz) synchronization following rule-shift errors (Cho et al., 2015, 2020), and is sensitive to disruptions of mPFC PV IN function (Canetta et al., 2016; Canetta et al., 2022; Cho et al., 2020; Cho et al., 2023).

**FIGURE 1:**
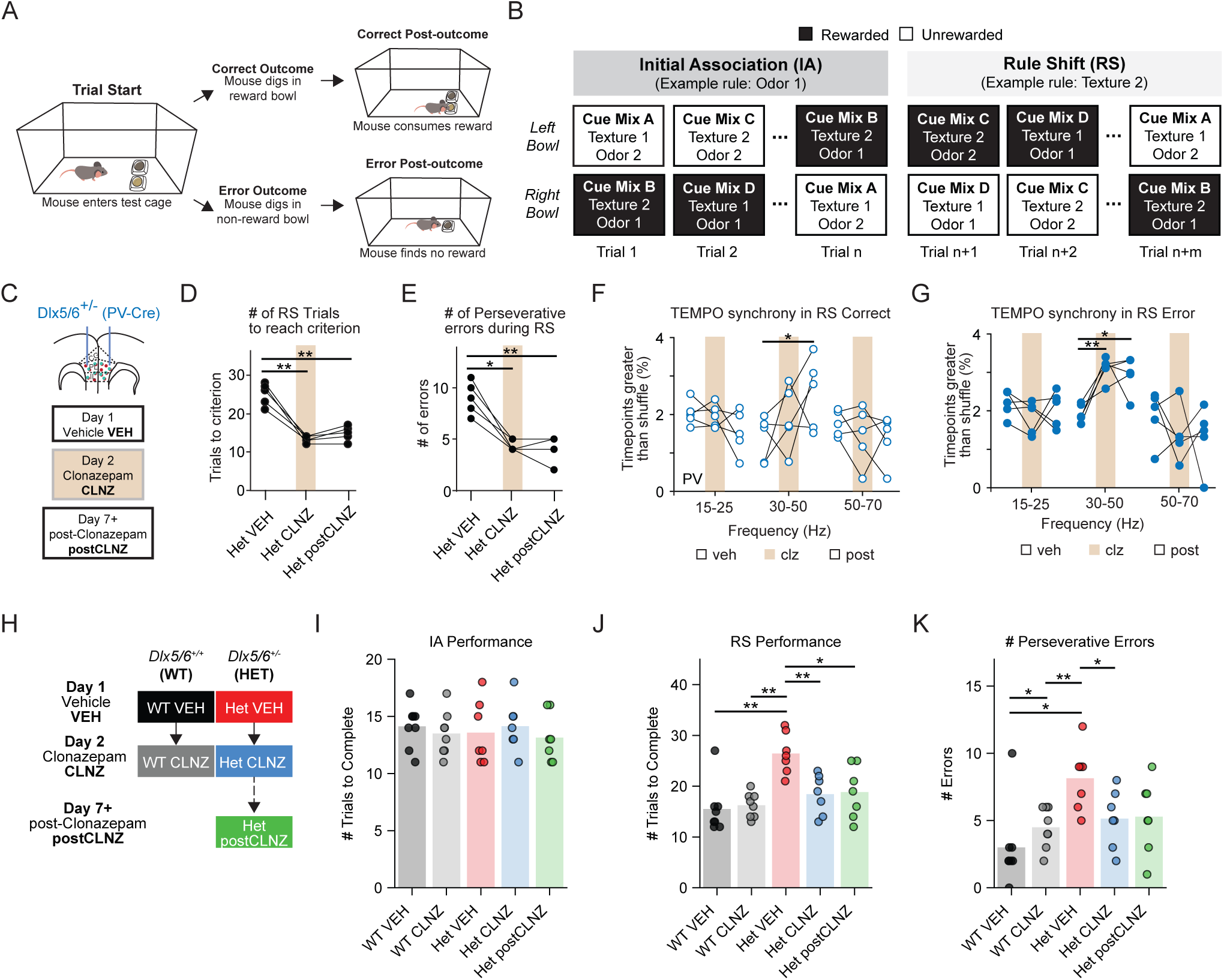
Clonazepam rescues deficits in rule shift learning and gamma synchrony in *Dlx5/6*^+/-^ mice. A. On each trial, mice are allowed to dig in only one of two bowls to find a hidden food reward, which triggers the trial outcome. During the post-outcome period following correct trials, mice consume food rewards. During the post-outcome period following error trials, the rewarded bowl is removed from the test cage. Regardless of outcome type, mice can freely explore the test cage for at least 15 seconds before being moved into the holding cage. B. Schematic showing a sample set of trials from a rule-shifting session. A rule assigns food reward to one of the two bowls. The rule is an association between reward and one sensory cue (out of 2 odors and 2 textures). Mice learn two rules in each session. In this example, the first rule (Initial Association, IA) is Odor 1, so the Odor 1-scented bowl always contains food reward. After mice reach the learning criterion (8/10 consecutive trials correct), the rule changes to an association based on a cue from the other sensory dimension (Rule Shift, RS). C. Schematic of photometry implants and experimental paradigm for experiments done as part of Cho et al., 2020. *Dlx5/6^+/-^*mutant and wild-type mice were tested on the rule-shift task over multiple days: Day 1 - vehicle injection, Day 2 - clonazepam injection (CLNZ), and Day 3 - at least 7 days later after CLNZ treatment (postCLNZ). D. (Experiments originally performed in Cho et al., 2020): CLNZ treatment significantly reduces the # of rule shift trials required for *Dlx5/6^+/-^*mice to reach the learning criterion both acutely and persistently. N = 5 *Dlx5/6^+/-^*mice. E. (Experiments originally performed in Cho et al., 2020): CLNZ treatment significantly reduces the # of perseverative errors by *Dlx5/6^+/-^*mice (n = 5). F. (Experiments originally performed in Cho et al., 2020): In *Dlx5/6^+/-^* mice, CLNZ treatment leads to a significant increase in interhemispheric gamma synchrony at the postCLNZ timepoint (measured by the fraction of timepoints with correlations greater than the 95^th^ percentile of the shuffled distribution, for signals filtered between 30-50Hz) following correct outcomes. N = 5 *Dlx5/6^+/-^* mice. G. (Experiments originally performed in Cho et al., 2020): In *Dlx5/6^+/-^* mice, CLNZ treatment leads to a significant increase in interhemispheric gamma synchrony (measured by the fraction of timepoints with correlations greater than the 95^th^ percentile of the shuffled distribution, for signals filtered between 30-50Hz) following error outcomes, both acutely and persistently. N = 5 *Dlx5/6^+/-^*mice. H. Experimental design for calcium imaging experiments using CLNZ: WT and mutant (*Dlx5/6^+/-^* HET) mice performed the rule-shifting task over multiple days. Each group received vehicle (VEH) injections and clonazepam (CLNZ) injections on Days 1 and 2, respectively. HET mice were tested again 7+ days after CLNZ. I. HET (n = 7) and WT mice (n = 8) learn the Initial Association (IA) in a similar number of trials. J. HET mice (n = 7) take significantly more trials than WT (n = 8) to learn the Rule Shift (RS). This abnormality is ameliorated during and after CLNZ treatment. K. HET mice make more perseverative errors than WT during the RS. This is reversed by CLNZ treatment. Panels D-E: p-value from Tukey’s multiple comparisons test. Panels F-G: p-value from 2-sided t-tests with Bonferroni correction. Panels I-K: p-value from Mann-Whitney U tests. All panels: \**P* < 0.05, \*\**P* < 0.01, \*\*\**P* < 0.001

As discussed above, in one such model of PV IN dysfunction – mice heterozygous for *Dlx5* and *Dlx6* (*Dlx5/6^+/-^*, ‘HET’) – optogenetically restoring gamma synchrony persistently reverses deficits in rule shift learning. Furthermore, we found that treating *Dlx5/6^+/-^*HET mice with clonazepam not only rescued RS learning (**Figure 1D-E**) but also restored interhemispheric gamma-frequency (∼40 Hz) synchronization between prefrontal PVINs (Cho et al., 2020). To confirm that these therapeutic effects of CLNZ, like those of optogenetic stimulation, are persistent, we performed additional analyses of data previously collected as part of our prior study (Cho et al., 2020). Specifically, to quantify gamma synchrony between mPFC PV INs during rule shift learning, we had used fiber photometry to measure fluorescence from the genetically encoded voltage indicator Ace2N-4AA-mNeon (Ace-mNeon) expressed in PVINs, as well as from a reference fluorophore (tdTomato). After filtering signals between 30-50 Hz to resolve gamma-frequency activity, we computed correlations between voltage signals collected from the left and right mPFC. We used the fraction of timepoints with correlations greater than those obtained using time-shuffled data as a synchrony metric. Using this approach, we found that interhemispheric gamma synchrony between prefrontal PVINs typically increases following RS errors (i.e., when mice received feedback that the previously learned association was no longer valid). This increase in interhemispheric PV gamma synchrony was absent at baseline in *Dlx5/6^+/-^*HET mice, but acutely restored by CLNZ (**Figure 1G**).

That study had also collected data during an additional RS, performed >10 days after the CLNZ treatment. As shown in Figure 1E-G, CLNZ improved RS learning (measured by the number of trials to reach the learning criterion) and increased interhemispheric PVIN gamma synchrony following RS errors on this post-CLNZ day. Thus, our previously collected data showed that CLNZ elicits a *persistent* restoration of RS learning and associated increases in interhemispheric PVIN gamma synchrony.

### Calcium imaging of neuronal ensembles as mice learn rule shifts

We used head-mounted microendoscopes (Inscopix, nVoke2.0) for single-photon calcium imaging in freely moving adult mice as they learned initial associations and rule shifts. Mice received a unilateral GRIN lens implant and AAV9-synapsin-GCaMPF7f injections in mPFC (**Figure 2A-B**). We recorded behavioral performance and neural activity for at least 2 rule-shifting sessions on consecutive days. On Day 1, mice received a control intraperitoneal (i.p.) injection of vehicle 30 minutes prior to beginning the task (VEH condition). On Day 2, mice received a sub-sedative and sub-anxiolytic dose of the benzodiazepine clonazepam (CLNZ; 0.0625 mg/kg i.p.). We previously showed that CLNZ at this dose rescues rule shifting deficits in *Dlx5/6^+/-^* (HET) mice (Cho et al., 2015). After Day 2, we waited at least seven days (and in many cases substantially longer), then imaged mutant mice during a final rule-shift test (post-CLNZ) (**Figure 1C**). WT mice were not retested following CLNZ washout. We categorize datasets by their specific intersection of genotype (WT or HET) and treatment (VEH, CLNZ, or post-CLNZ), and analyze behavior and neural data from these 5 groups.

**FIGURE 2:**
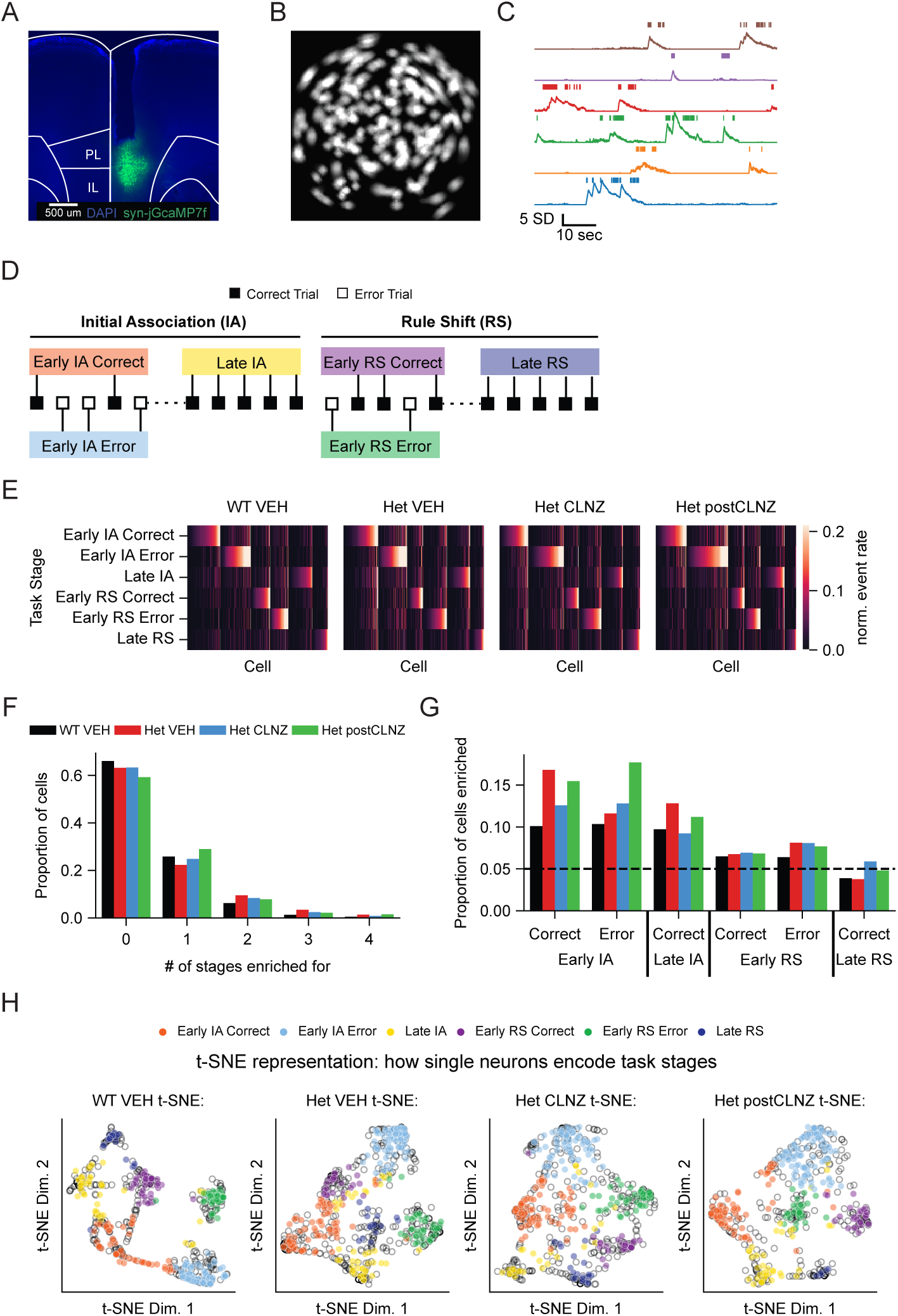
Recruitment of prefrontal neurons during different task stages in WT and *Dlx5/6*^+/-^ mice. A. Representative image showing mPFC implant location and neurons transfected with syn-jGCaMP7f (green) amidst cell bodies stained with DAPI (blue). B. Sample field of view containing EXTRACT-detected spatial components (putative neurons). C. Example dF/F calcium traces from a subset of detected ROIs. Frames classified as containing a calcium event are denoted by the vertical ticks above each trace. D. Example schematized sequence of trial types from an example mouse. E. Matrix containing the mean stage response by neuron per genotype-treatment group. Each column contains the mean normalized activity of one cell in each task stage. Cells sorted based on the trial type / task stage in which they were most active. (cells from recordings without early IA error trials were not included for visual clarity). F. The number of trial types / task stages in which each cell exhibits significantly elevated activity (enrichment) is not significantly different across genotype-treatment conditions. G. The fraction of cells with significantly elevated activity (enrichment) as a function of trial type / task stage does not significantly vary across genotype-treatment conditions. The dashed line indicates the level of enrichment expected by chance (0.05). H. t-SNE projections of the activity of each cell across trial types / task stages. Each point represents one cell. Transparent black circles are non-enriched cells (not significantly active in any task stage). Filled circles indicate enriched cells and are color coded according to the stage in which that cell is most active.

During Day 1 of testing (VEH condition), HET mice (n = 8) and their WT littermates (n = 7) learned the initial association (IA) in a similar number of trials (p = 0.6) (**Figure 1I**). However, during the rule shift phase of the task, HET mice required significantly more trials to consistently locate reward (WT VEH vs Het VEH, RS TTC p = 0.0071, MWU), and made significantly more perseverative errors (errors where mice selected bowls corresponding to the cue that was rewarded during the IA period) than WT (p = 0.012, MWU), aligning with our previous findings (Cho et al., 2015) (**Figure 1J-K**). This cognitive impairment has been previously observed to persist in HET mice over at least three consecutive days of rule shifting (Cho et al., 2015). Sex did not significantly affect task performance or the number of perseverative errors made for any group (Supplementary Fig. 1).

Administering the benzodiazepine clonazepam (CLNZ) 30 minutes before the task (i.e., prior to learning the initial association) improved the performance of HET mice during the rule shift (<u>mean trials to RS criterion:</u> 26.4 +/- 1.5 in VEH vs. 18.4 +/- 1.5 in CLNZ, p = 0.0072, MWU, n = 8 mice) but did not significantly alter performance in WT mice (<u>mean trials to RS criterion:</u> 15.5 +/- 1.7 in VEH vs. 16.3 +/- 0.9 in CLNZ, p = 0.19, MWU, n = 7 mice). Our previous work has shown that the RS performance of HET mice does not improve with repeated testing in the absence of other manipulations (Cho et. al., 2015). However, the CLNZ-induced improvement in the RS performance of HET mice persisted 10-64 days later, in the absence of any additional interventions (trials to RS criterion: 26.4 +/- 1.5 in VEH vs. 18.9 +/- 1.9 postCLNZ, p = 0.017, MWU, n = 8 mice).

### Prefrontal neurons are selectively recruited during specific task stages

Successful learning of a rule shift could result from changes in the recruitment of individual neurons and their organization into neuronal ensembles. To assess these changes, we divided task sessions into six stages/trial types and quantified the recruitment of individual neurons during each task stage. Specifically, we focused on the “Early” and “Late” stages of learning for each rule, defined as the first 5 and last 5 trials, respectively, of either the initial association (IA) or rule shift (RS). The early IA or RS periods were further subdivided into correct or error trials (Figure 2A). The last 5 trials of the IA or RS portions of the task were mainly comprised of correct trials (98% and 92% for IA and RS, respectively). As a result, we excluded error trials occurring in the last 5 trials from the “late IA” and “late RS” groups (to focus on stable behavior). In this way, we defined 6 different task stages (trial types): early IA correct, early IA error, late IA; early RS correct, early RS error, late RS.

We recorded a total of 866-1080 cells per treatment condition across 7 WT and 8 HET mice. We analyzed the DF/F time series from each cell to identify frames in which it was classified as ‘active’ (**Figure 2C**; Methods). Mice take variable amounts of time on each trial to choose one bowl to dig in. Our previous work highlighted abnormal PV interneuron recruitment and gamma synchrony occurring in HET mice during the post-outcome period, after mice dig in a bowl and then either find or fail to obtain a reward (Cho et al., 2020). Therefore, we focused our analysis on the period immediately preceding each dig, and the subsequent post-dig period (-3 to +15 sec relative to the dig). (Note: Our study seeks to detect changes in HET mice, relative to WT, that CLNZ persistently reverses. Therefore, to facilitate relevant visual comparisons, most of our main figures do not show the WT CLNZ group. However, data from this group is included in Supplementary Figures 1-2, and differences from WT VEH are described in a dedicated subsection of the Results).

Individual neurons exhibited recruitment that was largely specific for one of the 6 trial stages listed above (**Figure 2E**). Correspondingly, different patterns of activity were associated with each trial stage. The average level of activity, defined as the average fraction of frames in which each cell was active, was 1.4 - 3.1% in WT mice. Cells were most active on average during early IA error trials and least active during the late RS (Supplementary Figure 1C). Average activity levels were significantly higher in HET mice compared to WT mice in the early IA correct, late IA, and early RS error task stages. CLNZ treatment caused a significant decrease in active frames in HET mice but not in WT mice during early IA correct and late IA (Supplementary Figure 1C).

Cortical neural ensembles facilitate complex computation via dynamic and patterned co-activity (Carrillo-Reid et al., 2019, Yuste 2015). In particular, prefrontal ensembles encode information that can be difficult to discern from single-cell activity due to the mixed selectivity and relatively low signal-to-noise ratios of single prefrontal neurons (Fusi et al., 2024). To quantify the membership and activity dynamics of mPFC neural ensembles during rule-shifting, we used permutation testing to determine the statistical significance of activation per task stage via comparison to a null distribution of 1000 circularly shuffled datasets (**Figure 2E**). We group cells into a stage ensemble if their activity exceeded the 95th percentile of shuffle data in that task stage (i.e., are enriched in said stage).

Approximately 60% of mPFC cells were not significantly recruited by any task stage (**Figure 2F**). Approximately 20% of cells significantly increased activity in exactly 1 task stage, and ∼15% did so for two or more task stages (**Figure 2F**). We observed no significant difference in the total number of cells that significantly increased their task stage activity between treatment or genotype groups (WT VEH vs. Het VEH: p = 0.2; Het VEH vs. Het CLNZ: p = 0.99; χ^2^ test). On average, the ensemble associated with a given task stage comprised 60-80% of all cells active on a given trial during that stage (**Supplementary Figure 2**). Task stage ensembles could also be visualized using t-SNE as distinct clusters within a state space defined by the average activity of a neuron on different task stages (**Figure 2H**).

Identifying task stage ensembles allows us to examine how neurons with specific tuning stably encode specific task-related information and/or how their encoding evolves during learning. For example, it is possible that cells previously strongly recruited during early IA correct or error trials might exhibit stable outcome-related encoding during the RS, even though they are recruited more modestly during the RS. Based on this logic, subsequent analyses will examine the activity of a specific group of neurons – a task stage ensemble – on different trial types. E.g., we could examine the activity of the early IA error ensemble on early RS correct and early RS error trials to determine whether this group of neurons stably encode trial outcomes. We examine encoding by specific ensembles of neurons (defined by the trial type on which they exhibit significantly elevated activity) to detect patterns of normal or abnormal encoding that may manifest within specific subsets of neurons, rather than in the broader population.

Because we name ensembles based on the task stage during which they exhibit significantly elevated activity (e.g., the ‘early IA correct ensemble’), examining their activity and encoding on other task stages might initially seem odd. However, while ensembles are defined by the task stage/trial type on which they exhibit significantly elevated activity, they also meaningfully modulate their activity during other task stages/trial types (as shown below). Furthermore, from a population standpoint, although we have identified task stage ensembles based on post-hoc analyses, during the task, the brain does not have access to the explicit task stage labels, making it reasonable to assume that various task stage ensembles could contribute to internal processes underlying learning and decision making. Finally, we identify task stage ensembles based on the neurons that significantly increase activity during particular task stages, but obviously, decreases in activity could also encode important task features. For these reasons, we examine the encoding of various task-related variables across multiple task stages/trial types using multiple task stage ensembles, not only the ensemble corresponding to the same task stage.

### *Dlx5/6^+/-^* mutants have abnormally stable trial outcome encoding

To identify patterns of encoding associated with normal or pathological rule shifting, we calculated the *cross-condition generalizability* using linear classifiers (Bernardi et al., 2020). Linear classifiers such as support vector machines (SVMs) utilize the high-dimensional distribution of neural activity to separate activity patterns related to different classes. Thus, training classifiers on data from one condition (e.g., during the IA), then testing them on data from a different condition (e.g., the RS), quantifies the degree to which the geometry of encoding is preserved between these two conditions. Simple neural circuits can implement linear classification, meaning that SVM performance also indirectly indicates the information available to a hypothetical downstream neuron (Boyle et al., 2024)

We first examined whether neural ensembles preserved encoding of outcomes (e.g., correct vs. error trials) across IA and RS task stages. Specifically, we trained SVMs to classify activity patterns observed on early IA correct vs. early IA error trials. Then, we tested model performance at classifying activity patterns observed during early RS correct and error trials as either ‘Correct’ (i.e., more similar to early IA correct) or ‘Error’ (i.e., more similar to early IA error) (**Figure 3A**). We refer to classifier accuracy in this method as *cross-condition generalizability performance* (CCGP) (Bernardi et al., 2020; Boyle et al., 2024). We trained and tested separate classifiers for four different pseudopopulations derived from the early IA correct, early IA error, early RS correct, or early RS error task stage ensembles (pooled across mice). In other words, we identified an ensemble of neurons *significantly recruited during a particular trial type*, then used their activity *during early IA correct and error trials* to train an SVM, and finally tested how that SVM classified activity *from early RS correct vs. error trials*.

**FIGURE 3:**
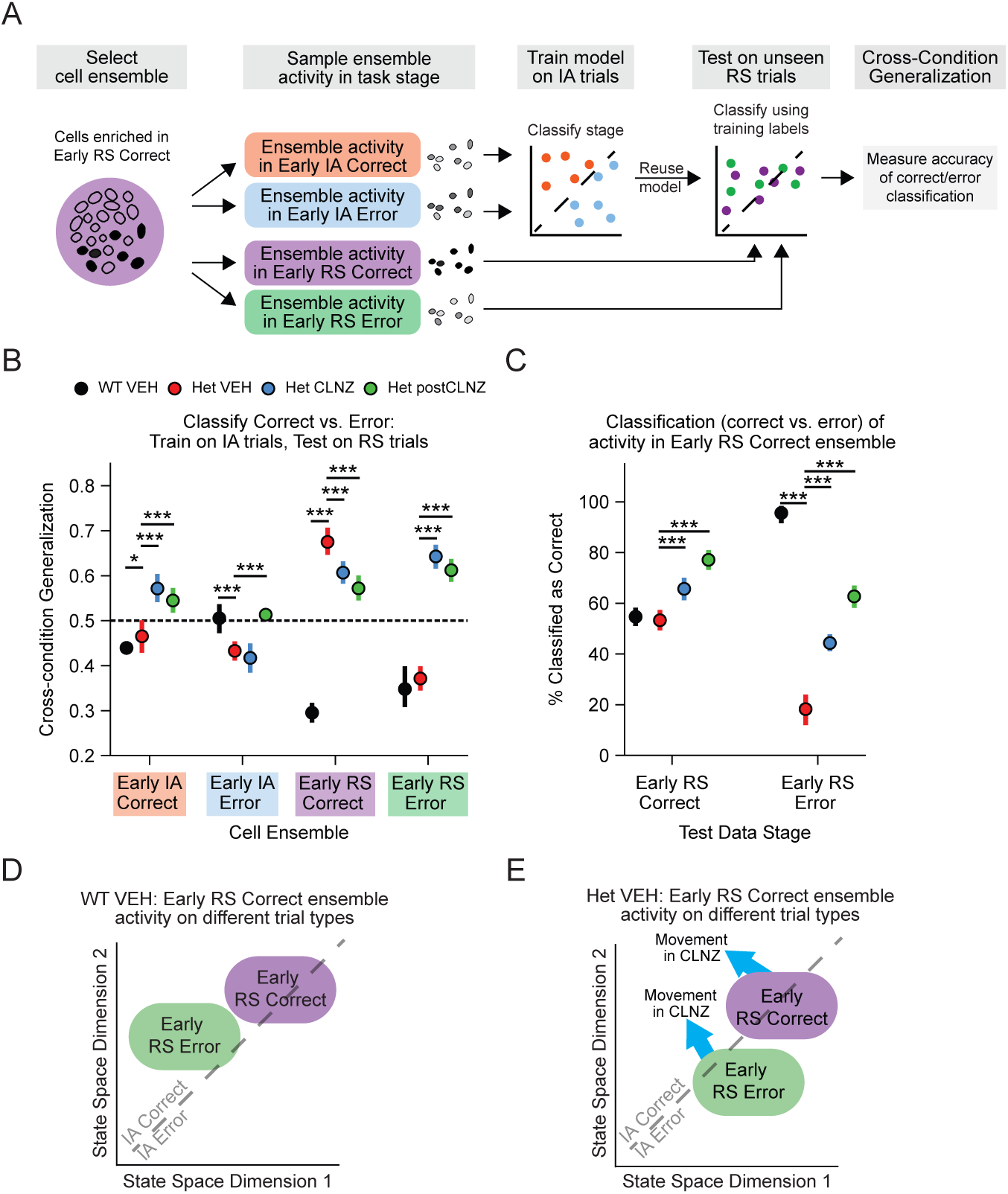
*Dlx5/6*^+/-^ mice have abnormally stable trial outcome encoding. A. Schematic for Cross-Condition Generalization Performance (CCGP). Neuronal ensemble activity is resampled from early IA Correct, early RS Correct, early IA error, and early RS correct. SVMs are trained to classify activity of each ensemble on early IA correct vs. early IA error trials. The trained model is then tested on unseen data: that ensemble’s activity on early RS correct and early RS error trials. CCG Performance computes how well the SVM classifies the test data as ‘Correct’ or ‘Error’. B. The CCGP is plotted for each ensemble (early IA / RS correct / error) and each genotype-treatment condition. Notably, outcomes encoding is never maintained in the WT VEH condition (CCGP never > 0.5). In Het VEH, CCGP becomes abnormally high in the early RS ensemble, which is reversed by CLNZ treatment. C. Percent of samples from early RS correct trials or early RS error trials classified as early IA correct by CCG models trained on early RS correct ensemble data for each genotype-condition. WT VEH activity on early RS error trials is mainly classified as ‘Correct’, but this reversed in the Het VEH condition. CLNZ treatment reduces this abnormality. D. Diagram of the inferred geometry of task stage / trial type activity for early RS correct ensemble in WT VEH mice, relative to the CCG model classification boundary separating activity on early IA correct vs. early IA error trials. In this condition, early RS correct trial activity lies on the decision boundary (resulting in near equal proportions of classifications as Correct vs. Error), whereas activity on early RS error trials lies mainly in the ‘Correct’ region. These positions are estimated from panel C’s quantifications, and distances are not drawn to scale. E. Similar to D, but illustrating evolution of early RS correct ensemble activity in the Het VEH group. Activity on early RS error trials is shifted into the ‘Error’ region. Blue arrows represent the effects of CLNZ treatment. Panels B-C: p-value calculated from Cohen’s U3, which is derived from each comparison’s Cohen’s *d.* Points and error bars show the mean +/- 75%ile of N = 1000 bootstrap distributions per genotype-treatment ensemble. \**P* < 0.05, \*\**P* < 0.01, \*\*\**P* < 0.001.

Surprisingly, in the WT VEH condition, none of these four ensembles maintained outcome encoding from IA to RS (filled black circles in **Figure 3B**). Specifically, SVMs trained on the early IA correct or error ensemble activity from WT VEH mice led to chance-level classification of early RS correct vs. early RS error trial activity as ‘correct’ vs. ‘error’ (early IA correct ensemble accuracy: 0.44 +/- 0.01; early IA error ensemble: WT VEH 0.51 +/- 0.02). Unexpectedly, classification in WT VEH mice based on early RS correct or error ensemble activity was substantially *below-chance* (early RS error ensemble: WT VEH 0.35 +/- 0.02; early RS correct ensemble: 0.29 +/- 0.01).

The only case where generalization was significantly perturbed in the Het VEH condition and altered by CLNZ treatment was when SVMs were trained on early RS correct ensemble activity. Surprisingly, for this ensemble, SVMs trained on Het VEH data had significantly *higher* accuracy than models trained on WT VEH data; this accuracy was significantly decreased (closer to chance levels) in both the Het CLNZ and post-CLNZ conditions (<u>early RS correct ensemble</u> <u>generalization accuracy:</u> WT VEH 0.29 +/- 0.01, Het VEH 0.68 +/- 0.01, Het CLNZ 0.61 +/- 0.01, Het postCLNZ 0.57 +/- 0.01. WT VEH vs Het VEH Cohen’s *d* = 32, p < 10^-15^; Het VEH vs Het CLNZ Cohen’s *d* = 5.4, p = 3×10^-8^; Het VEH vs Het postCLNZ Cohen’s *d* = 7.7, p = 1×10^-14^).

To better understand this surprising result, we looked at how the SVMs trained on early RS correct ensemble data, specifically classified activity observed on either early RS error (**Figure 3C**) or early RS correct (**Figure 3D**) trials. To evaluate the difference between class predictions of CCG models, we quantified the proportion of test samples classified as “Correct” (i.e., resembling early IA correct). We observed that activity on early RS error trials was mainly classified as Correct in WT VEH data (% classified as Correct: 96.2% +/- 1.1). In contrast, we observed the opposite pattern for Het VEH data, where early RS error trial activity was mainly classified as Error (% classified as Correct: 18.2% +/- 3.0). The % of early RS error trial activity patterns classified as Correct was increased (compared to Het VEH) in the Het CLNZ and Het postCLNZ conditions (Het CLNZ: 44.3% +/- 1.3, Het postCLNZ: 62.8% +/- 2.0). By contrast, classification of early RS correct trial activity within the same ensemble was much less variable between conditions: in both WT VEH and Het VEH, these activity patterns were classified as Correct vs. Error at similar levels (WT VEH: 54.7% +/- 1.5, Het VEH: 53.3% +/- 1.8, Het CLNZ: 65.8% +/- 2.0, Het postCLNZ: 77.2% +/- 1.7). Thus, the Het VEH condition is specifically associated with *abnormally stable* encoding of *error* outcomes across the IA and RS portions of the task, whereas the encoding of correct outcomes is substantially less affected.

This suggests that not only is outcome encoding typically not conserved from IA to RS, but there is a neuronal ensemble in which outcome encoding typically reverses from IA to RS. Error outcomes during the RS evoke activity patterns resembling those which followed correct outcomes during the IA. Furthermore, within this ensemble, encoding of errors becomes abnormally preserved in Het VEH mice, and CLNZ treatment serves to ameliorate this abnormality both acutely and persistently. Thus, the stable encoding of error outcomes appears to be pathological during rule shifts.

### Changes in activity patterns from the IA to RS are diminished in *Dlx5/6^+/-^* mutants

Next, we looked for other examples of cross-condition generalization, or the lack thereof. Specifically, we examined whether there is encoding of the early IA vs. early RS portions of the task that generalizes across correct vs. error trials (**Figure 4A**).

**FIGURE 4:**
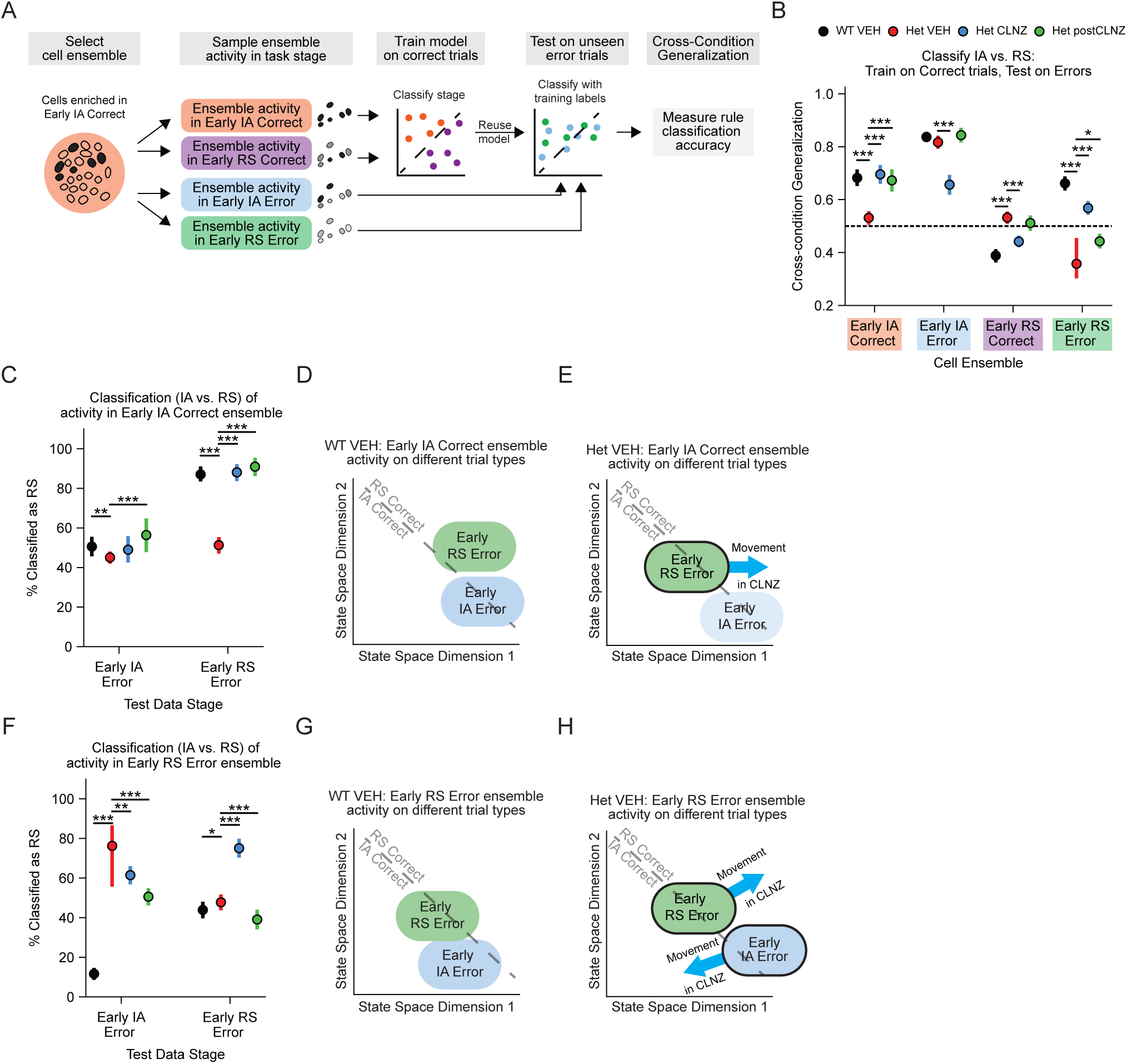
Encoding of the early IA vs. early RS task stages generalizes across correct vs. error outcomes in wild-type but not mutant mice. A. Schematic for Cross-Condition Generalization Performance (CCGP). Neuronal ensemble activity is resampled from early IA Correct, early RS Correct, early IA error, and early RS correct. SVMs are trained to classify that ensemble’s activity on early IA correct vs. early RS correct trials. The trained CCG model is then tested on unseen data: that ensemble’s activity on early IA error and early RS error trials. CCG Performance computes how well the SVM classifies the test data as ‘IA’ or ‘RS’. B. CCGP is plotted for each ensemble (early IA / RS correct / error) and each genotype-treatment condition (different colored filled circles). Generalization of WT VEH encoding of ‘IA’ vs. ‘RS’ is maintained (CCGP > 0.5) for multiple ensembles. In Het VEH, this encoding is lost in the early IA correct and early RS error ensembles, but is improved by CLNZ treatment. C. Percent of samples from early IA error trials or early RS error trials classified as early RS correct by CCG models trained on early IA correct ensemble activity, for each genotype-condition. In WT VEH, activity on early RS error trials is mainly classified as ‘RS’, which reverts to chance levels in the Het VEH condition. CLNZ treatment ameliorates this abnormality. D. Diagram of the inferred geometry of task stage / trial type activity for the early IA correct ensemble in WT VEH, relative to the CCG model classification boundary separating activity of early IA correct vs. early RS correct trials. This schematizes the quantitative changes in encoding shown in panel C. The blue arrow represents the effects of CLNZ treatment. E. Similar to Panel D, but for early IA correct ensemble dynamics in Het VEH mice. F. Similar to panel C, but showing the percent of samples of early RS error ensemble activity from early IA error or early RS error classified as early IA correct. G-H. Similar to panels D and E but schematizing the quantitative changes in early RS error ensemble encoding (shown in panel F) for WT Veh (panel G) and Het VEH (panel H). Panels B, C, & F: p-value calculated from Cohen’s U3, which is derived from each comparison’s Cohen’s *d.* Points and error bars show the mean +/- 75%ile of N = 1000 bootstrap distributions per genotype-treatment ensemble. \**P* < 0.05, \*\**P* < 0.01, \*\*\**P* < 0.001.

When we trained SVMs to classify early IA correct vs. early RS correct trials, then tested them using early IA error vs. early RS error trials, we found that SVMs trained using many different WT VEH ensembles consistently performed this classification at better-than-chance levels (filled black circles in **Figure 4B**). This was true for SVMs based on the early IA correct, early IA error, and early RS error ensembles. Notably, this successful cross-condition generalization of task phase information (IA vs. RS) was diminished in the Het VEH condition for SVMs based on both the early IA correct and early RS error ensembles. In both of these cases, CLNZ treatment improved performance in HET mice acutely as well as persistently (<u>early IA correct ensemble</u> <u>generalization accuracy:</u> WT VEH = 0.68 +/- 0.01, Het VEH = 0.53 +/- 0.01, Het CLNZ = 0.70 +/- 0.02, Het postCLNZ 0.67 +/- 0.02; WT VEH vs Het VEH Cohen’s *d* = 12, p < 10^-15^; Het VEH vs Het CLNZ Cohen’s *d* = 12, p < 10^-15^; Het VEH vs Het postCLNZ Cohen’s *d* = 9.0, p < 10^-15^; <u>early RS error ensemble generalization accuracy</u>: WT VEH = 0.66 +/- 0.01, Het VEH = 0.35 +/- 0.04, Het CLNZ = 0.57 +/- 0.01, Het postCLNZ = 0.44 +/- 0.01; WT VEH vs Het VEH Cohen’s *d* = 12, p < 10^-15^; Het VEH vs Het CLNZ Cohen’s *d* = 8.1, p < 10^-15^; Het VEH vs Het postCLNZ Cohen’s *d* = 3.5, p = 0.00027).

We further investigated this result by examining how SVMs specifically classified activity from early IA error or early RS error trials. In the WT VEH, Het CLNZ, and Het postCLNZ conditions, SVMs based on activity of the early IA correct ensemble consistently classified activity during early RS error trials as ‘RS’ rather than ‘IA,’ i.e., more similar to early RS correct than early IA correct trials <u>(% classified as RS:</u> WT VEH = 86.6% +/- 1.5, Het CLNZ = 88.2% +/- 1.9, Het postCLNZ = 91.2% +/- 2.0 ). By contrast, in the Het VEH condition, this classification was almost evenly split between RS and IA (Het VEH, % classified as RS = 51.4% +/- 1.8) (**Figure 4C**). In contrast to this disruption of IA vs. RS encoding during early RS error trials, the activity classification on early IA error trials was unperturbed. SVMs classified activity on early IA error trials as IA vs. RS similarly (near-even split) across all conditions.

This indicates that neurons that had been strongly recruited in early IA correct trials normally exhibit a similar change in their activity patterns across early RS correct and early RS error trials; this change in activity patterns seems to be lost in HET mice, but is restored by CLNZ treatment. Similar to what we observed in Figure 3, this suggests that during the learning of rules shifts, there is normally a change in activity patterns that supersedes any potential stable encoding of outcomes. This change in activity patterns is particularly prominent (and becomes disrupted in the Het VEH condition) for neurons strongly recruited following early IA correct outcomes.

We also examined SVMs based on activity within the early RS error ensemble (**Figure 4F**). In this case, activity on early IA error trials usually is (in WT mice) classified as early IA correct (<u>% classified as RS</u>: WT VEH: 11.6% +/- 1.3), rather than early RS correct. This flips in the Het VEH condition, such that activity in this ensemble becomes predominately classified as early RS correct, but is significantly attenuated (though not completely reversed) by CLNZ treatment (<u>%</u> <u>classified as RS:</u> Het VEH: 78.3 % +/- 7.8; Het CLNZ: 75.1% +/- 2.1; Cohen’s *d* = 2.9, p = 0.0016). Again, this suggests that typical shifts in activity patterns from the IA to RS portion of the task, become disturbed in the Het VEH condition, although the reversal of this abnormality by CLNZ is less robust than prior examples.

To summarize our findings from CCGP analysis: (1) outcome signals usually are not maintained from the IA to RS; (2) the abnormally stable encoding of error outcomes from IA to RS is associated with perseveration (specifically for neurons that are most strongly recruited on early RS correct trials); (3) there are changes in activity patterns from IA to RS that are shared across correct and error outcomes; (4) these task phase-specific activity patterns are disrupted during perseverative behavior (particularly in neurons that had been strongly recruited on early IA correct trials).

### Activity following IA and RS errors becomes less separable in *Dlx5/6^+/-^* mutants

Next, we used linear classification to investigate our basic finding further, which is that HET mice exhibit abnormal persistence of activity patterns associated with the IA during the RS, particularly for error-related signals. First, we specifically trained SVMs to distinguish activity observed on early IA error vs. early RS error trials within either the early IA error or early RS error ensemble. In this case, the data used for training and testing was drawn from the same underlying distribution used to generate pseudopopulation activity. Thus, this should not be taken as a cross-validated measure of classifier performance on novel data, but rather as a measure of the linear separability of activity during these two types of trials.

Notably, the two ensembles were largely distinct but exhibited some membership overlap (**Figure 5A**). The degree of overlap was not significantly different from chance in WT VEH (p = 0.57, χ2 test), Het CLNZ (p = 0.35, χ2 test), and Het postCLNZ conditions (p = 0.19, χ2 test) but was significantly greater than chance in Het VEH (p = 0.041, χ2 test). No significant difference in the degree of overlap was observed between genotype-treatment groups (**Figure 5B**).

**FIGURE 5:**
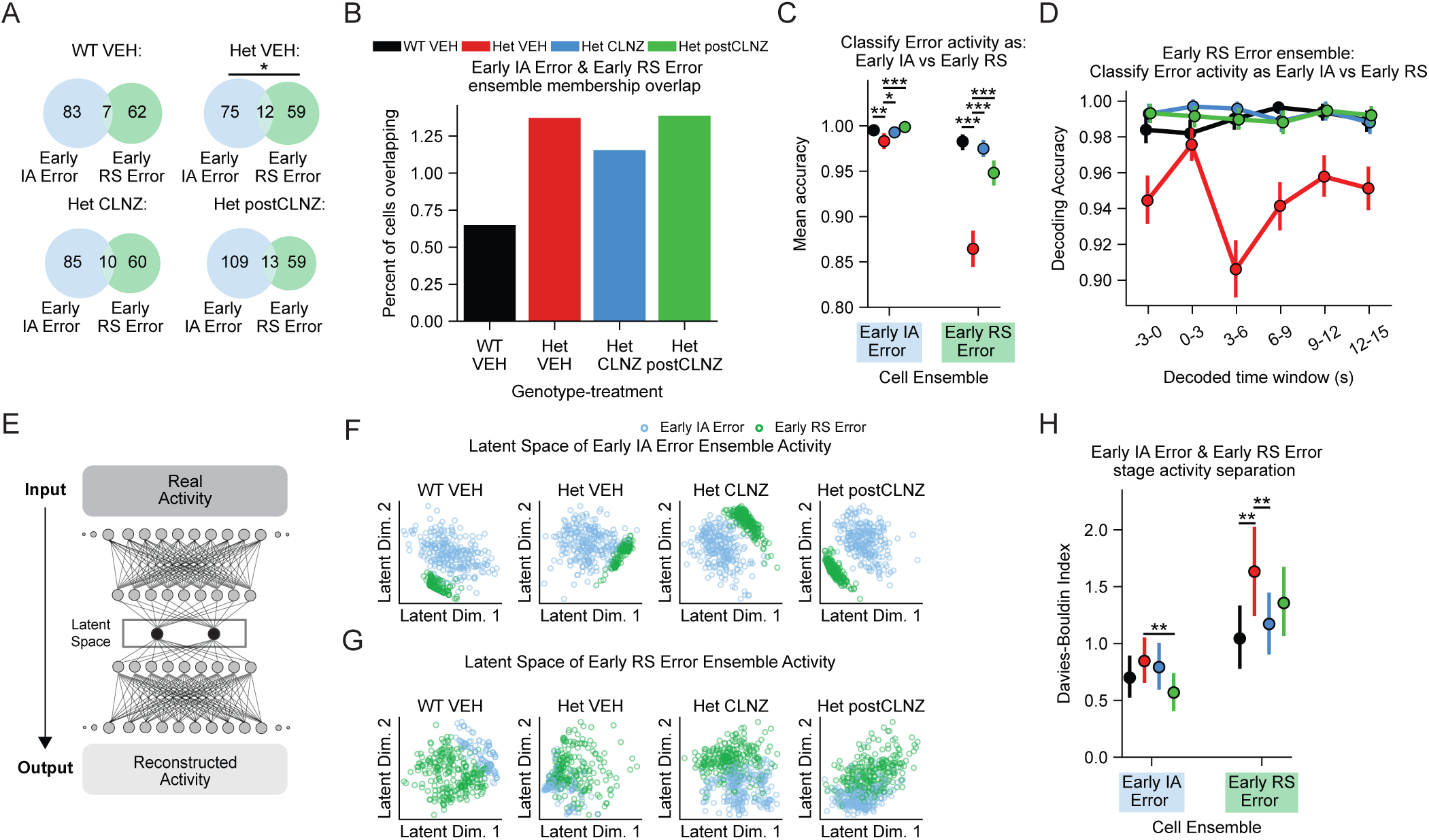
Activity following IA and RS errors becomes less separable in Dlx5/6 mutants. A. The number of cells overlapping between the early IA error and early RS error ensembles is only greater than chance in Het VEH mice. B. The % of all cells that overlap between the early IA error and early RS error ensembles is not significantly different by genotype-treatment group. C. Linear classifiers (SVM) trained to distinguish early IA error vs. early RS error trial activity from the early IA error or early RS error ensembles have significantly worse accuracy in both ensembles of Het VEH mice, which is lastingly improved by CLNZ treatment. D. Similar to panel C, but shows classifier performance dependent on time elapsed since trial outcome. Performance is most impaired 3-6 seconds post-outcome. Points and error bars show the mean +/- 75%ile of N = 1000 bootstrap distributions per genotype-treatment ensemble. E. Schematic of the neural network autoencoder. Networks are trained to compress data into a low-rank bottleneck latent space, and then to reconstruct data from this latent representation. F. Representative autoencoder latent space projections of early IA error ensemble activity from early RS error and early IA error trials. G. Same as E, but plotting the latent space of early RS error ensemble data. H. Quantifying cluster separation of early RS error and early IA error activity in the latent space of autoencoders trained on data from early IA error then early RS error ensembles. The Davies-Bouldin index of activity in the early RS error ensemble’s latent space is significantly higher in Het VEH vs WT VEH, indicating high cluster mixing/low separation. This lastingly decreases as a result of CLNZ treatment. Panels A, B: p-value derived from chi-squared test on ensemble member counts. Panels C, H: p-value calculated from Cohen’s U3, which is derived from each comparison’s Cohen’s *d.* Points and error bars show the mean +/- 75%ile of N = 1000 bootstrap distributions per genotype-treatment ensemble. \**P* < 0.05, \*\**P* < 0.01, \*\*\**P* < 0.001.

The classifier accuracy across all genotypes and conditions was near 100% when decoding trial type (early IA vs. RS error) from activity in the early IA error ensemble (**Figure 5C**). However, when decoding trial type using activity within the early RS error ensemble, performance for Het VEH was significantly lower than all other conditions (<u>SVM accuracy</u>: WT VEH = 0.98 +/- 0.004, Het VEH = 0.87 +/- 0.01, Het CLNZ = 0.98 +/- 0.004, Het postCLNZ = 0.95 +/- 0.006; WT VEH vs Het VEH Cohen’s *d* = 16, p < 10^-15^; Het VEH vs Het CLNZ Cohen’s *d* = 15,p < 10^-15^; Het VEH vs Het postCLNZ Cohen’s *d* = 10, p < 10^-15^).

To further elucidate the nature of this abnormality, we trained and tested SVMs using activity from more restricted time windows in the early RS ensemble (**Figure 5D**). Classifier performance was lowest in Het VEH mice 3-6 seconds following the outcome and remained low thereafter. This suggests that during the outcome period, when mice receive feedback that they have made an incorrect choice, activity on early RS error trials becomes less distinguishable from activity previously observed on early IA error trials in the Het VEH condition.

We then used deep feed-forward neural network autoencoders to produce low-dimensional nonlinear representations of ensemble activity in task stages. Autoencoder networks contain a low-rank hidden layer acting as a bottleneck. They are trained to reconstruct input data based on the compressed representations within this low-dimensional bottleneck, without regard to input class (**Figure 5E**). This makes it possible to visualize and analyze neural activity by examining compressed representations within the latent space of the trained autoencoder network.

SVM accuracy measures the maximum separability between activity patterns from different trial types that can be achieved using supervised learning. By contrast, the distance between representations in the autoencoder latent space reflects the separability that emerges when a neural network learns the structure of the data in an unsupervised manner, i.e., without knowing which activity patterns derive from specific trial types. Thus, these latent space representations do not suffer from the caveats noted above for SVMs, and may better capture the biological functions of mPFC cell ensembles.

We trained a neural network autoencoder to reconstruct the activity of neurons in the early IA error ensemble on early IA error and early RS error trials (**Figure 5F**). We then plotted activity from these two trial types in the autoencoder latent space, and quantified the separation between these trial types using the Davis-Bouldin index (DB Index), which quantifies the cohesion and separation of clusters of points (here, each cluster corresponds to one trial type). Specifically, the DB index is the sum of the distances between points corresponding to the same trial type, relative to the distance between the centroids of these two clusters. Lower values of the DB index indicate greater separation between clusters and decreased spread (increased cohesion) of points within each cluster.

We projected 200 random samples of activity (within the early IA error ensemble) from each trial type into the latent space of autoencoders, which had been trained (using 500 samples from each trial type) to reconstruct the input, without regard to trial type. We then calculated the DB index 500 times, each based on 200 random activity samples from each trial type. The DB index for early IA error ensemble activity was not significantly different in Het VEH compared to WT VEH (**Figure 5F, H**). Next, we applied this autoencoder analysis to the early RS error ensemble activity during early IA error vs. early RS error trials (**Figure 5G, H**). In this case, the DB index was significantly higher for Het VEH than WT VEH and Het CLNZ, but this difference was not significant for the Het postCLNZ conditions (**Figure 5H**) (<u>early RS error ensemble DB Index</u>: WT VEH = 1.0 +/- 0.1, Het VEH = 1.6 +/- 0.2, Het CLNZ = 1.2 +/- 0.1, Het postCLNZ = 1.3 +/- 0.2. WT VEH vs Het VEH Cohen’s *d* = 3.1, p = 0.001; Het VEH vs Het CLNZ Cohen’s *d* = 2.4, p = 0.009; Het VEH vs Het postCLNZ Cohen’s *d* = 1.3, p = 0.097).

Examining the distribution of points corresponding to the projections of activity patterns into the autoencoder latent space shows that not only are points from these two clusters more overlapping in all of the HET conditions, but the cluster of points corresponding to activity on early RS error trials also becomes much less cohesive in the Het VEH condition.

These results broadly align with our previous findings that error encoding becomes abnormally stable in the Het VEH condition. However, whereas our CCGP analysis found abnormally stable error encoding in the *RS correct* ensemble, the current analysis specifically examined error encoding by the *early IA error* and *early RS error* ensembles. Furthermore, quantifying SVM accuracy reveals decreased separability between activity on early RS error and early IA error trials within the early RS error ensemble. By quantifying distances within a non-linear autoencoder latent space, we find decreased separability of these two trial types for the early RS error ensemble. These two metrics – SVM accuracy and latent space distance – quantify separability differently (as discussed further in the Discussion). However, both reveal abnormal activity structure within cells enriched during early RS error trials.

### Activity on early IA and RS correct trials becomes less separable in *Dlx5/6^+/-^* mutants

Next, we applied the same analyses to compare activity patterns during early IA and RS correct trials. In this case, the ensembles that were significantly recruited on early IA correct and early RS correct trials (**Figure 6B**) were consistently more overlapping than expected by chance across all conditions (p = 0.0011, 1.8 x 10^-15^, 0.028, 2.7 x 10^-6^ in WT VEH, Het VEH, Het CLNZ, and HET post-CLNZ, respectively, χ2 test). However, the degree of overlap was significantly increased in Het VEH compared to both WT VEH (p = 0.0011, χ2 test) and Het CLNZ (p = 0.0045, χ2 test); for Het VEH vs. HET post-CLNZ, there was a non-significant trend towards higher overlap (p = 0.13; χ2 test). Despite this high overlap between ensembles, the SVM accuracy was consistently close to 1 for all genotype-treatment conditions when we trained linear classifiers to distinguish these two trial types based on activity within the early IA correct ensemble (**Figure 6C**). For the early RS correct ensemble, SVM accuracy was also high in the WT VEH condition, but was significantly lower in Het VEH mutant mice. CLNZ treatment led to both an acute and persistent rescue of decoding accuracy using the RS correct ensemble (<u>early</u> <u>RS correct ensemble mean accuracy</u>: WT VEH = 0.98 +/- 0.004, Het VEH = 0.92 +/- 0.008, Het CLNZ = 0.98 +/- 0.005, Het postCLNZ 0.98 +/- 0.004; WT VEH vs Het VEH Cohen’s *d* = 8.9, p < 10^-15^; Het VEH vs Het CLNZ Cohen’s *d* = 8.7, p < 10^-15^; Het VEH vs Het postCLNZ Cohen’s *d* = 9.4, p <10^-15^).

**FIGURE 6:**
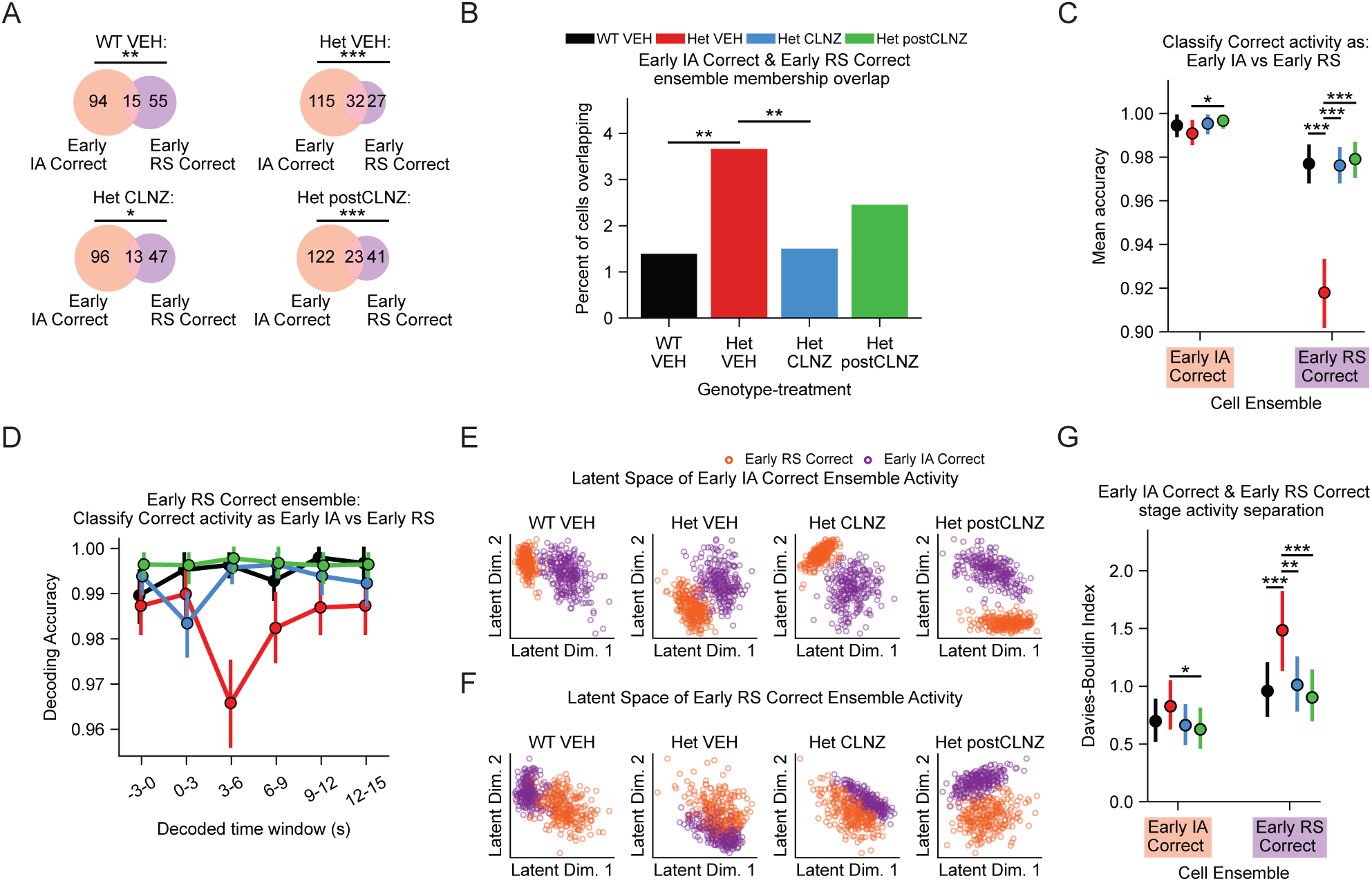
Activity on early IA and RS correct trials becomes less separable in *Dlx5/6* mutants. A. The number of cells overlapping between the early IA correct and early RS correct ensembles is significantly greater than chance in all genotype-treatment conditions. B. The % of all recorded cells that overlap between the early IA correct and early RS correct ensembles is significantly higher in Het VEH versus WT VEH, and is acutely decreased in the Het CLNZ condition. C. Linear classifiers (SVM) trained to distinguish the activity of either the early IA correct or early RS correct ensembles on early IA correct vs. early RS correct trials. Classification is significantly worse in Het VEH mice using the early RS correct ensemble, and is lastingly improved by CLNZ treatment. D. Similar to panel C, but showing classifier performance by time window relative to the bowl dig triggering the trial outcome. Performance is most impaired 3-6 seconds post-outcome. Points and error bars show the mean +/- 75%ile of N = 1000 bootstrap distributions per genotype-treatment ensemble. E. Representative latent space projections of activity from early IA correct and early RS correct trials using the early IA correct ensembles. F. Same as E, but plotting the latent space of early RS correct ensemble data. G. Using the Davies-Bouldin Index to quantify cluster separation of early IA correct or early RS correct activity in the latent space of autoencoders trained on data from early IA correct then early RS correct ensembles. DB index of activity in latent space of early RS correct ensemble data is significantly higher in Het VEH vs WT VEH, indicating high cluster mixing/low separation. This lastingly decreases as a result of CLNZ treatment Panels A, B: p-value derived from chi-squared test on ensemble member counts. Panels C, G: p-value calculated from Cohen’s U3, which is derived from each comparison’s Cohen’s *d.* Points and error bars show the mean +/- 75%ile of N = 1000 bootstrap distributions per genotype-treatment ensemble. \**P* < 0.05, \*\**P* < 0.01, \*\*\**P* < 0.001.

Similar to how we previously used autoencoders, we now projected activity observed on early IA correct and early RS correct trials onto the two-dimensional latent space of a neural network autoencoder (**Figure 6D**). We trained separate autoencoder networks for the early IA and RS correct ensembles. Clouds of points corresponding to activity patterns of the early IA correct ensemble during either early IA correct or early RS correct trials were visually separated for all genotype-conditions. The DB index for this ensemble’s activity was not significantly different between Het VEH and either WT VEH or Het CLNZ, only showing a small but significant difference between Het VEH and Het postCLNZ (<u>early IA correct ensemble DB index:</u> Het VEH = 0.81 +/- 0.11, Het postCLNZ = 0.61 +/- 0.09; Het VEH vs Het postCLNZ Cohen’s *d* = 1.9, p = 0.026). However, within the latent space of early RS correct ensembles, activity during early IA correct and early RS correct trials became highly overlapping in the Het VEH condition, mirroring our SVM results. The DB index was significantly higher (indicating reduced separability / decreased cluster cohesion) for samples from early RS correct ensemble for Het VEH compared to all other genotype-treatment groups (**Figure 6G**; <u>early RS correct ensemble DB index</u>: WT VEH = 0.94 +/- 0.12, Het VEH 1.42 +/- 0.18, Het CLNZ = 0.99 +/- 0.12, Het postCLNZ = 0.89 +/- 0.11; WT VEH vs Het VEH Cohen’s *d* = 3.1, p = 0.0009; Het VEH vs Het CLNZ Cohen’s *d* = 2.8, p = 0.0025; Het VEH vs Het postCLNZ Cohen’s *d* = 3.5, p = 0.0002).

Thus, results from ensemble overlap, SVM accuracy, and autoencoder latent space distance all consistently indicate that abnormalities are most prominent in cells enriched during early *RS correct trials*. For this ensemble, activity patterns in *Dlx5/6^+/-^* mutants on early RS correct trials become abnormally similar to those previously observed during early IA correct trials. This abnormality is reversed by CLNZ.

### Abnormally similar correct trial activity is reinstated during the RS

The preceding results raise a question: is the abnormal similarity of early IA correct and early RS correct activity patterns in Het VEH (and the abnormal overlap between the early IA correct and early RS correct ensembles) simply a reflection of activity patterns that emerge after early IA correct trials and *persist* across the late IA period, into the early RS? Or is abnormally similar activity inappropriately *reinstated* during the early RS?

To distinguish between these two possibilities, we began by examining the relationship between activity occurring during early IA correct trials and late IA trials. First, we quantified the overlap between the early IA correct and late IA ensembles. Similar to what we had seen for early IA correct and early RS correct ensembles, early IA correct and late IA ensembles were more overlapping than expected by chance in all conditions (p = 0.0001 in WT VEH, 9 x 10^-16^ in Het VEH, 4 x 10^-7^ in Het CLNZ, and 2 x 10^-7^ in HET post-CLNZ, χ^2^ test) (**Figure 7A**). Furthermore, the degree of overlap was significantly greater in Het VEH compared to all other conditions (Het VEH vs. WT HET: p = 0.0001, Het VEH vs. Het CLNZ: p = 0.004, Het VEH vs. HET post-CLNZ: p = 0.032, χ2 test) (**Figure 7B**).

**FIGURE 7:**
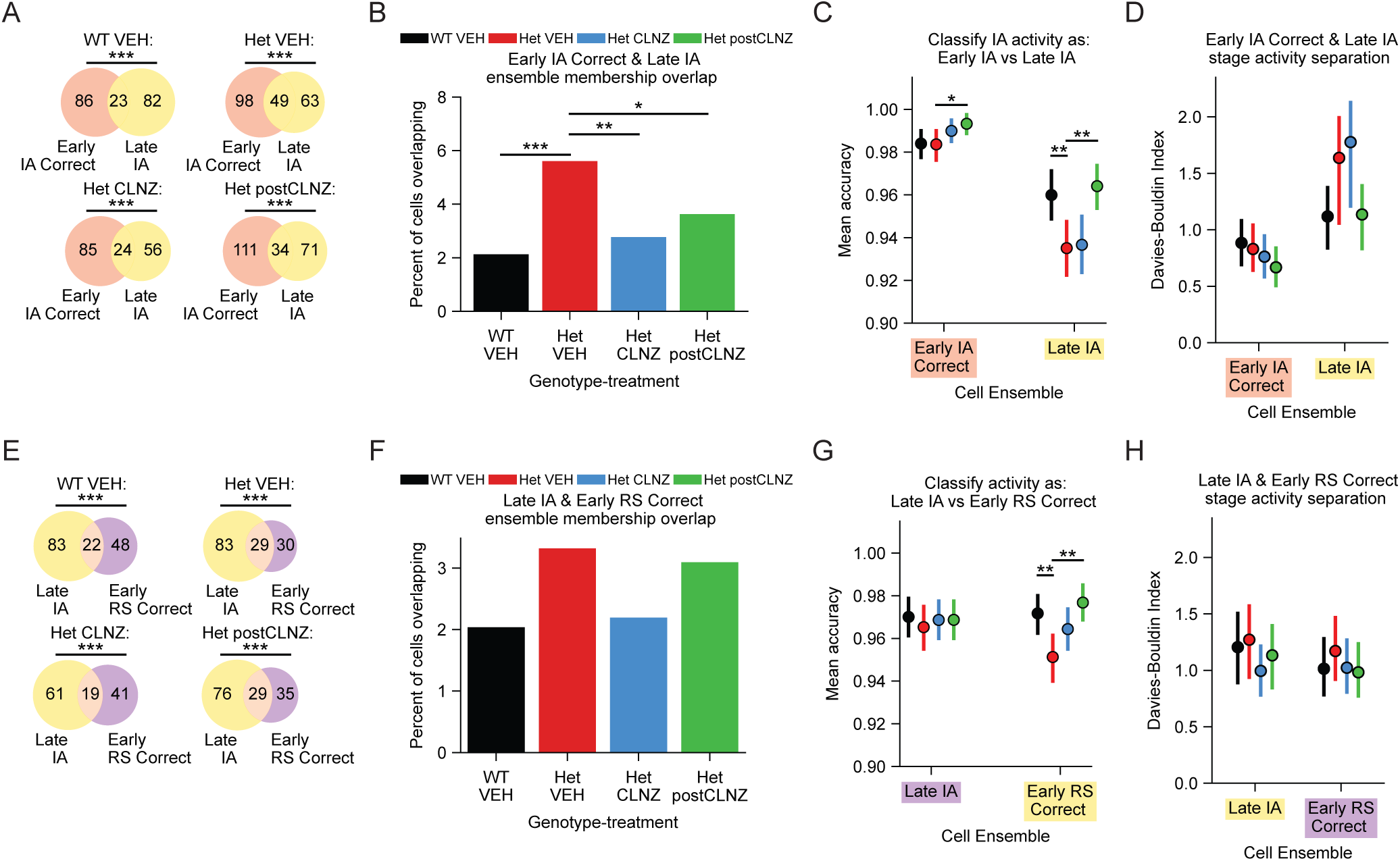
Abnormally similar correct trial activity in *Dlx5/6*^+/-^ mice persists from the early to late IA, but not from the late IA to early RS period. A. The number of cells overlapping between the early IA correct and late IA ensembles is significantly greater than chance in all genotype-treatment conditions. B. The % of recorded cells overlapping between the early IA correct and late IA ensembles is significantly greater in Het VEH versus all other conditions. C. Linear models (SVM) classify whether the activity of either the early IA correct or late IA ensembles originate from early IA correct vs. late IA trials. Classification accuracy is not significantly different between late IA ensembles of Het VEH and WT VEH. Accuracy significantly decreases in late IA ensemble of Het VEH compared to WT VEH, but is not acutely altered by CLNZ treatment. D. Using the Davies-Bouldin Index to quantify cluster separation of early IA correct or late IA stage activity in the latent space of autoencoders trained on data from early IA correct and late IA ensembles. Data are mean +/- 75%ile of 1000 bootstrap distributions. P-value derived from Cohen’s U3. \**P* < 0.05, \*\**P* < 0.01, \*\*\**P* < 0.001. E-H: Similar to A-D but comparing the late IA and early RS correct trial stages. Panels A, B, E, F: p-value derived from chi-squared test on ensemble member counts. Panels C, D, G, H: p-value calculated from Cohen’s U3, which is derived from each comparison’s Cohen’s *d.* Points and error bars show the mean +/- 75%ile of N = 1000 bootstrap distributions per genotype-treatment ensemble. \**P* < 0.05, \*\**P* < 0.01, \*\*\**P* < 0.001.

Next, we measured how well activity from these two trial types could be separated using linear SVMs. We observed a slight decrease in decoding accuracy when using the late IA ensemble in Het VEH mice relative to WT VEH – this was reversed in the Het postCLNZ condition but not in the Het CLNZ condition (**Figure 7C**). In the latent space of autoencoders, we observed no significant difference in the separability of early IA correct vs. late IA trial activity for the early IA correct ensemble (**Figure 7D**). No differences in the separability of stage activity were observed for late IA ensembles either. Thus, ensemble overlap and SVM accuracy results demonstrate abnormally increased similarity of activity patterns between early IA correct and late IA trials, specifically in the Het VEH condition. However, no significant abnormalities were observed in the nonlinear autoencoder latent space.

Next, we tested whether abnormal similarity persisted between late IA and early RS correct trials. Task stage ensemble membership was more overlapping than expected by chance for all genotype-treatment groups (p = 2×10^-10^ in WT VEH, 5×10^-18^ in Het VEH, 5×10^-10^ in Het CLNZ, and 3×10^-19^ in HET post-CLNZ, χ^2^ test) (**Figure 7E**). However, we did not observe any significant differences in overlap between untreated mutant mice and wildtype mice, nor did we observe significant changes in overlap during or after CLNZ treatment in mutant mice (Het VEH vs. WT HET: p = 0.08. Het VEH vs. Het CLNZ: p = 0.15. Het VEH vs. HET post-CLNZ: p= 0.79, χ^2^ test; **Figure 7F**).

When we measured the accuracy of SVMs for decoding late IA vs. early RS correct trials using early RS correct ensemble data, we found a slight but significant decrease in Het VEH relative to the WT VEH, Het CLNZ and Het postCLNZ conditions (**Figure 7G**; <u>SVM accuracy</u>: WT VEH = 0.972 +/- 0.005, Het VEH = 0.952 +/- 0.006, Het CLNZ = 0.965 +/- 0.005, Het postCLNZ = 0.977 +/- 0.004; WT VEH vs Het VEH Cohen’s *d* = 3.7, p = 0.00011, Het VEH vs Het CLNZ Cohen’s *d* = 2.3, p = 0.01, Het VEH vs Het postCLNZ Cohen’s *d* = 5.0, p = 4×10^-7^). However, latent space projections of activity from either late IA ensemble or early RS correct ensembles again revealed no significant abnormalities in stage activity separation for mutant mice (**Figure 7H**).

In summary, in the Het VEH condition, there is evidence for increased persistence of activity patterns from early IA correct to late IA trials. However, if these activity patterns persisted from late IA trials into early RS correct trials, we would expect to observe reduced separability of activity patterns from late IA and early RS correct trials. In fact, in the Het VEH condition, activity patterns on late IA and early RS correct trials remained separated in the autoencoder latent space. This suggests that in mutants, the abnormal similarity between activity patterns on early RS correct trials and those which previously occurred during early IA correct trials reflects an active re-instantiation of these early-stage activity patterns, not just their passive persistence over time.

### Effects of CLNZ in WT mice

To assess whether the effects of CLNZ in Het mice reflect therapeutic effects versus nonspecific perturbations, we also examined neural activity in wild-type mice treated with CLNZ. We did not observe effects of CLNZ on WT task performance in IA or RS (**Figure 1I, J**) (<u>IA trials to</u> <u>complete:</u> WT VEH 14.1 +/- 0.7 vs WT CLNZ 13.5 +/- 0.7, p = 0.45; RS: WT VEH 15.5 +/- 1.7 vs. WT CLNZ 16.3 +/- 0.9, p = 0.19, MWU, n = 7 mice). We observed a slight but significant increase in perseverative errors made during the RS for WT CLNZ vs WT VEH (**Figure 1K**; WT VEH 3.0 +/- 1.1 vs. WT CLNZ 4.5 +/- 0.5, p = 0.047, MWU, n = 7 mice).

We observed minor effects of CLNZ on overall cell activity levels (**Supplementary Figure 1C, D**). In WT mice, CLNZ did not affect the proportions of cells enriched that were active in specific task stages (**Supplementary Figure 1J**) or assigned to the corresponding task stage ensembles (**Supplementary Figure 1G, H**). We also did not observe significant differences in cell ensemble overlap between WT VEH and CLNZ (**Supplementary Figure 2B**). Similar to WT mice, we did not observe significant overlap between the early IA error and early RS error ensembles, or between the early IA correct and early RS correct ensembles. However, as was the case for all other conditions (**Figure 7**), in WT CLNZ there was significant overlap between the early IA correct and late IA ensembles, as well as between the late IA and early RS correct ensembles (**Supplementary Figure 2A**; early IA correct and late IA p = 0.006, late IA and early RS correct p = 0.002, χ^2^ test).

Next, we examined several of the metrics that were abnormal in Het mice (relative to WT) and normalized by CLNZ. First, we examined results from our Cross-Condition Generalization (CCG) analysis. When examining classification generalization as ‘Correct’ vs. ‘Error’, we observed that for Het VEH, activity of the early RS correct ensemble on early RS error trials was classified mainly as Error. By contrast, this was classified mainly as Correct in WT VEH, and evenly split in HET mice during and after CLNZ treatment (**Figure 3**). Similar to what we observed in the Het CLNZ condition, in WT CLNZ datasets, classification of early RS correct ensemble activity on early RS error trials was nearly equally split between Correct and Error (**Supplementary Figure 2C**).

Similarly, we found abnormal generalization of ‘IA’ vs. ‘RS’ classification in Het VEH for two cases: early IA correct ensemble activity on early RS error trials and early RS error ensemble activity on early IA error trials. In both cases, abnormalities in HET mice were ameliorated by CLNZ, and the classification pattern in WT CLNZ was similar to that observed in Het CLNZ **(Supplementary Figure 2D-E**).

Notably, there was one case in which the effects of CLNZ in WT mice deviated from the pattern noted above and seemed to recapitulate the abnormality observed in the Het VEH condition. This is for the analyses shown in Figure 4 – examining the separation between activity of the early RS error ensemble on early IA error and early RS error trials using SVMs or autoencoders. For both of these, we saw that the decreased separability observed in Het VEH also occurred in WT CLNZ, suggesting that this particular abnormality may not be associated with impaired flexibility **(Supplementary Figure 2F-G)**. By contrast, the decreased separation between activity of the early RS correct ensemble on early IA correct and early RS correct trials we observed in Het VEH mice did not occur in the WT CLNZ condition (which was very similar to WT VEH for these metrics; **Supplementary Figure 2H-I**).

## DISCUSSION

Encoding information via high-dimensional activity within prefrontal ensembles facilitates behavioral adaptation in dynamic environments (Tye et al., 2024; Durstewitz et al., 2010; Rikhye et al., 2018; Bernardi et al., 2020; Rigotti et al., 2013). mPFC cell ensembles represent concrete information (e.g., encoding trial outcomes) and abstract variables (e.g., current rule, level of uncertainty, etc.) as mice perform behavioral tasks (Spellman et al., 2021; Rikhye et al., 2018; Jun et al., 2024; Bernardi et al., 2020). Parvalbumin-positive inhibitory interneurons regulate mPFC microcircuit activity and support efficient cognitive flexibility, but their precise role in normal mPFC encoding remains unclear (Ferguson & Gao, 2018). In this study, we characterize neural activity during a rule-shifting task, evaluating normal patterns of information encoding during cognitive flexibility and how these patterns are perturbed and selectively restored in mutant mice with PVIN dysfunction but reversible cognitive impairments. Our results shed light on the specific aspects of prefrontal encoding that are necessary for successfully learning extradimensional rule changes and regulated by inhibitory circuits.

Our previous studies have shown that prefrontal PVINs normally increase their activity and gamma synchrony after errors during rule shift learning, and that both of these signals are lost in *Dlx5/6^+/-^* mutant mice. Based on this, a reasonable model would be that: 1) increases in PVIN activity and gamma synchrony help create prefrontal error signals; 2) the robust signaling of error outcomes by prefrontal cortex is critical for learning extradimensional rule shifts; and 3) weak or unstable error signals, resulting from PVIN dysfunction, are a primary contributor to perseveration in *Dlx5/6^+/-^*mutants. This hypothesis would be consistent with findings that prefrontal projection neurons mainly encode trial feedback information during extradimensional rule changes in a similar cognitive flexibility task (Spellman et al., 2021). However, our findings suggest a different model.

In particular, we find that the population-level encoding of trial outcomes by prefrontal ensembles is normally extremely dynamic, such that rule shift *errors* evoke patterns of activity closely resembling those which previously followed *correct* outcomes during learning of the initial rule. It is unexpected and surprising that more stable outcome encoding is associated with perseveration in *Dlx5/6^+/-^*mutants; as noted above, a prevailing hypothesis has been that high-fidelity outcome encoding facilitates flexible behavior, specifically the learning of new rules based on a set of cues that were previously irrelevant to trial outcomes, i.e., an extradimensional shift. Our findings suggest that it is important for trial outcome encoding to be dynamic, particularly on error trials.

We also find activity patterns which seem to reflect the stage of learning (IA vs. RS) rather than trial outcome (correct vs. error). Activity patterns which are shared across correct and error trials during the early RS could signal a state of uncertainty and promote attention to previously-irrelevant cues and/or exploratory behaviors through actions on downstream structures. Alternatively, they might represent nascent representations for candidate strategies, e.g., an emerging representation for the RS strategy.

Both of these phenomena – dynamic encoding of errors and outcome-independent encoding of RS learning -- are disrupted in *Dlx5/6^+/-^* mutants. We observed activity patterns, originally occurring after correct outcomes while learning the IA, being abnormally reinstated (not just passively maintained) following correct outcomes during RS learning. These results imply that perseveration represents an active process whereby outdated activity patterns are inappropriately recalled, not just a passive failure to dislodge pre-existing activity patterns.

### Why focus on activity within specific ensembles?

In recent years, systems neuroscience has increasingly emphasized examining population-level neural encoding, rather than simply quantifying the encoding properties of individual neurons. However, combining the activity of diverse neurons into a single population-level readout also runs risks. For example, neurons are believed to be organized selectively into ensembles that implement specific computations. Similarly, downstream neurons typically receive input from specific subsets of a larger input population. Thus, examining activity at the population level has a potential blind spot: it could miss alterations in encoding that selectively occur within specific ensembles (but not the population at large), even though those selective ensembles may be critical for specific computations or transmitting information to specific downstream targets. For this reason, we examined encoding by specific ensembles. As visualized by the t-SNE plots in Fig. 2I, most neurons form clusters corresponding to elevated activity during a particular trial type / task stage (as opposed to being evenly distributed across a continuum characterized by intermediate levels of recruitment in multiple trial types / task stages). Because of this, we defined ensembles by identifying neurons that significantly increased their activity during a specific trial type / task stage. Neurons with similar recruitment patterns might plausibly represent interconnected ensembles, groups of neurons receiving similar inputs, or neurons which form similar downstream connectivity patterns due to Hebbian plasticity.

Interestingly, by defining ensembles in this manner, we found that individual behavioral variables were often specifically encoded by particular ensembles, and that abnormal encoding in *Dlx5/6^+/-^* mutants manifested in an ensemble-specific manner. Thus, examining task stage-defined ensembles provides a valuable lens for resolving population-level activity. Notably, information related to a particular task stage / trial type was not exclusively present in an ensemble defined by that task stage. This reflects two factors: first, even though ∼60% of neurons that significantly increased activity during any task stage did so exclusively during one stage, the remainder significantly elevated their activity during multiple stages. Second, the ensemble associated with a given task stage comprised between 60-80% of the cells active during that stage, which means that a significant minority of active cells come from other ensembles.

Task stage ensembles are grossly intact in *Dlx5/6^+/-^* mutants, but how each encodes other trial types is altered. Some of these alterations can even be visualized in the t-SNE plots in Figure 2i. While absolute distances between points in t-SNE are not meaningful, the t-SNE algorithm preserves local relationships, i.e., which points are neighbors vs. more distant in the original state space. In this context, it is notable that clusters corresponding to neurons with elevated activity on early IA correct trials and those with elevated activity on early IA error trials are adjacent to each other in t-SNE plots derived from the WT VEH, Het CLNZ, and Het postCLNZ conditions. By contrast, in the Het VEH condition, points corresponding to neurons with elevated activity on early RS correct trials become interspersed between these two clusters.

This aligns with our finding of multiple abnormalities involving the early RS correct ensemble in the Het VEH condition: this ensemble becomes abnormally overlapping with the early IA correct ensemble (Fig. 6A-B), activity in this ensemble because less well-separated on early RS correct and early IA correct trials (Fig. 6G), and error encoding within this ensemble becomes abnormally stable (Fig. 3).

### Why did we quantify particular metrics?

Linear SVMs are commonly employed to analyze population-level neural encoding by identifying patterns of neural activity associated with different behavioral conditions. Cross-condition generalization is a particular application of this approach that has been successfully used to examine neural encoding geometry (Bernardi et al., 2020; Boyle et al., 2024), by determining whether a linear SVM trained to classify a behavioral variable (e.g., correct vs. error outcomes) based on data from one condition (e.g., the IA) performs above chance when performing the same classification in another condition (e.g., the RS). This provides information about whether the network represents specific task variables abstractly, i.e., in a way not confounded by a different task variable. The activity of neurons in a specific ensemble (e.g., the early RS correct ensemble) defines a manifold, and the directions along that manifold separating early IA error and correct trials may or may not be the same as those which separate early RS error and correct trials.

Neural network autoencoders using linear activation functions and mean squared error loss can learn subspaces equivalent to those identified by PCA (Chen & Guo, 2023). By incorporating nonlinear activation functions in neural network autoencoders, we may capture some additional aspects of how cortical networks, which are recurrent and nonlinear, process activity patterns. Autoencoders are also trained in an unsupervised manner without respect to the labels of input points. Thus, whereas SVMs identify the linear subspace that best separates two sets of activity patterns, autoencoders identify the subspace that best explains the variability of activity patterns, then quantify how well that subspace separates two sets of activity patterns. This provides information about whether encoding of a specific task variable occurs in the same dimensions as overall variation in activity patterns (on specific trial types). Furthermore, while linear SVM-based classifiers with knowledge of ground truth may easily learn to separate activity patterns from two trial types, mice must differentiate trial types (e.g., early IA error vs. early RS error) without this ground truth knowledge. Points within the autoencoder latent space are unlabeled, so the separability of points in this latent space (quantified using the DB index) may be better than SVMs for capturing how well the brain can use neural activity to perform this separation, in the absence of explicit knowledge that the rule has changed.

### Relationship to previous work

We previously found that inhibiting callosal fibers from prefrontal PV neurons impairs interhemispheric gamma synchrony and causes activity patterns following early RS errors to become more similar to those previously occurring following early IA errors (Cho et al., 2023). This is consistent with our finding here that in *Dlx5/6^+/-^* mutants (which also have impaired gamma synchrony), activity patterns on early RS error and early IA error trials become more similar / less separable (Fig. 3). Overall, our results support previous findings that manipulations which enhance gamma synchrony in *Dlx5/6^+/-^* mutants lead to a persistent rescue of rule shift learning, by identifying enduring changes in activity patterns which accompany this rescue. While not all aspects of neural activity are persistently normalized, it is remarkable that many of the acute effects of CLNZ on neural activity and encoding outlast the period of CLNZ administration by multiple weeks.

A previous study also used linear decoders to analyze pseudopopulation activity during a rule shifting task similar to this one (Benoit et al., 2022). That study recorded prefrontal spiking activity and found that adolescent inhibition of the mediodorsal thalamus caused impaired decoding of correct vs. error outcomes, which could be rescued by stimulating the thalamus using step-function opsins. Here, we examined outcome encoding using a different approach, quantifying generalization from the IA to RS, rather than focusing on outcome encoding specifically during the rule shift.

Our interpretation – that normal rule shifting involves dynamic outcome encoding and that more stable outcome encoding is associated with perseveration – differs from the conclusions of an earlier study, which performed calcium imaging during a similar task but found outcome encoding that was stable within neurons even over days (Spellman et al., 2021). While both studies examined how mice learn extra-dimensional rule changes, the task design and training paradigm was quite different. Here, we studied freely moving mice digging in bowls to find hidden food rewards. This was preceded by a single day of habituation consisting of 10 trials.

By contrast, Spellman et al. studied head-fixed mice receiving water rewards. These mice underwent multiple sessions of behavioral training prior to performing the task, which consisted of multiple stages prior to the extradimensional shift. Finally, the number of trials within each stage was substantially larger than the number of trials in the initial association in our rule-shifting task. Thus, in Spellman et al., mice possess substantially greater familiarity with trial mechanics (including reward delivery) by the time they encounter a rule shift. This contrasts with our experimental paradigm, where task aspects remain relatively novel during rule shift learning. This likely explains why outcome encoding was dynamic in our study but stable in Spellman et al. Indeed, Spellman et al. observed that the intersession correlation of outcome encoding was relatively low during initial learning stages but much higher by the time of the extra-dimensional shift, consistent with this possibility.

### Caveats and limitations

Due to our nonspecific viral targeting strategy, it remains unclear whether certain classes of projection neurons and/or interneurons preferentially encode subsets of task information (e.g., outcome versus rule period). Various studies have demonstrated cell-type specificity in encoding. For example, during the rule-shift task, we found that PV+ cells increase their activity specifically after RS errors (not during RS correct trials) (Cho et al., 2020). Another study using a different task (also in mice) found that regular spiking cells encode trial-specific information about sensory cues, whereas fast spiking cells encode abstract environmental information about the current rule (Rikhye et al., 2018). Conversely, the study described above examined extra-dimensional shifts using water rewards in head-fixed mice and found similar outcome encoding in thalamic and striatally-projecting prefrontal neurons (Spellman et al., 2021). Thus, examining encoding within specific cell types, e.g., classes of interneurons, neurons which project to specific targets, etc., is a logical next step.

Mice rapidly learn cue-reward association in our version of the task, limiting the number of trials available to analyze for each task stage. Furthermore, the exact number of such trials varies between mice. This motivated our analytical approach, which examines activities in pseudopopulations using bootstrap resampling. One limitation of this method is that while we can examine activity on a specific *type* of trial (e.g., early RS errors), we cannot focus on an individual trial or compare one trial to the preceding / following trials in order to identify changes in neural activity that accompany changes in behavior on a trial-by-trial basis.

As a starting point to address this, we have analyzed activity patterns during the period preceding and immediately following the choice to dig in a specific bowl. Repeating some analyses using shorter time windows (c.f., Fig. 5d) shows that abnormalities in Het VEH mice are most prominent a few seconds after the beginning of the dig, at which point the mouse is likely to be processing the presence or absence of reward. That said, because we analyzed activity over extended periods (∼10-15 second windows), we ignored potential time-dependent neural dynamics, e.g., sequential patterns of activity or state space trajectories, in favor of focusing on averaged levels of activity.

### Relevance to disease

Schizophrenia affects approximately 1% of the global population (Velligan and Rao, 2023). Cognitive deficits in schizophrenia affect up to 80% of patients, lack targeted FDA-approved treatments, and may stem from PVIN dysfunction (Hashimoto et al., 2003; Dienel et al., 2023; Sohal, 2024; Maroney, 2022; Bowie & Harvey, 2006). *Dlx5/6*^+/-^ mice do not model a specific genetic disruption linked to neuropsychiatric disease. That said, they do model disease-relevant relationships between abnormal PVIN development, deficient gamma synchrony, and cognitive inflexibility. In this context, it is notable that treating *Dlx5/6^+/-^* mutants with clonazepam seems to restore some aspects of normal function, specifically rescuing the dynamic encoding of error outcomes and outcome-independent activity patterns in the early RS period. Focusing on these specific readouts of circuit function and the mechanisms through which PVINs promote them may inform the development of novel therapies for addressing cognitive deficits associated with schizophrenia.

### Future Directions

In addition to the directions outlined above (measuring activity in specific classes of prefrontal neurons, examining neural dynamics during the pre-/and post-decision period), a central question is how clonazepam produces persistent changes in neural encoding and behavior? The persistence of clonazepam’s effects on neural encoding over 1 week later raises the possibility that some GABAergic synapses acutely enhanced by clonazepam may also undergo an enduring potentiation. This would be consistent with earlier studies identifying long-term potentiation in synapses from PVINs (Lourenço et al., 2014). Locating persistently strengthened synapses and/or other relevant substrates for clonazepam could inform the development of interventions promoting therapeutic network plasticity to improve cognition.

### Summary

Here, we leveraged mutant mice with abnormal PVIN development and reversible cognitive deficits to isolate aspects of prefrontal network encoding that are necessary for successful cognitive flexibility and critically dependent on intact inhibitory circuits. We find that the stable encoding of trial outcomes is not required for learning new rules. Instead, successful rule shift learning requires dynamic outcome representations and outcome-independent representations of the rule shift. Both of these depend on intact PVIN function. Pharmacologically enhancing inhibitory circuits restores these aspects of prefrontal encoding acutely and persistently, revealing correlates for therapeutic network plasticity.

## ACKNOWLEDGEMENTS

This work was supported by NIH grants R01NS116594 and R01MH106507.

## METHODS

### Animal care and surgical protocol

Guidelines from both the University of California, San Francisco’s Administrative Panels on Laboratory Animal Care and the National Institutes of Health were followed for animal care. Mice received ad libitum access to food and water and were housed in temperature-controlled conditions with a 12/12 light/dark cycle. *Dlx5/6^+/-^*mice were generated as described in Cho et al., 2015 and maintained on a mixed CD1 and C57Bl/6 background. All experiments were performed with *Dlx5/6^+/-^* mice and their age-matched wild-type littermates. Mice underwent stereotactic surgery to implant the virus and lens in preparation for mPFC micro-endoscopic imaging. While anesthetized with isoflurane, mice were injected with 600 µL of AAV9-Syn-jGcAMP7f (1:3 dilution, Addgene) at four depths in mPFC (+1.7 (AP), 0.3 (ML), and -2.75, -2.5, - 2.25, -2.0 (DV) in Bregma relative mm). 0.5 mm diameter GRIN lenses (Inscopix Inc) were implanted 0.3mm above the uppermost injection site, and at least four weeks elapsed before behavioral experiments to allow sufficient time for viral expression.

### Imaging site verification and Implant localization

After the conclusion of testing, mice were injected with a lethal dose of a ketamine-xylazine solution (Euthasol). Mice received a transcardial perfusion of a 4% paraformaldehyde (PFA) solution, followed by brain extraction. Slices were obtained using a Leica VT100S and then mounted on slides. Slide images were collected with a Keyence BZ-X All-in-One Fluorescence Microscope.

All mice, apart from 2 subjects, were verified to have implant tracts in mPFC. The location of viral fluorescence in the mPFC was used to confirm targeting in 1 subject where the implant impression could not be located. Additionally, due to an equipment malfunction during perfusion, another brain did not receive adequate perfusion and could not have its implant located.

### ‘Rule shifting’ task tests cognitive flexibility in mice

Our lab’s cognitive flexibility task (or rule-shifting task) has been described previously (Cho et al. 2015, 2020, 2023) but is summarized below. At the start of each trial, mice are moved from a ‘rest cage’ into the adjacent testing cage containing two small bowls. Each bowl contains a combination of one of two possible odors and one of two possible textures, meaning each trial presents two of four possible odor/texture combinations. A food reward (a peanut butter chip) is buried in bowls based on a rule, where a specific odor or texture is selected as the stimulus cue indicating the presence of a food reward (e.g., a bowl with sand will contain a food reward) (Fig. 1B).

We categorized trials as ‘error’ or ‘correct’ based on the mouse’s choice. If mice dug in a bowl for three cumulative seconds, we considered that bowl to be the mouse’s choice. Trials were categorized as errors if mice chose the non-rewarded bowl. In error trials, the reward bowl was removed immediately after a bowl choice; mice were allowed to dig in the non-reward bowl for at least 15 seconds. After both types of trials, mice were moved to the rest cage after they stopped digging in their chosen bowl (and after eating food reward during correct trials). During error trials, mice were kept in the rest cage for an extended inter-trial interval.

Each testing session is divided into the Initial Association (IA) rule period and the Rule shift (RS). During the IA period, mice first learn an initial rule in sensory modality 1 (e.g., that the bowl with sand will always contain a reward). Once the mouse achieves 80% correct performance over ten consecutive trials, it is deemed to have learned the IA rule. Mice then perform three more trials before the experiment enters the RS phase, and the rule changes to the *other* sensory modality (e.g., from sand signifying reward presence to coriander odor being the signal). This rule change is called the rule shift (**Figure 1B**).

Before being tested on the rule-shifting task, mice are single-housed and food-deprived for several days, with food placed inside the bowls that will later be used to contain reward during the main rule-shifting task. They then undergo a habituation session of 10 trials where reward placement is randomly assigned to one bowl. This teaches mice that only one bowl has a food reward in the testing environment.

### Single-cell resolution calcium imaging in mouse mPFC via 1-photon microendoscopy during the rule shifting task

We recorded calcium activity in mPFC via implanted GRIN lenses while mice performed the rule-shifting task on two consecutive days. On day 1, mice received a control intraperitoneal (i.p.) injection (volume: 0.01 ml/g) of saline (VEH). On day 2, mice received an intraperitoneal (i.p.) injection (volume: 0.01 ml/g) of clonazepam (at 0.0625 mg/kg) 30 minutes before starting the task. This dose causes no significant locomotor effects and is sub-sedative and sub-anxiolytic while improving task performance in *Dlx5/6^+/-^*mice (Cho et al., 2015), and was motivated by a study of Scn1a+/- mice, which used this dose for similar reasons in (Han et al., 2012). After two days of consecutive testing, we waited at least seven days to guarantee CLNZ clearance, then imaged HET mice for the ‘post-CLNZ’ condition. A head-mounted one-photon microendoscope (nVoke2, Inscopix Inc.) records fluorescence detected from an implanted GRIN lens in mPFC at 20 Hz. This raw fluorescence time-series data is spatially down-sampled, then spatial bandpass filtered (with a 0.03 pixel^-1^ high cutoff and a 0.008 pixel^-1^ low cutoff) and motion corrected.

### ROI detection and time-series creation

Putative ROIs labeling neurons in the recording were found using the EXTRACT algorithm (Inan et al., 2021). ROIs labeled by EXTRACT were first manually sorted to remove false-positive cells, then filtered with a custom spatial deduplication algorithm, which leveraged the distribution of correlations between spatially distant ROIs to remove false-positive cells. When ROIs were spatially adjacent, we iteratively removed the least active cell within that spatial radius until no adjacent cells were more correlated than the 95^th^ percentile of spatially distant cell pairs. Finally, this processed time series was converted to a binary raster of calcium events via a custom threshold-based event-detection algorithm (Frost et al., 2021).

### Ensemble permutation testing

We used permutation testing to identify ensembles of cells significantly activated above chance in particular stages of the task (e.g., late RS). We created a null distribution by circularly shuffling binary event matrices and quantifying mean activity in those shuffled datasets during the frames in the task stage of interest. We repeat this 1000 times to create null distributions of mean activity per neuron. We then identify those cells active above the 95^th^ percentile of the distribution of the shuffled data’s mean activity during that stage.

### Behavioral analysis

Video recordings were captured from two fixed vantage positions during experiments, with one camera mounted above the test cage and one mounted on the test cage (Anymaze 3). This recording was manually annotated for behavioral events (e.g., dig start) at a 1-second resolution. Timestamps for behaviors were then synced to the binary raster of calcium events using TTL pulses logged in the calcium data, occurring at the initiation of the behavioral recording. Timestamps were sorted into the pre-outcome (time points after the mouse enters the test cage but before the bowl choice), post-outcome (time points after the mouse chooses a bowl but before the mouse is removed from the test cage), and inter-trial interval (time points where the mouse is in the rest cage) corresponding to each trial of the recording session.

The analysis focused on the last 3 seconds of the pre-outcome before the bowl choice and the first 15 seconds following the mouse’s choice of the bowl. Because mice had varying numbers of trials necessary to reach the criterion on each rule, the analysis focused on the first or last five trials in each rule, referred to as the *Early* or *Late* stages. The *Early* stage was further divided into *Error* and *Correct* groups of trials, resulting in a total of 3 unique stages per rule, yielding six total task stages (**Figure 2D**). Because the late trials occurred when mice demonstrated knowledge of the rule and achieved the preset criteria, most late trials were correct trials, and the analysis of ‘Late’ trials excluded sporadically occurring error trials.

### Quantification and statistical analysis

Statistical analysis was performed in MATLAB and Python. The details of the analysis are provided in the main text. We categorize each day’s recording into groups designated as a combination of the subject’s genotype abbreviation (WT or HET) and the treatment on that day’s abbreviation (VEH, CLNZ, and postCLNZ), and refer to this label as the *genotype-treatment*. Analysis of calcium activity was restricted to the post-outcome section, which consisted of the frames in each trial between the start of a mouse digging in the chosen bowl and the mouse being removed from the test cage.

### Pseudopopulation Creation

To combine cells from various subjects into a single time series, we create a *pseudopopulation* consisting of all cells from a genotype-treatment significantly active in a given stage. As subjects can have different numbers of trials in each task section (e.g., Mouse 1 has 2 early IA errors, but Mouse 2 has 1), we resample with replacement to create a new bootstrapped time series for each cell. We combine those individual cell bootstrap samples across subjects into each ensemble pseudopopulation. For a cell A significantly active in stage N, we randomly sample with replacement from all trials that belong to stage N to create a new time series for cell A. We oversample to 400 time-bins (the maximum number of time-bins present in 5 trials, given 250ms time-bins and a maximum trial length of 15 seconds) to ensure all cells had equal frames. We perform this for all cells in the stage N ensemble. We then resample stage N ensemble cell activity from all other stages. This process repeats for all ensembles, all task stages, and all genotype-treatment groups. The stage of origin for each of these datasets is then decoded using a linear classifier. We repeat this bootstrapping process 1000 times and analyze those distributions of decoding accuracies.

### Robust Cohen’s d and comparing bootstrap distributions

Cohen’s *d* measures the standardized mean difference between two distributions, and is defined as: *d* = (*m*1 − *m*2)/ *SD_pooled_* (Cohen et al., 2013). We calculate the pooled standard deviation by finding 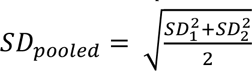. where *SD*_1_ and *SD*_2_ are the standard deviations of groups 1 and 2, respectively (Bonnet et al., 2008). Because we perform non-parametric bootstrapping, we compare groups using a robust estimator of Cohen’s *d*, which does not assume normality nor homogeneity of variance (Li et al., 2016). Robust Cohen’s *d* leverages winsorization to reduce the influence of outliers on distributions, which consists of finding the 20^th^ and 80^th^ percentiles of a dataset and then replacing all values greater than the 80^th^ percentile with the 80^th^ percentile value, and replacing all values below the 20^th^ percentile with the value at the 20^th^ percentile. We then calculated Cohen’s *d* on the now winsorized dataset (Algina, 2005).

To evaluate the practical significance of effect sizes/Cohen’s *d*, we use Cohen’s *U3*, which measures the overlap between two distributions. *U3* quantifies the % of the lesser distribution that is exceeded by the larger distribution (Cohen, 1988). *U3* is calculated by taking the inverse cumulative distribution function of the standard normal distribution at a given Cohen’s *d* value. We then subtract this value from 1 to obtain the probability of obtaining a difference this large or greater. We deem post hoc comparisons of bootstraps to be significantly different if the Cohen’s *d* yields a *U3* indicating a distribution overlap of 5% or less (d > 1.65). We then verify via permutation testing that the observed effect size exceeds the chance level (95th percentile) of effect size for group differences in label-shuffled data.

### Decoding analysis (SVM)

We used Python’s scikit-learn library to create support vector machines (SVMs) to classify pseudopopulation frames into 1 of 2 possible task stages per classifier run. For each pair of task stages being differentiated, we generate the resampled activity matrices of the ensemble of interest in each stage. We concatenated the two resampled task stage activity matrices for the ensemble pseudopopulation of interest. We dropped cells that were not active in one of the two task stages from data used for classification. Our resampling technique allowed us to generate class-balanced training and testing data. As this classification outputs binary classifications, we separately trained classifiers for each pair of task stages analyzed. L2-regularized SVMs with linear kernels were created using scikit-learn’s LinearSVC class with the following parameters: C= 1, max_iter=1000, penalty= ”l2”, dual=False. Cross validation was performed using scikit-learn’s StratifiedShuffleSplit cross-validator, which creates class-stratified random folds. We used 25% of data used as test set for each of 4 total cross-validation folds (with settings n_splits = 4, test_size = 0.25) (Pedregosa et al., 2011). SVM accuracy was quantified by averaging accuracy over all CV folds, for each bootstrap iteration (4 folds per iteration, 1000 total iterations).

### Cross-condition Generalization Performance (SVM)

Cross-condition generalization performance (CCGP) measures the ability of linear classifiers to generalize classification rules to classes not included in the training data. We train linear classifiers (SVMs) on along a specific dichotomy (e.g. error vs. correct trials during the early IA portion of the task), then test the performance of this SVM at classifying 2 test classes with a similar dichotomy using a different set of trials (e.g. error vs. correct trials during the early RS portion of the task). For testing, we evaluate the model’s performance at classifying test sets based on the training set, e.g., what % of early RS correct trials are classified as early IA correct trials). In this example, a model would have a CCGP of 1 if it classified all early RS error test samples as early IA error and all early RS correct test samples as early RS correct. In this way, CCGP measures whether the activity patterns which differentiate the two training classes also differentiate the training classes.

### Dimensionality Reduction

T-distributed stochastic neighborhood embedding (t-SNE) produces low-dimensional visualizations of a high-dimensional dataset, that balance conserving both local and global relationships between dataset samples. t-SNE was performed in Python, using the TSNE function from the scikit-learn package’s manifold module (version 1.5.2), setting the desired number of dimensions to 2 and the perplexity to 90, using cosine distance as the underlying distance metric, and using PCA embedding initialization (settings were: ’n_components’: 2, “perplexity”: 90, “method”: ’exact’, ’init’:’pca’). T-SNE was performed on each genotype-treatment’s mean neural stage-response matrix, a 6 x N matrix where N is the number of cells in each dataset, and 6 is the number of defined task stages. Matrix elements consisted of cell normalized event rates averaged across all trials in that stage.

### Neural Network Autoencoders

We train neural network undercomplete autoencoders using the *PyTorch* library in Python. Autoencoders are designed to minimize reconstruction loss of input data, given network-mediated compression of input data through a low-dimensional ‘bottleneck layer’. Therefore, the first and last layers of each autoencoder contained a number of units equal to the number of cells in the ensemble of interest. Cells were dropped if they had no active frames in at least 1 of the relevant task stages.

Between these input and output layers that matched the dimensionality of the input dataset, we standardized a hidden layer architecture across datasets. Model hidden layers consisted of a linear layer with 9 units) feeding into a bottleneck layer of 2 units, followed by another linear layer with 9 units. Neural Activation functions were set to *Mish*, a self-regularized non-monotonic activation function, which outperformed rectified linear units at minimizing reconstruction error on datasets used for validation. *Mish* is defined as *x tanh*(*softplus*(x*x*)) (Misra et al., 2020).

Models were trained to minimize the Mean Squared Error (MSE) reconstruction loss between the original input and reconstructed output matrices. Network gradient updates were performed using the *AdamW* optimizer. The number of units in the pre- and post-bottleneck hidden layers, learning rate, and choice of activation function were determined empirically using the *Optuna* hyperparameter optimization suite on a sample dataset, and held constant across all neural networks created.

### Cluster Separation Analysis

We calculated the Davies-Bouldin index for task stage neural activity manifolds by projecting 1000 bootstrap iterations, each using 400 samples of task stage activity, through autoencoders trained to minimize reconstruction error. The Davies-Bouldin index measures the ratio of inter-cluster distance to the sum of within-cluster scatters, averaged across all pairs of clusters. For a dataset with two clusters, *i* and *j*, the DB index can be defined as 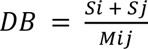, where *S_i_* = within-cluster scatter of cluster *i*, *S_j_* = within-cluster scatter of cluster j, and *M_ij_* measures the Euclidean distance between the centroids of clusters *i* and *j*. Lower values of the Davies-Bouldin index indicate greater separation between cluster centroids and decreased spread of points from their corresponding cluster centroid. We measure the DB index once per bootstrap iteration and use robust Cohen’s *d* to quantify the differences in these distributions.

## FIGURE LEGENDS

**SUPPLEMENTARY FIGURE 1:**
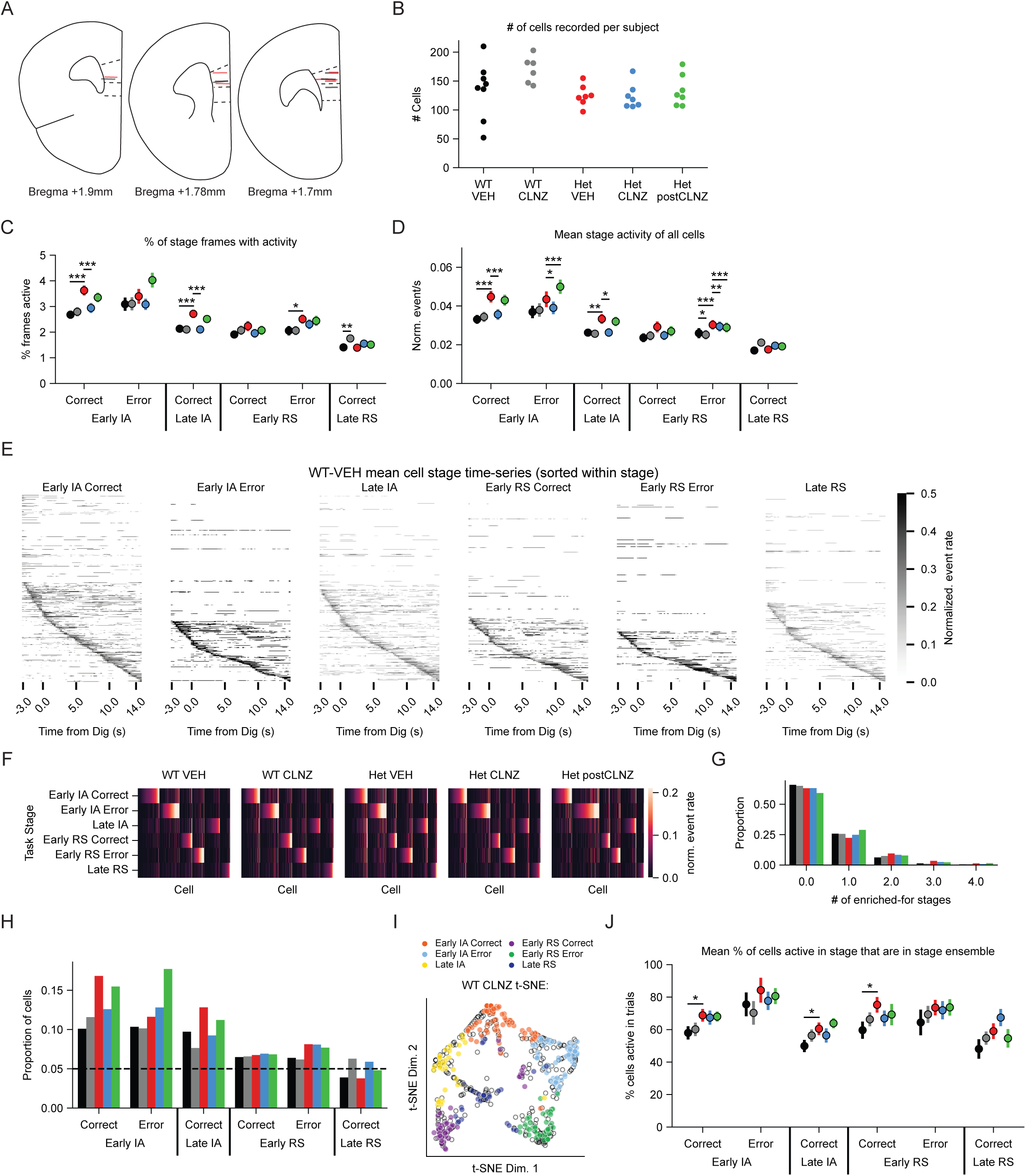
Histology, behavior, and task stage activity, including WT CLNZ A. GRIN lens implant locations, based on post-hoc histology. Colored lines indicate the approximate ventral edge of lens impressions. Grey lines correspond to WT mice, orange lines correspond to HET mice. Atlas images adapted from Paxinos & Franklin (2004). B. Number of cells analyzed per recording session, grouped by genotype-treatment condition. C. Average % of frames active per genotype-treatment by task stage. D. Mean normalized event rate of all cells recorded per stage E. Average timeseries of neuronal activity during different task stages in the WT VEH condition. Each row shows activity from one neuron, and neurons have been sorted by time of their maximum normalized activity (relative to the time of the bowl dig). F. Heat maps showing the activity of each neuron across task stages for each genotype / condition. Each column shows activity from one neuron, and neurons have been sorted based on the task stage in which they are most active. Neurons from recordings without early IA error trials are not included here for visual clarity. G. The distribution of the number of task stages in which neurons show enrichment (i.e., significantly elevated activity) is not significantly different across genotype-treatment conditions. H. The fraction of cells with significantly elevated activity (enrichment) as a function of trial type / task stage and genotype-treatment condition does not significantly vary across conditions. I. t-SNE plots showing the activity of WT CLNZ neurons in different task stages. Cells are color-coded based on the ensemble to which they belong, i.e., the task stage in which their activity is highest / significantly elevated. J. The average % of cells activate in each task stage that are members of that stage ensemble (i.e., have significantly elevated activity during that task stage). Panel C: p-value derived from permutation test with 1000 permutations. Panel D, J: p-value derived from the Mann-Whitney U test. Panels C, D, J: Points are mean +/- SEM. \**P* < 0.05, \*\**P* < 0.01, \*\*\**P* < 0.001.

**SUPPLEMENTARY FIGURE 2:**
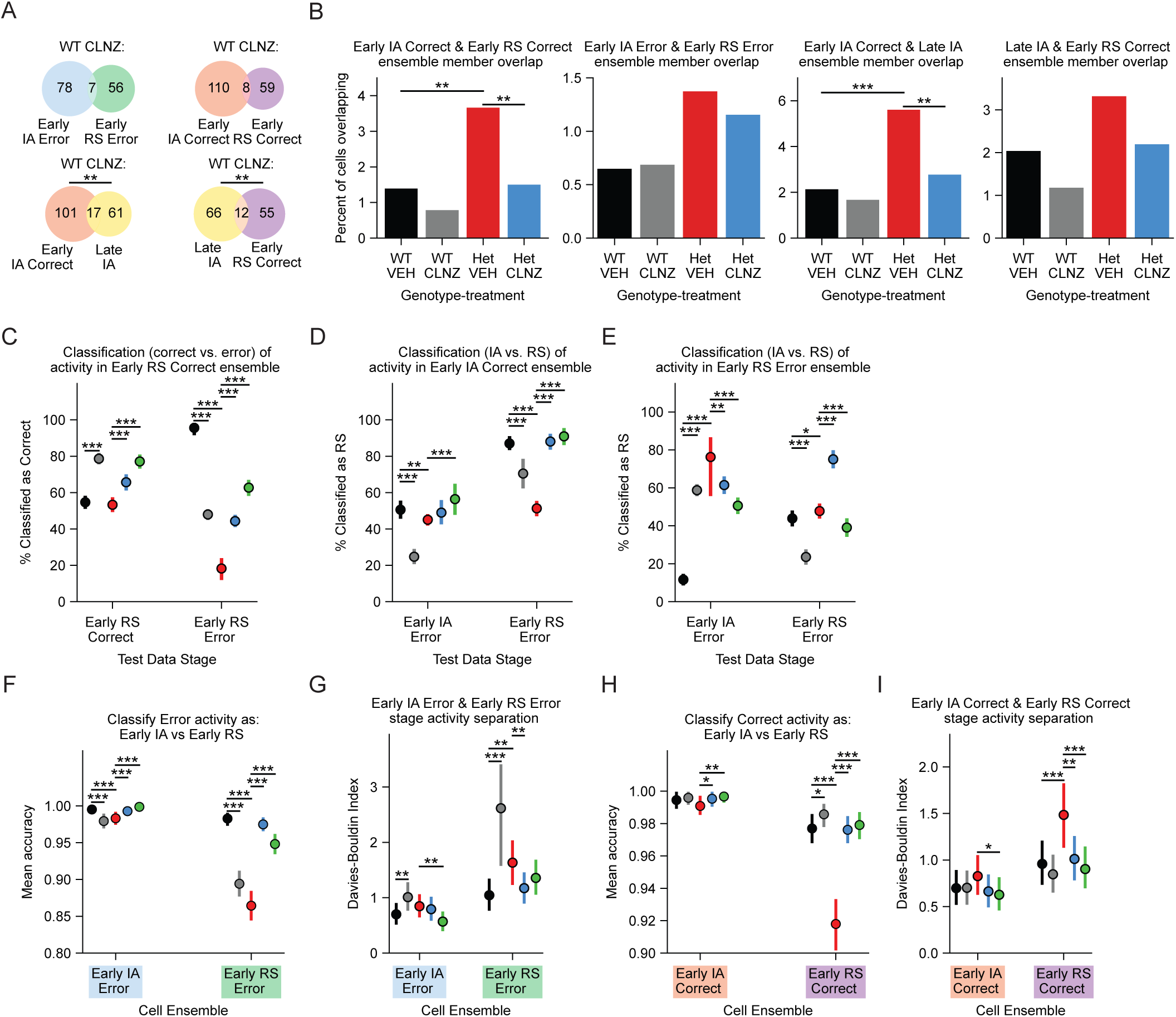
WT CLNZ classification is broadly similar to WT VEH A. Cell membership overlap between ensembles in the WT CLNZ condition. The late IA ensemble overlaps more than expected by chance with both the early IA correct and early RS correct ensembles. B. % of all recorded cells that overlap between the task stage ensembles analyzed, for WT VEH, WT CLNZ, Het VEH, and Het CLNZ. C. The percent of samples from early RS correct trials (left) or early RS error trials (right) that are classified as ‘Correct’ by CCG models that were trained on activity of the early RS correct ensemble during early IA correct vs. error trials, for each genotype-condition (including WT CLNZ). D. Similar to C, but for the percent of samples from early IA error or early RS error trials classified as ‘RS’ by CCG models trained on activity of the early IA correct ensemble during early IA vs. RS correct trials. E. Similar to D, but for CCG models trained activity of the early RS error ensemble. F. SVM accuracy at classifying early IA error vs. early RS error activity patterns, using the early IA error (left) or early RS error (right) ensembles. G. Quantification of stage activity separation in the latent space of autoencoders trained on activity during early IA vs. RS error trials, using either the early IA error (left) or early RS error (right) ensembles. H. Similar to F, but for SVM accuracy classifying early IA correct vs. early RS correct activity patterns, using the early IA correct (left) or early RS correct (right) ensembles. I. Similar to G, but for latent space cluster cohesion of activity during early IA vs. RS correct trials, using the early IA correct (left) or early RS correct (right) ensembles. Panels A, B: p-value derived from chi-squared test on ensemble member counts. Panels C-I: p-value calculated from Cohen’s U3, which is derived from each comparison’s Cohen’s *d.* Points and error bars show the mean +/- 75%ile of N = 1000 bootstrap distributions per genotype-treatment ensemble. \**P* < 0.05, \*\**P* < 0.01, \*\*\**P* < 0.001.

---

**Supplementary Table 1:**
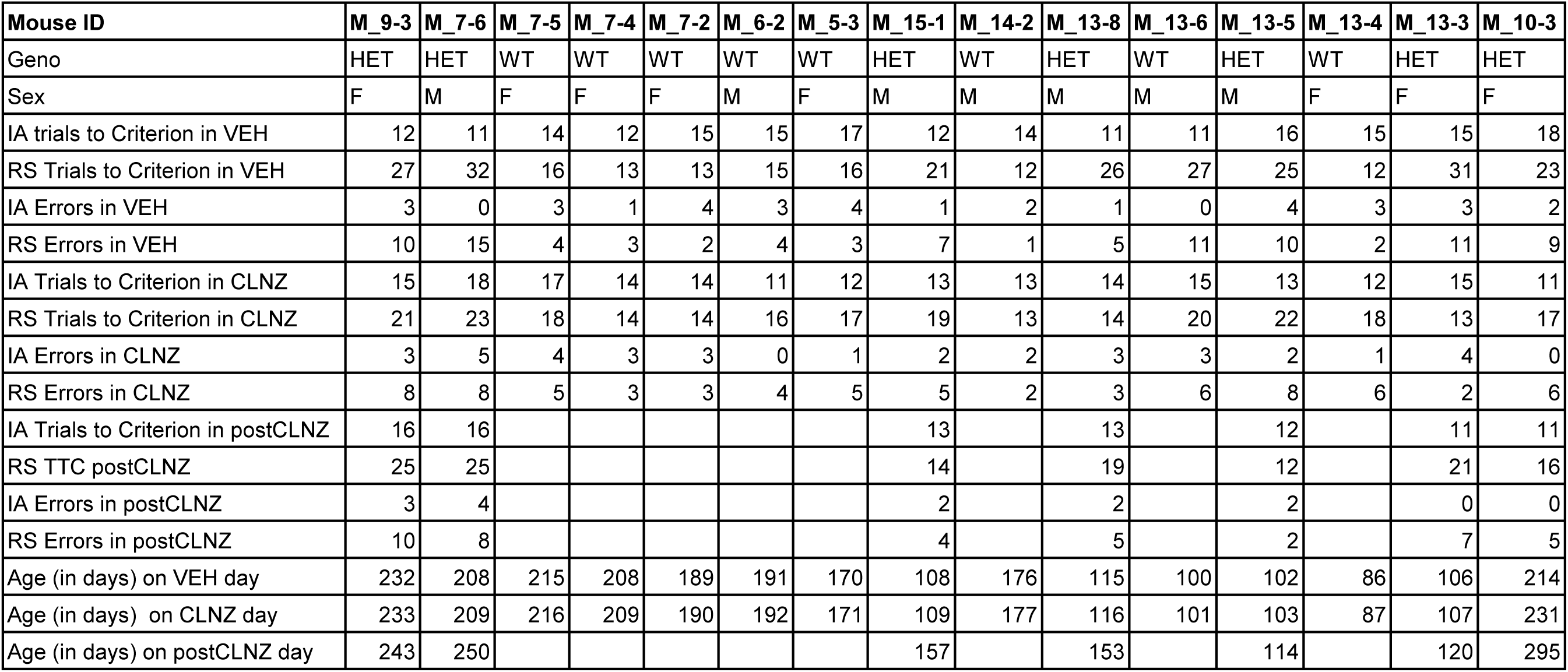
Subject Information.

**Supplementary Table 2:**
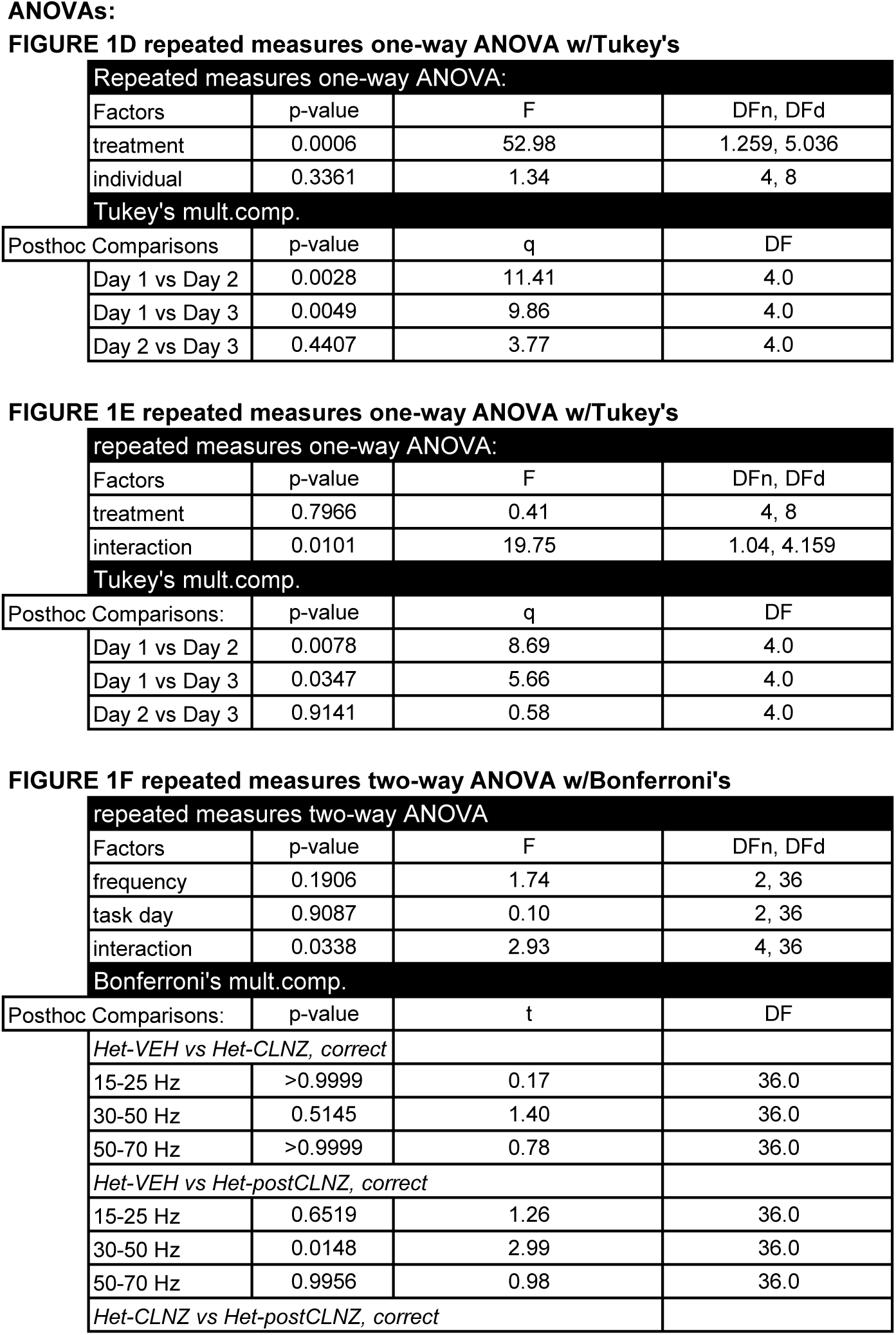
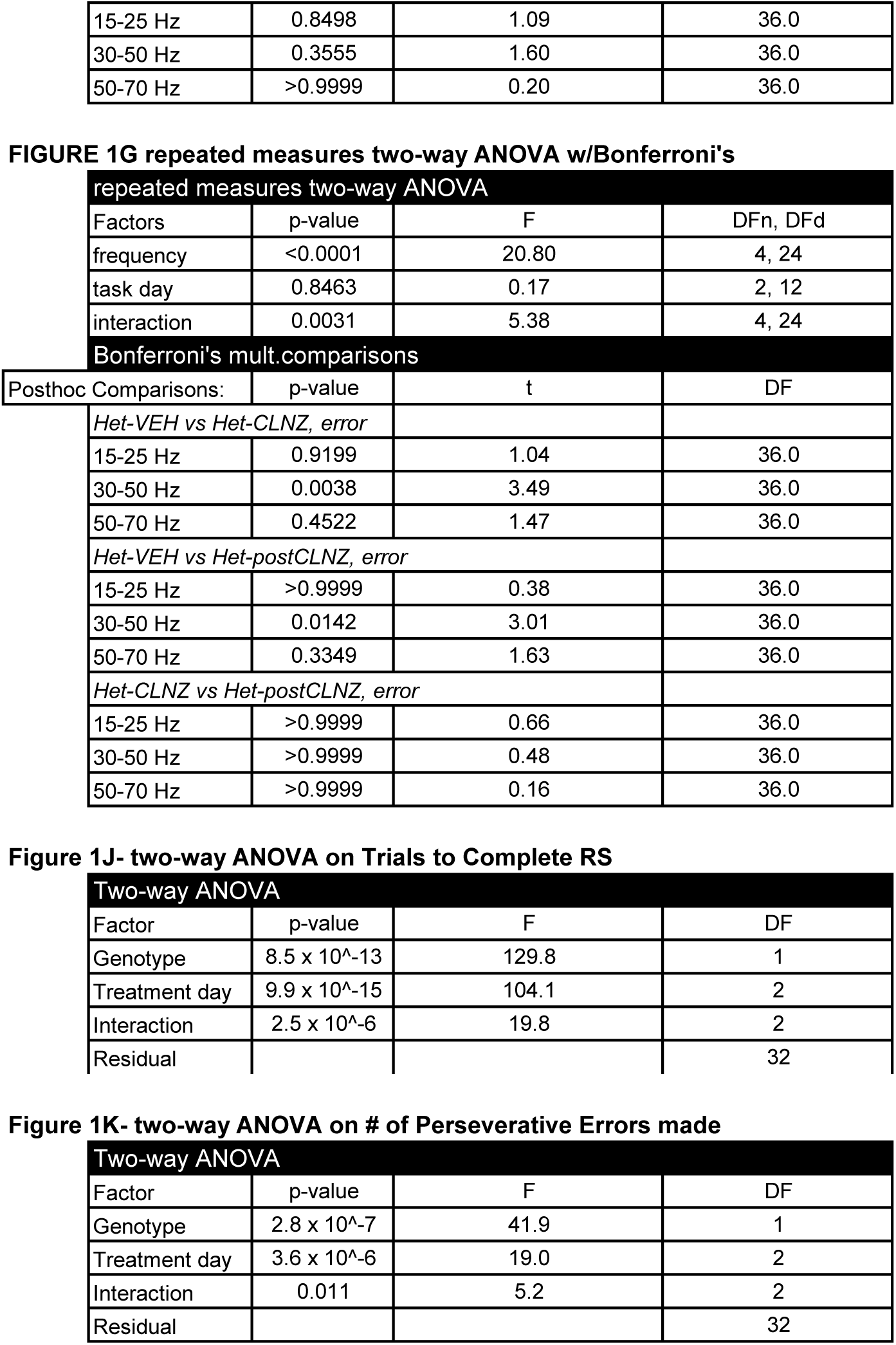
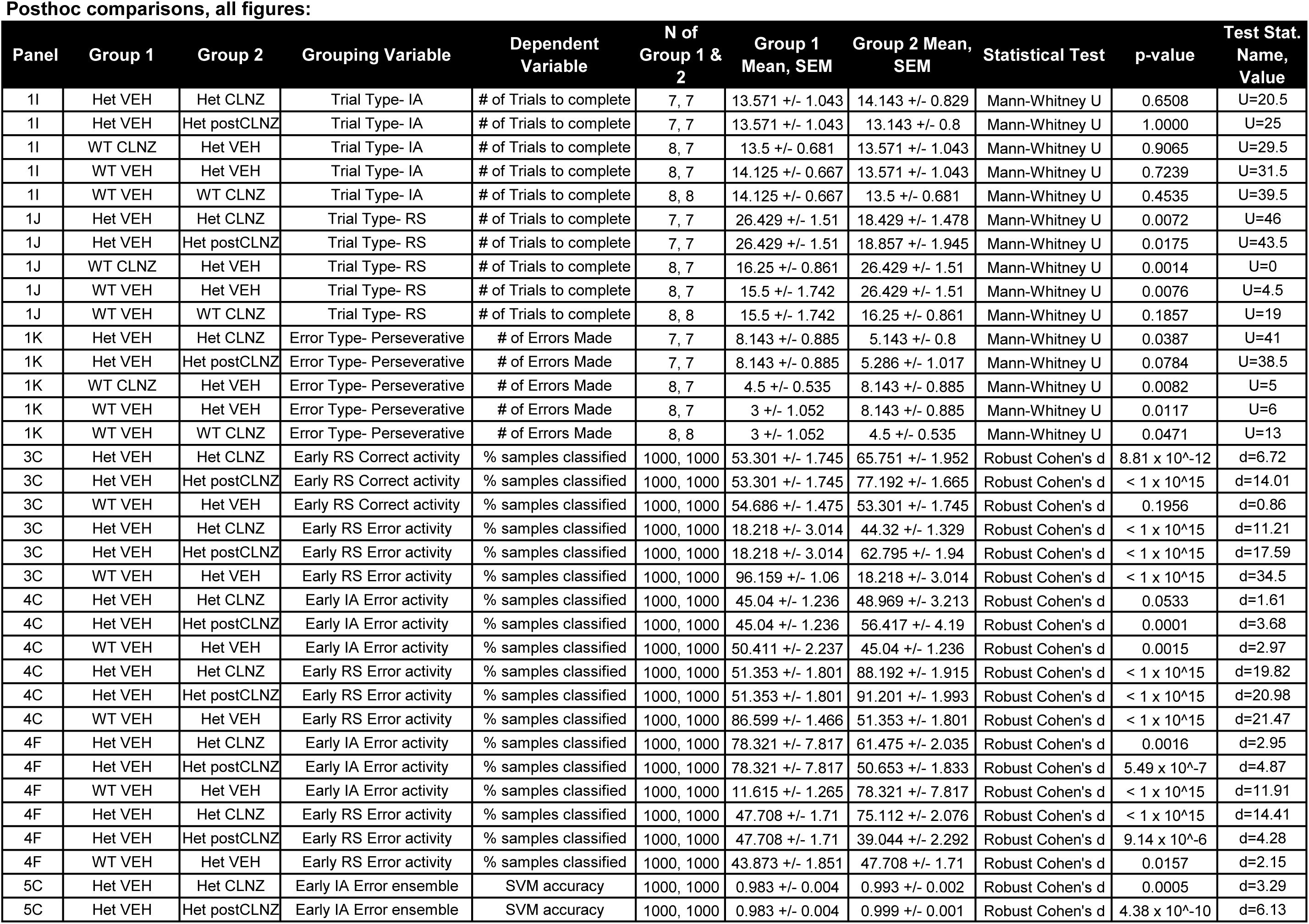
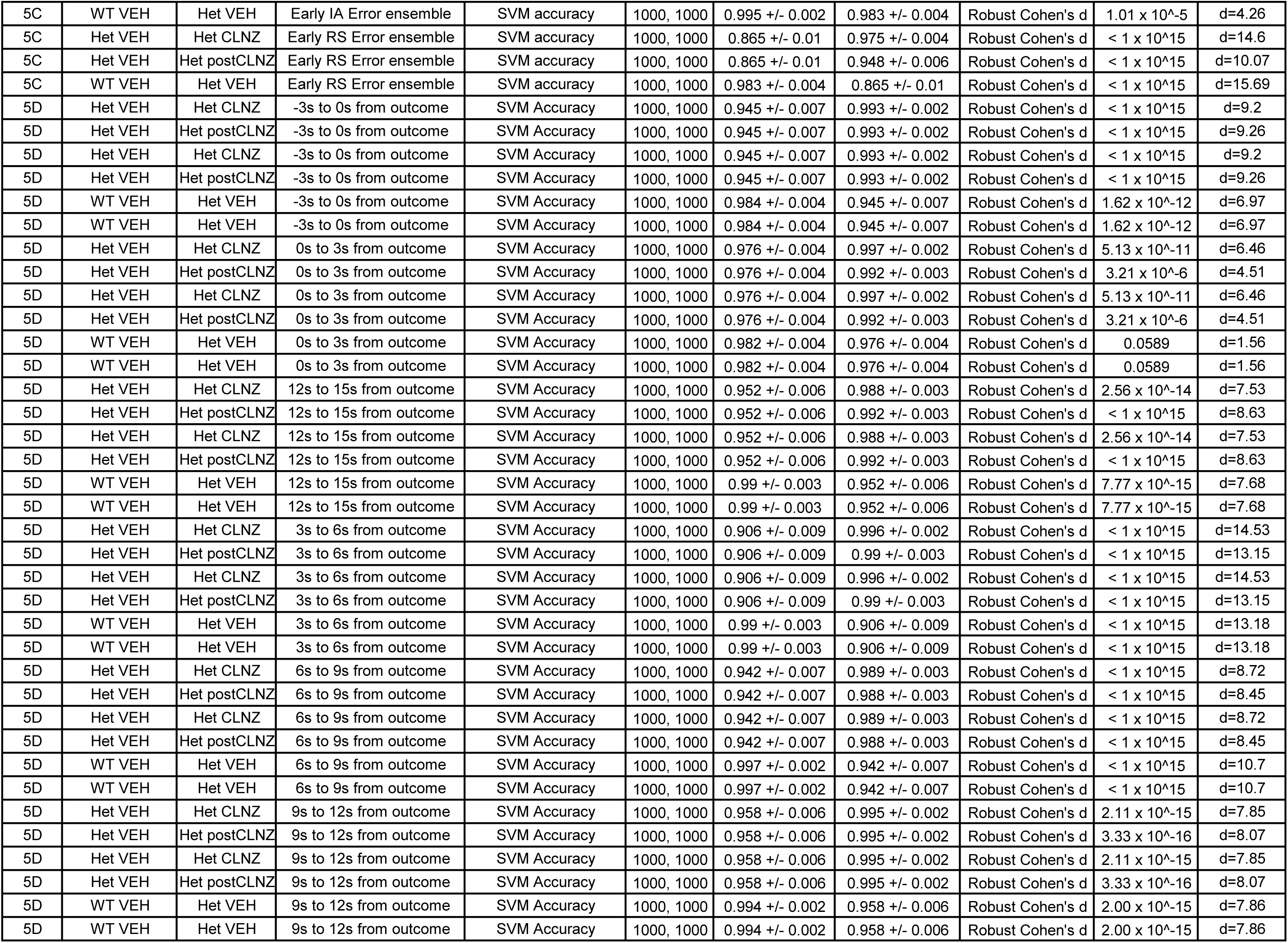
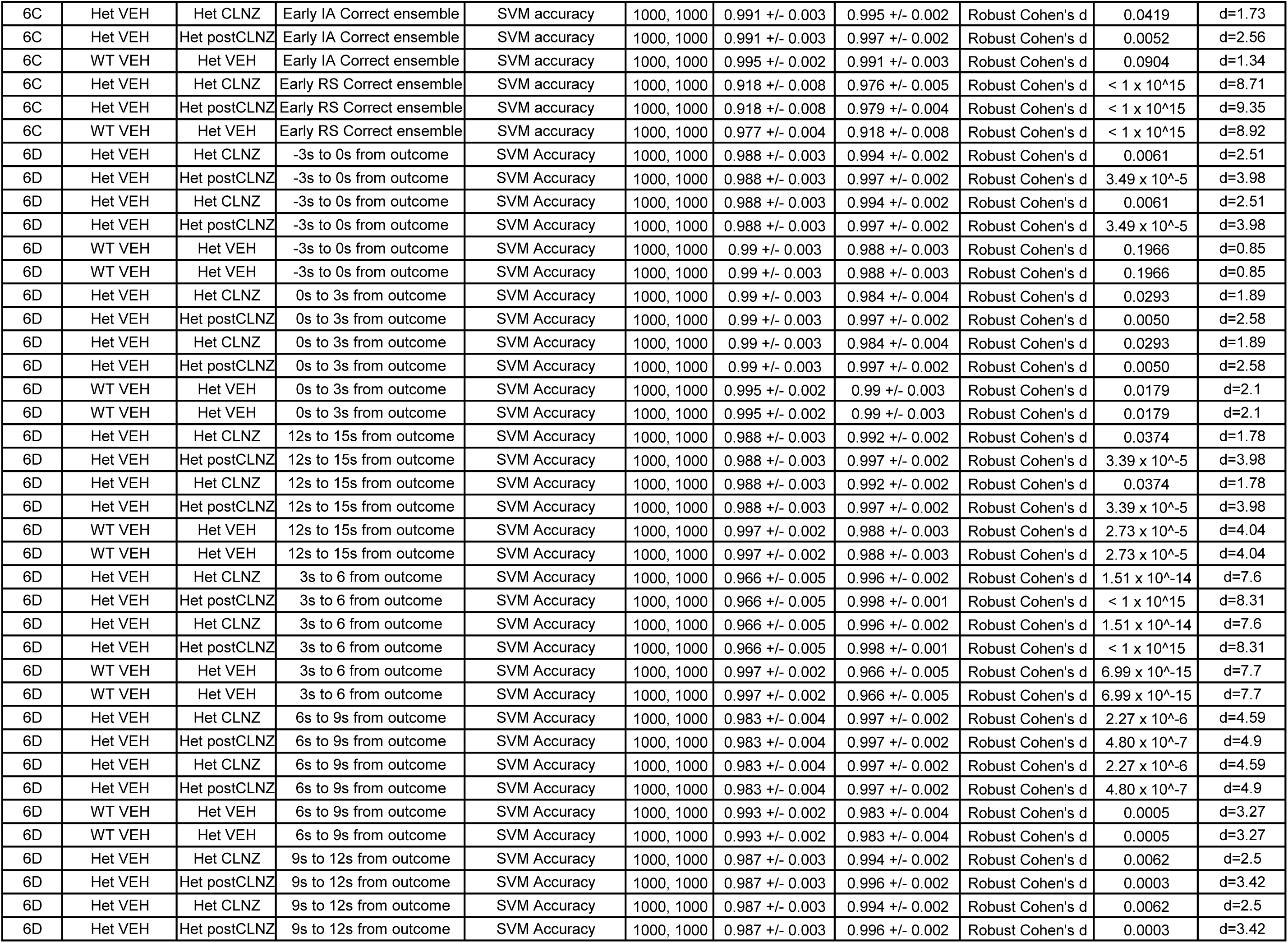
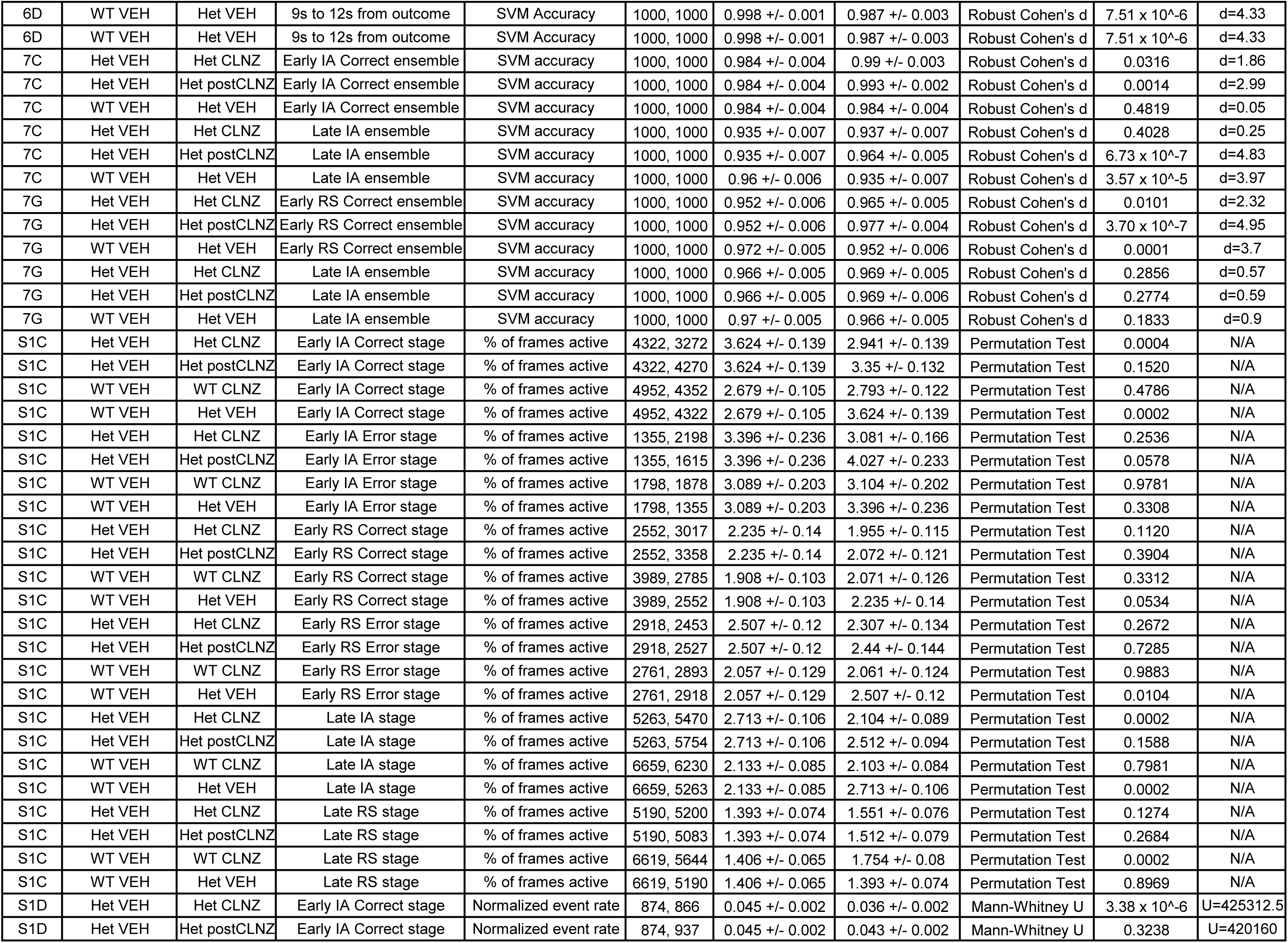
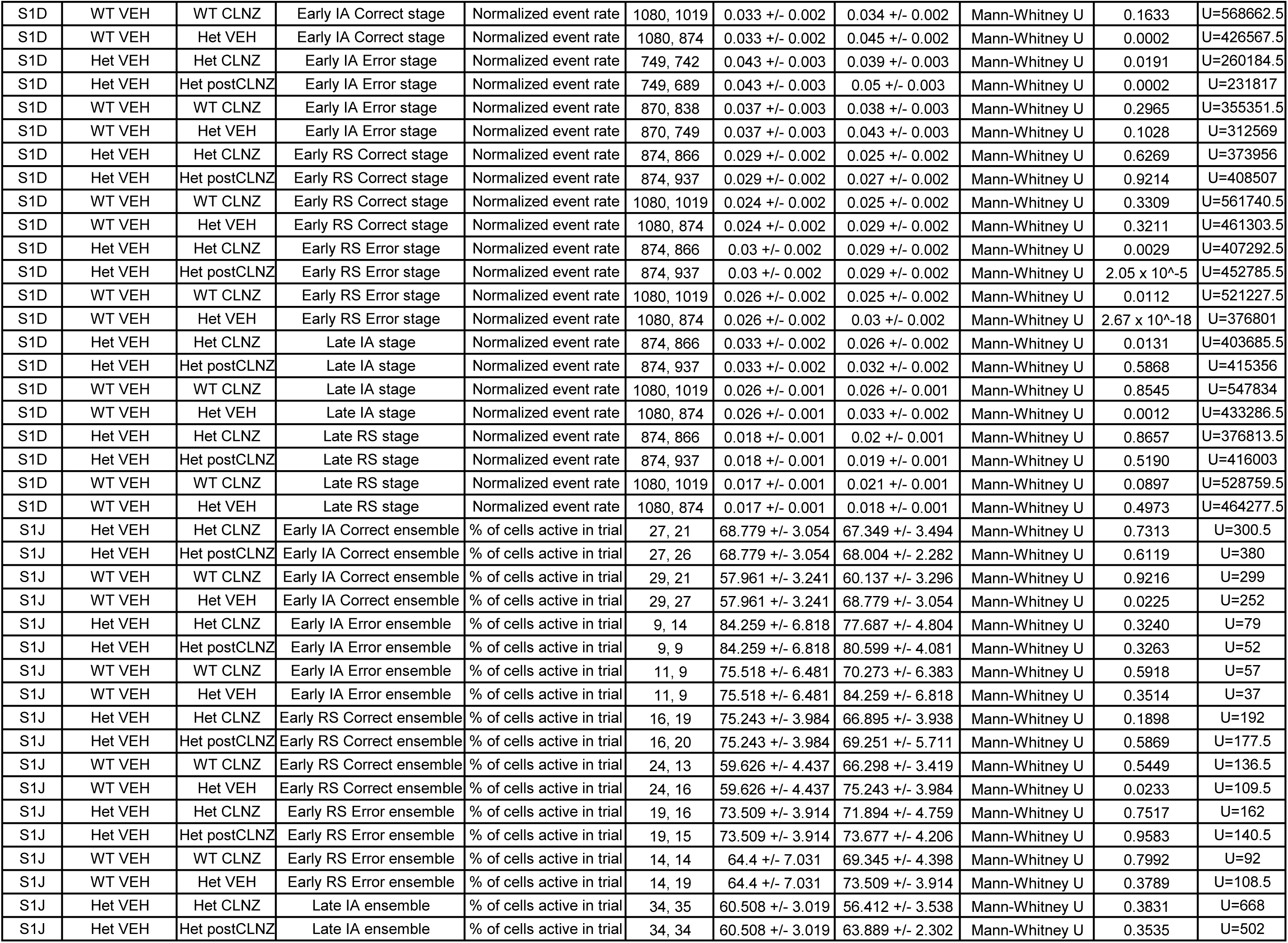
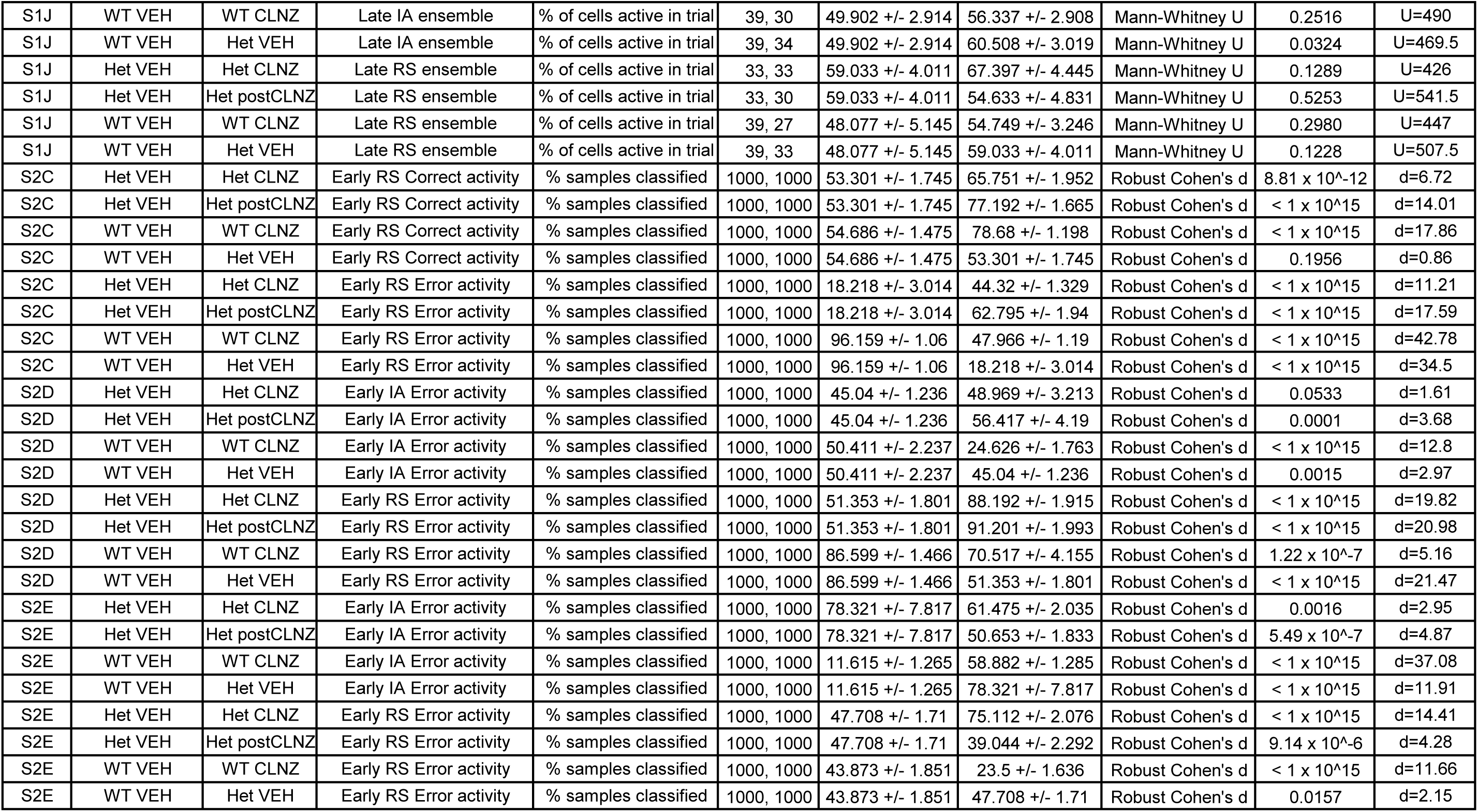
Figure Statistics.

**FIGURE 1D repeated measures one-way ANOVA w/Tukey's**
| Repeated measures one-way ANOVA: |  |  |  |
| --- | --- | --- | --- |
| Factors | p-value | F | DFn, DFd |
| treatment | 0.0006 | 52.98 | 1.259, 5.036 |
| individual | 0.3361 | 1.34 | 4, 8 |
| Tukey's mult.comp. |  |  |  |
| Posthoc Comparisons | p-value | q | DF |
| Day 1 vs Day 2 | 0.0028 | 11.41 | 4.0 |
| Day 1 vs Day 3 | 0.0049 | 9.86 | 4.0 |
| Day 2 vs Day 3 | 0.4407 | 3.77 | 4.0 |

**FIGURE 1E repeated measures one-way ANOVA w/Tukey's**
| repeated measures one-way ANOVA: |  |  |  |
| --- | --- | --- | --- |
| Factors | p-value | F | DFn, DFd |
| treatment | 0.7966 | 0.41 | 4, 8 |
| interaction | 0.0101 | 19.75 | 1.04, 4.159 |
| Tukey's mult.comp. |  |  |  |
| Posthoc Comparisons: | p-value | q | DF |
| Day 1 vs Day 2 | 0.0078 | 8.69 | 4.0 |
| Day 1 vs Day 3 | 0.0347 | 5.66 | 4.0 |
| Day 2 vs Day 3 | 0.9141 | 0.58 | 4.0 |

**FIGURE 1F repeated measures two-way ANOVA w/Bonferroni's**
| repeated measures two-way ANOVA |  |  |  |
| --- | --- | --- | --- |
| Factors | p-value | F | DFn, DFd |
| frequency | 0.1906 | 1.74 | 2, 36 |
| task day | 0.9087 | 0.10 | 2, 36 |
| interaction | 0.0338 | 2.93 | 4, 36 |
| Bonferroni's mult.comp. |  |  |  |
| Posthoc Comparisons: | p-value | t | DF |
| *Het-VEH vs Het-CLNZ, correct* |  |  |  |
| 15-25 Hz | >0.9999 | 0.17 | 36.0 |
| 30-50 Hz | 0.5145 | 1.40 | 36.0 |
| 50-70 Hz | >0.9999 | 0.78 | 36.0 |
| *Het-VEH vs Het-postCLNZ, correct* |  |  |  |
| 15-25 Hz | 0.6519 | 1.26 | 36.0 |
| 30-50 Hz | 0.0148 | 2.99 | 36.0 |
| 50-70 Hz | 0.9956 | 0.98 | 36.0 |
| *Het-CLNZ vs Het-postCLNZ, correct* |  |  |  |

Supplementary Table 2: Figure Statistics
|  |  |  |  |
| --- | --- | --- | --- |
| 15-25 Hz | 0.8498 | 1.09 | 36.0 |
| 30-50 Hz | 0.3555 | 1.60 | 36.0 |
| 50-70 Hz | >0.9999 | 0.20 | 36.0 |

**FIGURE 1G repeated measures two-way ANOVA w/Bonferroni's**
| repeated measures two-way ANOVA |  |  |  |
| --- | --- | --- | --- |
| Factors | p-value | F | DFn, DFd |
| frequency | <0.0001 | 20.80 | 4, 24 |
| task day | 0.8463 | 0.17 | 2, 12 |
| interaction | 0.0031 | 5.38 | 4, 24 |
| Bonferroni's mult.comparisons |  |  |  |
| Posthoc Comparisons: | p-value | t | DF |
| *Het-VEH vs Het-CLNZ, error* |  |  |  |
| 15-25 Hz | 0.9199 | 1.04 | 36.0 |
| 30-50 Hz | 0.0038 | 3.49 | 36.0 |
| 50-70 Hz | 0.4522 | 1.47 | 36.0 |
| *Het-VEH vs Het-postCLNZ, error* |  |  |  |
| 15-25 Hz | >0.9999 | 0.38 | 36.0 |
| 30-50 Hz | 0.0142 | 3.01 | 36.0 |
| 50-70 Hz | 0.3349 | 1.63 | 36.0 |
| *Het-CLNZ vs Het-postCLNZ, error* |  |  |  |
| 15-25 Hz | >0.9999 | 0.66 | 36.0 |
| 30-50 Hz | >0.9999 | 0.48 | 36.0 |
| 50-70 Hz | >0.9999 | 0.16 | 36.0 |

**Figure 1J- two-way ANOVA on Trials to Complete RS**
| Two-way ANOVA |  |  |  |
| --- | --- | --- | --- |
| Factor | p-value | F | DF |
| Genotype | $8.5 \times 10^{-13}$ | 129.8 | 1 |
| Treatment day | $9.9 \times 10^{-15}$ | 104.1 | 2 |
| Interaction | $2.5 \times 10^{-6}$ | 19.8 | 2 |
| Residual |  |  | 32 |

**Figure 1K- two-way ANOVA on # of Perseverative Errors made**
| Two-way ANOVA |  |  |  |
| --- | --- | --- | --- |
| Factor | p-value | F | DF |
| Genotype | $2.8 \times 10^{-7}$ | 41.9 | 1 |
| Treatment day | $3.6 \times 10^{-6}$ | 19.0 | 2 |
| Interaction | 0.011 | 5.2 | 2 |
| Residual |  |  | 32 |

Posthoc comparisons, all figures:
| Panel | Group 1 | Group 2 | Grouping Variable | Dependent Variable | N of Group 1 & 2 | Group 1 Mean, SEM | Group 2 Mean, SEM | Statistical Test | p-value | Test Stat. Name, Value |
| --- | --- | --- | --- | --- | --- | --- | --- | --- | --- | --- |
| 1I | Het VEH | Het CLNZ | Trial Type- IA | # of Trials to complete | 7, 7 | 13.571 +/- 1.043 | 14.143 +/- 0.829 | Mann-Whitney U | 0.6508 | U=20.5 |
| 1I | Het VEH | Het postCLNZ | Trial Type- IA | # of Trials to complete | 7, 7 | 13.571 +/- 1.043 | 13.143 +/- 0.8 | Mann-Whitney U | 1.0000 | U=25 |
| 1I | WT CLNZ | Het VEH | Trial Type- IA | # of Trials to complete | 8, 7 | 13.5 +/- 0.681 | 13.571 +/- 1.043 | Mann-Whitney U | 0.9065 | U=29.5 |
| 1I | WT VEH | Het VEH | Trial Type- IA | # of Trials to complete | 8, 7 | 14.125 +/- 0.667 | 13.571 +/- 1.043 | Mann-Whitney U | 0.7239 | U=31.5 |
| 1I | WT VEH | WT CLNZ | Trial Type- IA | # of Trials to complete | 8, 8 | 14.125 +/- 0.667 | 13.5 +/- 0.681 | Mann-Whitney U | 0.4535 | U=39.5 |
| 1J | Het VEH | Het CLNZ | Trial Type- RS | # of Trials to complete | 7, 7 | 26.429 +/- 1.51 | 18.429 +/- 1.478 | Mann-Whitney U | 0.0072 | U=46 |
| 1J | Het VEH | Het postCLNZ | Trial Type- RS | # of Trials to complete | 7, 7 | 26.429 +/- 1.51 | 18.857 +/- 1.945 | Mann-Whitney U | 0.0175 | U=43.5 |
| 1J | WT CLNZ | Het VEH | Trial Type- RS | # of Trials to complete | 8, 7 | 16.25 +/- 0.861 | 26.429 +/- 1.51 | Mann-Whitney U | 0.0014 | U=0 |
| 1J | WT VEH | Het VEH | Trial Type- RS | # of Trials to complete | 8, 7 | 15.5 +/- 1.742 | 26.429 +/- 1.51 | Mann-Whitney U | 0.0076 | U=4.5 |
| 1J | WT VEH | WT CLNZ | Trial Type- RS | # of Trials to complete | 8, 8 | 15.5 +/- 1.742 | 16.25 +/- 0.861 | Mann-Whitney U | 0.1857 | U=19 |
| 1K | Het VEH | Het CLNZ | Error Type- Perseverative | # of Errors Made | 7, 7 | 8.143 +/- 0.885 | 5.143 +/- 0.8 | Mann-Whitney U | 0.0387 | U=41 |
| 1K | Het VEH | Het postCLNZ | Error Type- Perseverative | # of Errors Made | 7, 7 | 8.143 +/- 0.885 | 5.286 +/- 1.017 | Mann-Whitney U | 0.0784 | U=38.5 |
| 1K | WT CLNZ | Het VEH | Error Type- Perseverative | # of Errors Made | 8, 7 | 4.5 +/- 0.535 | 8.143 +/- 0.885 | Mann-Whitney U | 0.0082 | U=5 |
| 1K | WT VEH | Het VEH | Error Type- Perseverative | # of Errors Made | 8, 7 | 3 +/- 1.052 | 8.143 +/- 0.885 | Mann-Whitney U | 0.0117 | U=6 |
| 1K | WT VEH | WT CLNZ | Error Type- Perseverative | # of Errors Made | 8, 8 | 3 +/- 1.052 | 4.5 +/- 0.535 | Mann-Whitney U | 0.0471 | U=13 |
| 3C | Het VEH | Het CLNZ | Early RS Correct activity | % samples classified | 1000, 1000 | 53.301 +/- 1.745 | 65.751 +/- 1.952 | Robust Cohen's d | 8.81 x 10<sup>-12</sup> | d=6.72 |
| 3C | Het VEH | Het postCLNZ | Early RS Correct activity | % samples classified | 1000, 1000 | 53.301 +/- 1.745 | 77.192 +/- 1.665 | Robust Cohen's d | < 1 x 10<sup>-15</sup> | d=14.01 |
| 3C | WT VEH | Het VEH | Early RS Correct activity | % samples classified | 1000, 1000 | 54.686 +/- 1.475 | 53.301 +/- 1.745 | Robust Cohen's d | 0.1956 | d=0.86 |
| 3C | Het VEH | Het CLNZ | Early RS Error activity | % samples classified | 1000, 1000 | 18.218 +/- 3.014 | 44.32 +/- 1.329 | Robust Cohen's d | < 1 x 10<sup>-15</sup> | d=11.21 |
| 3C | Het VEH | Het postCLNZ | Early RS Error activity | % samples classified | 1000, 1000 | 18.218 +/- 3.014 | 62.795 +/- 1.94 | Robust Cohen's d | < 1 x 10<sup>-15</sup> | d=17.59 |
| 3C | WT VEH | Het VEH | Early RS Error activity | % samples classified | 1000, 1000 | 96.159 +/- 1.06 | 18.218 +/- 3.014 | Robust Cohen's d | < 1 x 10<sup>-15</sup> | d=34.5 |
| 4C | Het VEH | Het CLNZ | Early IA Error activity | % samples classified | 1000, 1000 | 45.04 +/- 1.236 | 48.969 +/- 3.213 | Robust Cohen's d | 0.0533 | d=1.61 |
| 4C | Het VEH | Het postCLNZ | Early IA Error activity | % samples classified | 1000, 1000 | 45.04 +/- 1.236 | 56.417 +/- 4.19 | Robust Cohen's d | 0.0001 | d=3.68 |
| 4C | WT VEH | Het VEH | Early IA Error activity | % samples classified | 1000, 1000 | 50.411 +/- 2.237 | 45.04 +/- 1.236 | Robust Cohen's d | 0.0015 | d=2.97 |
| 4C | Het VEH | Het CLNZ | Early RS Error activity | % samples classified | 1000, 1000 | 51.353 +/- 1.801 | 88.192 +/- 1.915 | Robust Cohen's d | < 1 x 10<sup>-15</sup> | d=19.82 |
| 4C | Het VEH | Het postCLNZ | Early RS Error activity | % samples classified | 1000, 1000 | 51.353 +/- 1.801 | 91.201 +/- 1.993 | Robust Cohen's d | < 1 x 10<sup>-15</sup> | d=20.98 |
| 4C | WT VEH | Het VEH | Early RS Error activity | % samples classified | 1000, 1000 | 86.599 +/- 1.466 | 51.353 +/- 1.801 | Robust Cohen's d | < 1 x 10<sup>-15</sup> | d=21.47 |
| 4F | Het VEH | Het CLNZ | Early IA Error activity | % samples classified | 1000, 1000 | 78.321 +/- 7.817 | 61.475 +/- 2.035 | Robust Cohen's d | 0.0016 | d=2.95 |
| 4F | Het VEH | Het postCLNZ | Early IA Error activity | % samples classified | 1000, 1000 | 78.321 +/- 7.817 | 50.653 +/- 1.833 | Robust Cohen's d | 5.49 x 10<sup>-7</sup> | d=4.87 |
| 4F | WT VEH | Het VEH | Early IA Error activity | % samples classified | 1000, 1000 | 11.615 +/- 1.265 | 78.321 +/- 7.817 | Robust Cohen's d | < 1 x 10<sup>-15</sup> | d=11.91 |
| 4F | Het VEH | Het CLNZ | Early RS Error activity | % samples classified | 1000, 1000 | 47.708 +/- 1.71 | 75.112 +/- 2.076 | Robust Cohen's d | < 1 x 10<sup>-15</sup> | d=14.41 |
| 4F | Het VEH | Het postCLNZ | Early RS Error activity | % samples classified | 1000, 1000 | 47.708 +/- 1.71 | 39.044 +/- 2.292 | Robust Cohen's d | 9.14 x 10<sup>-6</sup> | d=4.28 |
| 4F | WT VEH | Het VEH | Early RS Error activity | % samples classified | 1000, 1000 | 43.873 +/- 1.851 | 47.708 +/- 1.71 | Robust Cohen's d | 0.0157 | d=2.15 |
| 5C | Het VEH | Het CLNZ | Early IA Error ensemble | SVM accuracy | 1000, 1000 | 0.983 +/- 0.004 | 0.993 +/- 0.002 | Robust Cohen's d | 0.0005 | d=3.29 |
| 5C | Het VEH | Het postCLNZ | Early IA Error ensemble | SVM accuracy | 1000, 1000 | 0.983 +/- 0.004 | 0.999 +/- 0.001 | Robust Cohen's d | 4.38 x 10<sup>-10</sup> | d=6.13 |

Supplementary Table 2: Figure Statistics
|  |  |  |  |  |  |  |  |  |  |  |
| --- | --- | --- | --- | --- | --- | --- | --- | --- | --- | --- |
| 5C | WT VEH | Het VEH | Early IA Error ensemble | SVM accuracy | 1000, 1000 | 0.995 +/- 0.002 | 0.983 +/- 0.004 | Robust Cohen's d | $1.01 \times 10^{-5}$ | d=4.26 |
| 5C | Het VEH | Het CLNZ | Early RS Error ensemble | SVM accuracy | 1000, 1000 | 0.865 +/- 0.01 | 0.975 +/- 0.004 | Robust Cohen's d | $< 1 \times 10^{-15}$ | d=14.6 |
| 5C | Het VEH | Het postCLNZ | Early RS Error ensemble | SVM accuracy | 1000, 1000 | 0.865 +/- 0.01 | 0.948 +/- 0.006 | Robust Cohen's d | $< 1 \times 10^{-15}$ | d=10.07 |
| 5C | WT VEH | Het VEH | Early RS Error ensemble | SVM accuracy | 1000, 1000 | 0.983 +/- 0.004 | 0.865 +/- 0.01 | Robust Cohen's d | $< 1 \times 10^{-15}$ | d=15.69 |
| 5D | Het VEH | Het CLNZ | -3s to 0s from outcome | SVM Accuracy | 1000, 1000 | 0.945 +/- 0.007 | 0.993 +/- 0.002 | Robust Cohen's d | $< 1 \times 10^{-15}$ | d=9.2 |
| 5D | Het VEH | Het postCLNZ | -3s to 0s from outcome | SVM Accuracy | 1000, 1000 | 0.945 +/- 0.007 | 0.993 +/- 0.002 | Robust Cohen's d | $< 1 \times 10^{-15}$ | d=9.26 |
| 5D | Het VEH | Het CLNZ | -3s to 0s from outcome | SVM Accuracy | 1000, 1000 | 0.945 +/- 0.007 | 0.993 +/- 0.002 | Robust Cohen's d | $< 1 \times 10^{-15}$ | d=9.2 |
| 5D | Het VEH | Het postCLNZ | -3s to 0s from outcome | SVM Accuracy | 1000, 1000 | 0.945 +/- 0.007 | 0.993 +/- 0.002 | Robust Cohen's d | $< 1 \times 10^{-15}$ | d=9.26 |
| 5D | WT VEH | Het VEH | -3s to 0s from outcome | SVM Accuracy | 1000, 1000 | 0.984 +/- 0.004 | 0.945 +/- 0.007 | Robust Cohen's d | $1.62 \times 10^{-12}$ | d=6.97 |
| 5D | WT VEH | Het VEH | -3s to 0s from outcome | SVM Accuracy | 1000, 1000 | 0.984 +/- 0.004 | 0.945 +/- 0.007 | Robust Cohen's d | $1.62 \times 10^{-12}$ | d=6.97 |
| 5D | Het VEH | Het CLNZ | 0s to 3s from outcome | SVM Accuracy | 1000, 1000 | 0.976 +/- 0.004 | 0.997 +/- 0.002 | Robust Cohen's d | $5.13 \times 10^{-11}$ | d=6.46 |
| 5D | Het VEH | Het postCLNZ | 0s to 3s from outcome | SVM Accuracy | 1000, 1000 | 0.976 +/- 0.004 | 0.992 +/- 0.003 | Robust Cohen's d | $3.21 \times 10^{-6}$ | d=4.51 |
| 5D | Het VEH | Het CLNZ | 0s to 3s from outcome | SVM Accuracy | 1000, 1000 | 0.976 +/- 0.004 | 0.997 +/- 0.002 | Robust Cohen's d | $5.13 \times 10^{-11}$ | d=6.46 |
| 5D | Het VEH | Het postCLNZ | 0s to 3s from outcome | SVM Accuracy | 1000, 1000 | 0.976 +/- 0.004 | 0.992 +/- 0.003 | Robust Cohen's d | $3.21 \times 10^{-6}$ | d=4.51 |
| 5D | WT VEH | Het VEH | 0s to 3s from outcome | SVM Accuracy | 1000, 1000 | 0.982 +/- 0.004 | 0.976 +/- 0.004 | Robust Cohen's d | 0.0589 | d=1.56 |
| 5D | WT VEH | Het VEH | 0s to 3s from outcome | SVM Accuracy | 1000, 1000 | 0.982 +/- 0.004 | 0.976 +/- 0.004 | Robust Cohen's d | 0.0589 | d=1.56 |
| 5D | Het VEH | Het CLNZ | 12s to 15s from outcome | SVM Accuracy | 1000, 1000 | 0.952 +/- 0.006 | 0.988 +/- 0.003 | Robust Cohen's d | $2.56 \times 10^{-14}$ | d=7.53 |
| 5D | Het VEH | Het postCLNZ | 12s to 15s from outcome | SVM Accuracy | 1000, 1000 | 0.952 +/- 0.006 | 0.992 +/- 0.003 | Robust Cohen's d | $< 1 \times 10^{-15}$ | d=8.63 |
| 5D | Het VEH | Het CLNZ | 12s to 15s from outcome | SVM Accuracy | 1000, 1000 | 0.952 +/- 0.006 | 0.988 +/- 0.003 | Robust Cohen's d | $2.56 \times 10^{-14}$ | d=7.53 |
| 5D | Het VEH | Het postCLNZ | 12s to 15s from outcome | SVM Accuracy | 1000, 1000 | 0.952 +/- 0.006 | 0.992 +/- 0.003 | Robust Cohen's d | $< 1 \times 10^{-15}$ | d=8.63 |
| 5D | WT VEH | Het VEH | 12s to 15s from outcome | SVM Accuracy | 1000, 1000 | 0.99 +/- 0.003 | 0.952 +/- 0.006 | Robust Cohen's d | $7.77 \times 10^{-15}$ | d=7.68 |
| 5D | WT VEH | Het VEH | 12s to 15s from outcome | SVM Accuracy | 1000, 1000 | 0.99 +/- 0.003 | 0.952 +/- 0.006 | Robust Cohen's d | $7.77 \times 10^{-15}$ | d=7.68 |
| 5D | Het VEH | Het CLNZ | 3s to 6s from outcome | SVM Accuracy | 1000, 1000 | 0.906 +/- 0.009 | 0.996 +/- 0.002 | Robust Cohen's d | $< 1 \times 10^{-15}$ | d=14.53 |
| 5D | Het VEH | Het postCLNZ | 3s to 6s from outcome | SVM Accuracy | 1000, 1000 | 0.906 +/- 0.009 | 0.99 +/- 0.003 | Robust Cohen's d | $< 1 \times 10^{-15}$ | d=13.15 |
| 5D | Het VEH | Het CLNZ | 3s to 6s from outcome | SVM Accuracy | 1000, 1000 | 0.906 +/- 0.009 | 0.996 +/- 0.002 | Robust Cohen's d | $< 1 \times 10^{-15}$ | d=14.53 |
| 5D | Het VEH | Het postCLNZ | 3s to 6s from outcome | SVM Accuracy | 1000, 1000 | 0.906 +/- 0.009 | 0.99 +/- 0.003 | Robust Cohen's d | $< 1 \times 10^{-15}$ | d=13.15 |
| 5D | WT VEH | Het VEH | 3s to 6s from outcome | SVM Accuracy | 1000, 1000 | 0.99 +/- 0.003 | 0.906 +/- 0.009 | Robust Cohen's d | $< 1 \times 10^{-15}$ | d=13.18 |
| 5D | WT VEH | Het VEH | 3s to 6s from outcome | SVM Accuracy | 1000, 1000 | 0.99 +/- 0.003 | 0.906 +/- 0.009 | Robust Cohen's d | $< 1 \times 10^{-15}$ | d=13.18 |
| 5D | Het VEH | Het CLNZ | 6s to 9s from outcome | SVM Accuracy | 1000, 1000 | 0.942 +/- 0.007 | 0.989 +/- 0.003 | Robust Cohen's d | $< 1 \times 10^{-15}$ | d=8.72 |
| 5D | Het VEH | Het postCLNZ | 6s to 9s from outcome | SVM Accuracy | 1000, 1000 | 0.942 +/- 0.007 | 0.988 +/- 0.003 | Robust Cohen's d | $< 1 \times 10^{-15}$ | d=8.45 |
| 5D | Het VEH | Het CLNZ | 6s to 9s from outcome | SVM Accuracy | 1000, 1000 | 0.942 +/- 0.007 | 0.989 +/- 0.003 | Robust Cohen's d | $< 1 \times 10^{-15}$ | d=8.72 |
| 5D | Het VEH | Het postCLNZ | 6s to 9s from outcome | SVM Accuracy | 1000, 1000 | 0.942 +/- 0.007 | 0.988 +/- 0.003 | Robust Cohen's d | $< 1 \times 10^{-15}$ | d=8.45 |
| 5D | WT VEH | Het VEH | 6s to 9s from outcome | SVM Accuracy | 1000, 1000 | 0.997 +/- 0.002 | 0.942 +/- 0.007 | Robust Cohen's d | $< 1 \times 10^{-15}$ | d=10.7 |
| 5D | WT VEH | Het VEH | 6s to 9s from outcome | SVM Accuracy | 1000, 1000 | 0.997 +/- 0.002 | 0.942 +/- 0.007 | Robust Cohen's d | $< 1 \times 10^{-15}$ | d=10.7 |
| 5D | Het VEH | Het CLNZ | 9s to 12s from outcome | SVM Accuracy | 1000, 1000 | 0.958 +/- 0.006 | 0.995 +/- 0.002 | Robust Cohen's d | $2.11 \times 10^{-15}$ | d=7.85 |
| 5D | Het VEH | Het postCLNZ | 9s to 12s from outcome | SVM Accuracy | 1000, 1000 | 0.958 +/- 0.006 | 0.995 +/- 0.002 | Robust Cohen's d | $3.33 \times 10^{-16}$ | d=8.07 |
| 5D | Het VEH | Het CLNZ | 9s to 12s from outcome | SVM Accuracy | 1000, 1000 | 0.958 +/- 0.006 | 0.995 +/- 0.002 | Robust Cohen's d | $2.11 \times 10^{-15}$ | d=7.85 |
| 5D | Het VEH | Het postCLNZ | 9s to 12s from outcome | SVM Accuracy | 1000, 1000 | 0.958 +/- 0.006 | 0.995 +/- 0.002 | Robust Cohen's d | $3.33 \times 10^{-16}$ | d=8.07 |
| 5D | WT VEH | Het VEH | 9s to 12s from outcome | SVM Accuracy | 1000, 1000 | 0.994 +/- 0.002 | 0.958 +/- 0.006 | Robust Cohen's d | $2.00 \times 10^{-15}$ | d=7.86 |
| 5D | WT VEH | Het VEH | 9s to 12s from outcome | SVM Accuracy | 1000, 1000 | 0.994 +/- 0.002 | 0.958 +/- 0.006 | Robust Cohen's d | $2.00 \times 10^{-15}$ | d=7.86 |

Supplementary Table 2: Figure Statistics
|  |  |  |  |  |  |  |  |  |  |  |
| --- | --- | --- | --- | --- | --- | --- | --- | --- | --- | --- |
| 6C | Het VEH | Het CLNZ | Early IA Correct ensemble | SVM accuracy | 1000, 1000 | 0.991 +/- 0.003 | 0.995 +/- 0.002 | Robust Cohen's d | 0.0419 | d=1.73 |
| 6C | Het VEH | Het postCLNZ | Early IA Correct ensemble | SVM accuracy | 1000, 1000 | 0.991 +/- 0.003 | 0.997 +/- 0.002 | Robust Cohen's d | 0.0052 | d=2.56 |
| 6C | WT VEH | Het VEH | Early IA Correct ensemble | SVM accuracy | 1000, 1000 | 0.995 +/- 0.002 | 0.991 +/- 0.003 | Robust Cohen's d | 0.0904 | d=1.34 |
| 6C | Het VEH | Het CLNZ | Early RS Correct ensemble | SVM accuracy | 1000, 1000 | 0.918 +/- 0.008 | 0.976 +/- 0.005 | Robust Cohen's d | < 1 x 10<sup>-15</sup> | d=8.71 |
| 6C | Het VEH | Het postCLNZ | Early RS Correct ensemble | SVM accuracy | 1000, 1000 | 0.918 +/- 0.008 | 0.979 +/- 0.004 | Robust Cohen's d | < 1 x 10<sup>-15</sup> | d=9.35 |
| 6C | WT VEH | Het VEH | Early RS Correct ensemble | SVM accuracy | 1000, 1000 | 0.977 +/- 0.004 | 0.918 +/- 0.008 | Robust Cohen's d | < 1 x 10<sup>-15</sup> | d=8.92 |
| 6D | Het VEH | Het CLNZ | -3s to 0s from outcome | SVM Accuracy | 1000, 1000 | 0.988 +/- 0.003 | 0.994 +/- 0.002 | Robust Cohen's d | 0.0061 | d=2.51 |
| 6D | Het VEH | Het postCLNZ | -3s to 0s from outcome | SVM Accuracy | 1000, 1000 | 0.988 +/- 0.003 | 0.997 +/- 0.002 | Robust Cohen's d | 3.49 x 10<sup>-5</sup> | d=3.98 |
| 6D | Het VEH | Het CLNZ | -3s to 0s from outcome | SVM Accuracy | 1000, 1000 | 0.988 +/- 0.003 | 0.994 +/- 0.002 | Robust Cohen's d | 0.0061 | d=2.51 |
| 6D | Het VEH | Het postCLNZ | -3s to 0s from outcome | SVM Accuracy | 1000, 1000 | 0.988 +/- 0.003 | 0.997 +/- 0.002 | Robust Cohen's d | 3.49 x 10<sup>-5</sup> | d=3.98 |
| 6D | WT VEH | Het VEH | -3s to 0s from outcome | SVM Accuracy | 1000, 1000 | 0.99 +/- 0.003 | 0.988 +/- 0.003 | Robust Cohen's d | 0.1966 | d=0.85 |
| 6D | WT VEH | Het VEH | -3s to 0s from outcome | SVM Accuracy | 1000, 1000 | 0.99 +/- 0.003 | 0.988 +/- 0.003 | Robust Cohen's d | 0.1966 | d=0.85 |
| 6D | Het VEH | Het CLNZ | 0s to 3s from outcome | SVM Accuracy | 1000, 1000 | 0.99 +/- 0.003 | 0.984 +/- 0.004 | Robust Cohen's d | 0.0293 | d=1.89 |
| 6D | Het VEH | Het postCLNZ | 0s to 3s from outcome | SVM Accuracy | 1000, 1000 | 0.99 +/- 0.003 | 0.997 +/- 0.002 | Robust Cohen's d | 0.0050 | d=2.58 |
| 6D | Het VEH | Het CLNZ | 0s to 3s from outcome | SVM Accuracy | 1000, 1000 | 0.99 +/- 0.003 | 0.984 +/- 0.004 | Robust Cohen's d | 0.0293 | d=1.89 |
| 6D | Het VEH | Het postCLNZ | 0s to 3s from outcome | SVM Accuracy | 1000, 1000 | 0.99 +/- 0.003 | 0.997 +/- 0.002 | Robust Cohen's d | 0.0050 | d=2.58 |
| 6D | WT VEH | Het VEH | 0s to 3s from outcome | SVM Accuracy | 1000, 1000 | 0.995 +/- 0.002 | 0.99 +/- 0.003 | Robust Cohen's d | 0.0179 | d=2.1 |
| 6D | WT VEH | Het VEH | 0s to 3s from outcome | SVM Accuracy | 1000, 1000 | 0.995 +/- 0.002 | 0.99 +/- 0.003 | Robust Cohen's d | 0.0179 | d=2.1 |
| 6D | Het VEH | Het CLNZ | 12s to 15s from outcome | SVM Accuracy | 1000, 1000 | 0.988 +/- 0.003 | 0.992 +/- 0.002 | Robust Cohen's d | 0.0374 | d=1.78 |
| 6D | Het VEH | Het postCLNZ | 12s to 15s from outcome | SVM Accuracy | 1000, 1000 | 0.988 +/- 0.003 | 0.997 +/- 0.002 | Robust Cohen's d | 3.39 x 10<sup>-5</sup> | d=3.98 |
| 6D | Het VEH | Het CLNZ | 12s to 15s from outcome | SVM Accuracy | 1000, 1000 | 0.988 +/- 0.003 | 0.992 +/- 0.002 | Robust Cohen's d | 0.0374 | d=1.78 |
| 6D | Het VEH | Het postCLNZ | 12s to 15s from outcome | SVM Accuracy | 1000, 1000 | 0.988 +/- 0.003 | 0.997 +/- 0.002 | Robust Cohen's d | 3.39 x 10<sup>-5</sup> | d=3.98 |
| 6D | WT VEH | Het VEH | 12s to 15s from outcome | SVM Accuracy | 1000, 1000 | 0.997 +/- 0.002 | 0.988 +/- 0.003 | Robust Cohen's d | 2.73 x 10<sup>-5</sup> | d=4.04 |
| 6D | WT VEH | Het VEH | 12s to 15s from outcome | SVM Accuracy | 1000, 1000 | 0.997 +/- 0.002 | 0.988 +/- 0.003 | Robust Cohen's d | 2.73 x 10<sup>-5</sup> | d=4.04 |
| 6D | Het VEH | Het CLNZ | 3s to 6 from outcome | SVM Accuracy | 1000, 1000 | 0.966 +/- 0.005 | 0.996 +/- 0.002 | Robust Cohen's d | 1.51 x 10<sup>-14</sup> | d=7.6 |
| 6D | Het VEH | Het postCLNZ | 3s to 6 from outcome | SVM Accuracy | 1000, 1000 | 0.966 +/- 0.005 | 0.998 +/- 0.001 | Robust Cohen's d | < 1 x 10<sup>-15</sup> | d=8.31 |
| 6D | Het VEH | Het CLNZ | 3s to 6 from outcome | SVM Accuracy | 1000, 1000 | 0.966 +/- 0.005 | 0.996 +/- 0.002 | Robust Cohen's d | 1.51 x 10<sup>-14</sup> | d=7.6 |
| 6D | Het VEH | Het postCLNZ | 3s to 6 from outcome | SVM Accuracy | 1000, 1000 | 0.966 +/- 0.005 | 0.998 +/- 0.001 | Robust Cohen's d | < 1 x 10<sup>-15</sup> | d=8.31 |
| 6D | WT VEH | Het VEH | 3s to 6 from outcome | SVM Accuracy | 1000, 1000 | 0.997 +/- 0.002 | 0.966 +/- 0.005 | Robust Cohen's d | 6.99 x 10<sup>-15</sup> | d=7.7 |
| 6D | WT VEH | Het VEH | 3s to 6 from outcome | SVM Accuracy | 1000, 1000 | 0.997 +/- 0.002 | 0.966 +/- 0.005 | Robust Cohen's d | 6.99 x 10<sup>-15</sup> | d=7.7 |
| 6D | Het VEH | Het CLNZ | 6s to 9s from outcome | SVM Accuracy | 1000, 1000 | 0.983 +/- 0.004 | 0.997 +/- 0.002 | Robust Cohen's d | 2.27 x 10<sup>-6</sup> | d=4.59 |
| 6D | Het VEH | Het postCLNZ | 6s to 9s from outcome | SVM Accuracy | 1000, 1000 | 0.983 +/- 0.004 | 0.997 +/- 0.002 | Robust Cohen's d | 4.80 x 10<sup>-7</sup> | d=4.9 |
| 6D | Het VEH | Het CLNZ | 6s to 9s from outcome | SVM Accuracy | 1000, 1000 | 0.983 +/- 0.004 | 0.997 +/- 0.002 | Robust Cohen's d | 2.27 x 10<sup>-6</sup> | d=4.59 |
| 6D | Het VEH | Het postCLNZ | 6s to 9s from outcome | SVM Accuracy | 1000, 1000 | 0.983 +/- 0.004 | 0.997 +/- 0.002 | Robust Cohen's d | 4.80 x 10<sup>-7</sup> | d=4.9 |
| 6D | WT VEH | Het VEH | 6s to 9s from outcome | SVM Accuracy | 1000, 1000 | 0.993 +/- 0.002 | 0.983 +/- 0.004 | Robust Cohen's d | 0.0005 | d=3.27 |
| 6D | WT VEH | Het VEH | 6s to 9s from outcome | SVM Accuracy | 1000, 1000 | 0.993 +/- 0.002 | 0.983 +/- 0.004 | Robust Cohen's d | 0.0005 | d=3.27 |
| 6D | Het VEH | Het CLNZ | 9s to 12s from outcome | SVM Accuracy | 1000, 1000 | 0.987 +/- 0.003 | 0.994 +/- 0.002 | Robust Cohen's d | 0.0062 | d=2.5 |
| 6D | Het VEH | Het postCLNZ | 9s to 12s from outcome | SVM Accuracy | 1000, 1000 | 0.987 +/- 0.003 | 0.996 +/- 0.002 | Robust Cohen's d | 0.0003 | d=3.42 |
| 6D | Het VEH | Het CLNZ | 9s to 12s from outcome | SVM Accuracy | 1000, 1000 | 0.987 +/- 0.003 | 0.994 +/- 0.002 | Robust Cohen's d | 0.0062 | d=2.5 |
| 6D | Het VEH | Het postCLNZ | 9s to 12s from outcome | SVM Accuracy | 1000, 1000 | 0.987 +/- 0.003 | 0.996 +/- 0.002 | Robust Cohen's d | 0.0003 | d=3.42 |

Supplementary Table 2: Figure Statistics
|  |  |  |  |  |  |  |  |  |  |  |
| --- | --- | --- | --- | --- | --- | --- | --- | --- | --- | --- |
| 6D | WT VEH | Het VEH | 9s to 12s from outcome | SVM Accuracy | 1000, 1000 | 0.998 +/- 0.001 | 0.987 +/- 0.003 | Robust Cohen's d | 7.51 x 10<sup>-6</sup> | d=4.33 |
| 6D | WT VEH | Het VEH | 9s to 12s from outcome | SVM Accuracy | 1000, 1000 | 0.998 +/- 0.001 | 0.987 +/- 0.003 | Robust Cohen's d | 7.51 x 10<sup>-6</sup> | d=4.33 |
| 7C | Het VEH | Het CLNZ | Early IA Correct ensemble | SVM accuracy | 1000, 1000 | 0.984 +/- 0.004 | 0.99 +/- 0.003 | Robust Cohen's d | 0.0316 | d=1.86 |
| 7C | Het VEH | Het postCLNZ | Early IA Correct ensemble | SVM accuracy | 1000, 1000 | 0.984 +/- 0.004 | 0.993 +/- 0.002 | Robust Cohen's d | 0.0014 | d=2.99 |
| 7C | WT VEH | Het VEH | Early IA Correct ensemble | SVM accuracy | 1000, 1000 | 0.984 +/- 0.004 | 0.984 +/- 0.004 | Robust Cohen's d | 0.4819 | d=0.05 |
| 7C | Het VEH | Het CLNZ | Late IA ensemble | SVM accuracy | 1000, 1000 | 0.935 +/- 0.007 | 0.937 +/- 0.007 | Robust Cohen's d | 0.4028 | d=0.25 |
| 7C | Het VEH | Het postCLNZ | Late IA ensemble | SVM accuracy | 1000, 1000 | 0.935 +/- 0.007 | 0.964 +/- 0.005 | Robust Cohen's d | 6.73 x 10<sup>-7</sup> | d=4.83 |
| 7C | WT VEH | Het VEH | Late IA ensemble | SVM accuracy | 1000, 1000 | 0.96 +/- 0.006 | 0.935 +/- 0.007 | Robust Cohen's d | 3.57 x 10<sup>-5</sup> | d=3.97 |
| 7G | Het VEH | Het CLNZ | Early RS Correct ensemble | SVM accuracy | 1000, 1000 | 0.952 +/- 0.006 | 0.965 +/- 0.005 | Robust Cohen's d | 0.0101 | d=2.32 |
| 7G | Het VEH | Het postCLNZ | Early RS Correct ensemble | SVM accuracy | 1000, 1000 | 0.952 +/- 0.006 | 0.977 +/- 0.004 | Robust Cohen's d | 3.70 x 10<sup>-7</sup> | d=4.95 |
| 7G | WT VEH | Het VEH | Early RS Correct ensemble | SVM accuracy | 1000, 1000 | 0.972 +/- 0.005 | 0.952 +/- 0.006 | Robust Cohen's d | 0.0001 | d=3.7 |
| 7G | Het VEH | Het CLNZ | Late IA ensemble | SVM accuracy | 1000, 1000 | 0.966 +/- 0.005 | 0.969 +/- 0.005 | Robust Cohen's d | 0.2856 | d=0.57 |
| 7G | Het VEH | Het postCLNZ | Late IA ensemble | SVM accuracy | 1000, 1000 | 0.966 +/- 0.005 | 0.969 +/- 0.006 | Robust Cohen's d | 0.2774 | d=0.59 |
| 7G | WT VEH | Het VEH | Late IA ensemble | SVM accuracy | 1000, 1000 | 0.97 +/- 0.005 | 0.966 +/- 0.005 | Robust Cohen's d | 0.1833 | d=0.9 |
| S1C | Het VEH | Het CLNZ | Early IA Correct stage | % of frames active | 4322, 3272 | 3.624 +/- 0.139 | 2.941 +/- 0.139 | Permutation Test | 0.0004 | N/A |
| S1C | Het VEH | Het postCLNZ | Early IA Correct stage | % of frames active | 4322, 4270 | 3.624 +/- 0.139 | 3.35 +/- 0.132 | Permutation Test | 0.1520 | N/A |
| S1C | WT VEH | WT CLNZ | Early IA Correct stage | % of frames active | 4952, 4352 | 2.679 +/- 0.105 | 2.793 +/- 0.122 | Permutation Test | 0.4786 | N/A |
| S1C | WT VEH | Het VEH | Early IA Correct stage | % of frames active | 4952, 4322 | 2.679 +/- 0.105 | 3.624 +/- 0.139 | Permutation Test | 0.0002 | N/A |
| S1C | Het VEH | Het CLNZ | Early IA Error stage | % of frames active | 1355, 2198 | 3.396 +/- 0.236 | 3.081 +/- 0.166 | Permutation Test | 0.2536 | N/A |
| S1C | Het VEH | Het postCLNZ | Early IA Error stage | % of frames active | 1355, 1615 | 3.396 +/- 0.236 | 4.027 +/- 0.233 | Permutation Test | 0.0578 | N/A |
| S1C | WT VEH | WT CLNZ | Early IA Error stage | % of frames active | 1798, 1878 | 3.089 +/- 0.203 | 3.104 +/- 0.202 | Permutation Test | 0.9781 | N/A |
| S1C | WT VEH | Het VEH | Early IA Error stage | % of frames active | 1798, 1355 | 3.089 +/- 0.203 | 3.396 +/- 0.236 | Permutation Test | 0.3308 | N/A |
| S1C | Het VEH | Het CLNZ | Early RS Correct stage | % of frames active | 2552, 3017 | 2.235 +/- 0.14 | 1.955 +/- 0.115 | Permutation Test | 0.1120 | N/A |
| S1C | Het VEH | Het postCLNZ | Early RS Correct stage | % of frames active | 2552, 3358 | 2.235 +/- 0.14 | 2.072 +/- 0.121 | Permutation Test | 0.3904 | N/A |
| S1C | WT VEH | WT CLNZ | Early RS Correct stage | % of frames active | 3989, 2785 | 1.908 +/- 0.103 | 2.071 +/- 0.126 | Permutation Test | 0.3312 | N/A |
| S1C | WT VEH | Het VEH | Early RS Correct stage | % of frames active | 3989, 2552 | 1.908 +/- 0.103 | 2.235 +/- 0.14 | Permutation Test | 0.0534 | N/A |
| S1C | Het VEH | Het CLNZ | Early RS Error stage | % of frames active | 2918, 2453 | 2.507 +/- 0.12 | 2.307 +/- 0.134 | Permutation Test | 0.2672 | N/A |
| S1C | Het VEH | Het postCLNZ | Early RS Error stage | % of frames active | 2918, 2527 | 2.507 +/- 0.12 | 2.44 +/- 0.144 | Permutation Test | 0.7285 | N/A |
| S1C | WT VEH | WT CLNZ | Early RS Error stage | % of frames active | 2761, 2893 | 2.057 +/- 0.129 | 2.061 +/- 0.124 | Permutation Test | 0.9883 | N/A |
| S1C | WT VEH | Het VEH | Early RS Error stage | % of frames active | 2761, 2918 | 2.057 +/- 0.129 | 2.507 +/- 0.12 | Permutation Test | 0.0104 | N/A |
| S1C | Het VEH | Het CLNZ | Late IA stage | % of frames active | 5263, 5470 | 2.713 +/- 0.106 | 2.104 +/- 0.089 | Permutation Test | 0.0002 | N/A |
| S1C | Het VEH | Het postCLNZ | Late IA stage | % of frames active | 5263, 5754 | 2.713 +/- 0.106 | 2.512 +/- 0.094 | Permutation Test | 0.1588 | N/A |
| S1C | WT VEH | WT CLNZ | Late IA stage | % of frames active | 6659, 6230 | 2.133 +/- 0.085 | 2.103 +/- 0.084 | Permutation Test | 0.7981 | N/A |
| S1C | WT VEH | Het VEH | Late IA stage | % of frames active | 6659, 5263 | 2.133 +/- 0.085 | 2.713 +/- 0.106 | Permutation Test | 0.0002 | N/A |
| S1C | Het VEH | Het CLNZ | Late RS stage | % of frames active | 5190, 5200 | 1.393 +/- 0.074 | 1.551 +/- 0.076 | Permutation Test | 0.1274 | N/A |
| S1C | Het VEH | Het postCLNZ | Late RS stage | % of frames active | 5190, 5083 | 1.393 +/- 0.074 | 1.512 +/- 0.079 | Permutation Test | 0.2684 | N/A |
| S1C | WT VEH | WT CLNZ | Late RS stage | % of frames active | 6619, 5644 | 1.406 +/- 0.065 | 1.754 +/- 0.08 | Permutation Test | 0.0002 | N/A |
| S1C | WT VEH | Het VEH | Late RS stage | % of frames active | 6619, 5190 | 1.406 +/- 0.065 | 1.393 +/- 0.074 | Permutation Test | 0.8969 | N/A |
| S1D | Het VEH | Het CLNZ | Early IA Correct stage | Normalized event rate | 874, 866 | 0.045 +/- 0.002 | 0.036 +/- 0.002 | Mann-Whitney U | 3.38 x 10<sup>-6</sup> | U=425312.5 |
| S1D | Het VEH | Het postCLNZ | Early IA Correct stage | Normalized event rate | 874, 937 | 0.045 +/- 0.002 | 0.043 +/- 0.002 | Mann-Whitney U | 0.3238 | U=420160 |

Supplementary Table 2: Figure Statistics
|  |  |  |  |  |  |  |  |  |  |  |
| --- | --- | --- | --- | --- | --- | --- | --- | --- | --- | --- |
| S1D | WT VEH | WT CLNZ | Early IA Correct stage | Normalized event rate | 1080, 1019 | 0.033 +/- 0.002 | 0.034 +/- 0.002 | Mann-Whitney U | 0.1633 | U=568662.5 |
| S1D | WT VEH | Het VEH | Early IA Correct stage | Normalized event rate | 1080, 874 | 0.033 +/- 0.002 | 0.045 +/- 0.002 | Mann-Whitney U | 0.0002 | U=426567.5 |
| S1D | Het VEH | Het CLNZ | Early IA Error stage | Normalized event rate | 749, 742 | 0.043 +/- 0.003 | 0.039 +/- 0.003 | Mann-Whitney U | 0.0191 | U=260184.5 |
| S1D | Het VEH | Het postCLNZ | Early IA Error stage | Normalized event rate | 749, 689 | 0.043 +/- 0.003 | 0.05 +/- 0.003 | Mann-Whitney U | 0.0002 | U=231817 |
| S1D | WT VEH | WT CLNZ | Early IA Error stage | Normalized event rate | 870, 838 | 0.037 +/- 0.003 | 0.038 +/- 0.003 | Mann-Whitney U | 0.2965 | U=355351.5 |
| S1D | WT VEH | Het VEH | Early IA Error stage | Normalized event rate | 870, 749 | 0.037 +/- 0.003 | 0.043 +/- 0.003 | Mann-Whitney U | 0.1028 | U=312569 |
| S1D | Het VEH | Het CLNZ | Early RS Correct stage | Normalized event rate | 874, 866 | 0.029 +/- 0.002 | 0.025 +/- 0.002 | Mann-Whitney U | 0.6269 | U=373956 |
| S1D | Het VEH | Het postCLNZ | Early RS Correct stage | Normalized event rate | 874, 937 | 0.029 +/- 0.002 | 0.027 +/- 0.002 | Mann-Whitney U | 0.9214 | U=408507 |
| S1D | WT VEH | WT CLNZ | Early RS Correct stage | Normalized event rate | 1080, 1019 | 0.024 +/- 0.002 | 0.025 +/- 0.002 | Mann-Whitney U | 0.3309 | U=561740.5 |
| S1D | WT VEH | Het VEH | Early RS Correct stage | Normalized event rate | 1080, 874 | 0.024 +/- 0.002 | 0.029 +/- 0.002 | Mann-Whitney U | 0.3211 | U=461303.5 |
| S1D | Het VEH | Het CLNZ | Early RS Error stage | Normalized event rate | 874, 866 | 0.03 +/- 0.002 | 0.029 +/- 0.002 | Mann-Whitney U | 0.0029 | U=407292.5 |
| S1D | Het VEH | Het postCLNZ | Early RS Error stage | Normalized event rate | 874, 937 | 0.03 +/- 0.002 | 0.029 +/- 0.002 | Mann-Whitney U | 2.05 x 10<sup>-5</sup> | U=452785.5 |
| S1D | WT VEH | WT CLNZ | Early RS Error stage | Normalized event rate | 1080, 1019 | 0.026 +/- 0.002 | 0.025 +/- 0.002 | Mann-Whitney U | 0.0112 | U=521227.5 |
| S1D | WT VEH | Het VEH | Early RS Error stage | Normalized event rate | 1080, 874 | 0.026 +/- 0.002 | 0.03 +/- 0.002 | Mann-Whitney U | 2.67 x 10<sup>-18</sup> | U=376801 |
| S1D | Het VEH | Het CLNZ | Late IA stage | Normalized event rate | 874, 866 | 0.033 +/- 0.002 | 0.026 +/- 0.002 | Mann-Whitney U | 0.0131 | U=403685.5 |
| S1D | Het VEH | Het postCLNZ | Late IA stage | Normalized event rate | 874, 937 | 0.033 +/- 0.002 | 0.032 +/- 0.002 | Mann-Whitney U | 0.5868 | U=415356 |
| S1D | WT VEH | WT CLNZ | Late IA stage | Normalized event rate | 1080, 1019 | 0.026 +/- 0.001 | 0.026 +/- 0.001 | Mann-Whitney U | 0.8545 | U=547834 |
| S1D | WT VEH | Het VEH | Late IA stage | Normalized event rate | 1080, 874 | 0.026 +/- 0.001 | 0.033 +/- 0.002 | Mann-Whitney U | 0.0012 | U=433286.5 |
| S1D | Het VEH | Het CLNZ | Late RS stage | Normalized event rate | 874, 866 | 0.018 +/- 0.001 | 0.02 +/- 0.001 | Mann-Whitney U | 0.8657 | U=376813.5 |
| S1D | Het VEH | Het postCLNZ | Late RS stage | Normalized event rate | 874, 937 | 0.018 +/- 0.001 | 0.019 +/- 0.001 | Mann-Whitney U | 0.5190 | U=416003 |
| S1D | WT VEH | WT CLNZ | Late RS stage | Normalized event rate | 1080, 1019 | 0.017 +/- 0.001 | 0.021 +/- 0.001 | Mann-Whitney U | 0.0897 | U=528759.5 |
| S1D | WT VEH | Het VEH | Late RS stage | Normalized event rate | 1080, 874 | 0.017 +/- 0.001 | 0.018 +/- 0.001 | Mann-Whitney U | 0.4973 | U=464277.5 |
| S1J | Het VEH | Het CLNZ | Early IA Correct ensemble | % of cells active in trial | 27, 21 | 68.779 +/- 3.054 | 67.349 +/- 3.494 | Mann-Whitney U | 0.7313 | U=300.5 |
| S1J | Het VEH | Het postCLNZ | Early IA Correct ensemble | % of cells active in trial | 27, 26 | 68.779 +/- 3.054 | 68.004 +/- 2.282 | Mann-Whitney U | 0.6119 | U=380 |
| S1J | WT VEH | WT CLNZ | Early IA Correct ensemble | % of cells active in trial | 29, 21 | 57.961 +/- 3.241 | 60.137 +/- 3.296 | Mann-Whitney U | 0.9216 | U=299 |
| S1J | WT VEH | Het VEH | Early IA Correct ensemble | % of cells active in trial | 29, 27 | 57.961 +/- 3.241 | 68.779 +/- 3.054 | Mann-Whitney U | 0.0225 | U=252 |
| S1J | Het VEH | Het CLNZ | Early IA Error ensemble | % of cells active in trial | 9, 14 | 84.259 +/- 6.818 | 77.687 +/- 4.804 | Mann-Whitney U | 0.3240 | U=79 |
| S1J | Het VEH | Het postCLNZ | Early IA Error ensemble | % of cells active in trial | 9, 9 | 84.259 +/- 6.818 | 80.599 +/- 4.081 | Mann-Whitney U | 0.3263 | U=52 |
| S1J | WT VEH | WT CLNZ | Early IA Error ensemble | % of cells active in trial | 11, 9 | 75.518 +/- 6.481 | 70.273 +/- 6.383 | Mann-Whitney U | 0.5918 | U=57 |
| S1J | WT VEH | Het VEH | Early IA Error ensemble | % of cells active in trial | 11, 9 | 75.518 +/- 6.481 | 84.259 +/- 6.818 | Mann-Whitney U | 0.3514 | U=37 |
| S1J | Het VEH | Het CLNZ | Early RS Correct ensemble | % of cells active in trial | 16, 19 | 75.243 +/- 3.984 | 66.895 +/- 3.938 | Mann-Whitney U | 0.1898 | U=192 |
| S1J | Het VEH | Het postCLNZ | Early RS Correct ensemble | % of cells active in trial | 16, 20 | 75.243 +/- 3.984 | 69.251 +/- 5.711 | Mann-Whitney U | 0.5869 | U=177.5 |
| S1J | WT VEH | WT CLNZ | Early RS Correct ensemble | % of cells active in trial | 24, 13 | 59.626 +/- 4.437 | 66.298 +/- 3.419 | Mann-Whitney U | 0.5449 | U=136.5 |
| S1J | WT VEH | Het VEH | Early RS Correct ensemble | % of cells active in trial | 24, 16 | 59.626 +/- 4.437 | 75.243 +/- 3.984 | Mann-Whitney U | 0.0233 | U=109.5 |
| S1J | Het VEH | Het CLNZ | Early RS Error ensemble | % of cells active in trial | 19, 16 | 73.509 +/- 3.914 | 71.894 +/- 4.759 | Mann-Whitney U | 0.7517 | U=162 |
| S1J | Het VEH | Het postCLNZ | Early RS Error ensemble | % of cells active in trial | 19, 15 | 73.509 +/- 3.914 | 73.677 +/- 4.206 | Mann-Whitney U | 0.9583 | U=140.5 |
| S1J | WT VEH | WT CLNZ | Early RS Error ensemble | % of cells active in trial | 14, 14 | 64.4 +/- 7.031 | 69.345 +/- 4.398 | Mann-Whitney U | 0.7992 | U=92 |
| S1J | WT VEH | Het VEH | Early RS Error ensemble | % of cells active in trial | 14, 19 | 64.4 +/- 7.031 | 73.509 +/- 3.914 | Mann-Whitney U | 0.3789 | U=108.5 |
| S1J | Het VEH | Het CLNZ | Late IA ensemble | % of cells active in trial | 34, 35 | 60.508 +/- 3.019 | 56.412 +/- 3.538 | Mann-Whitney U | 0.3831 | U=668 |
| S1J | Het VEH | Het postCLNZ | Late IA ensemble | % of cells active in trial | 34, 34 | 60.508 +/- 3.019 | 63.889 +/- 2.302 | Mann-Whitney U | 0.3535 | U=502 |

Supplementary Table 2: Figure Statistics
|  |  |  |  |  |  |  |  |  |  |  |
| --- | --- | --- | --- | --- | --- | --- | --- | --- | --- | --- |
| S1J | WT VEH | WT CLNZ | Late IA ensemble | % of cells active in trial | 39, 30 | 49.902 +/- 2.914 | 56.337 +/- 2.908 | Mann-Whitney U | 0.2516 | U=490 |
| S1J | WT VEH | Het VEH | Late IA ensemble | % of cells active in trial | 39, 34 | 49.902 +/- 2.914 | 60.508 +/- 3.019 | Mann-Whitney U | 0.0324 | U=469.5 |
| S1J | Het VEH | Het CLNZ | Late RS ensemble | % of cells active in trial | 33, 33 | 59.033 +/- 4.011 | 67.397 +/- 4.445 | Mann-Whitney U | 0.1289 | U=426 |
| S1J | Het VEH | Het postCLNZ | Late RS ensemble | % of cells active in trial | 33, 30 | 59.033 +/- 4.011 | 54.633 +/- 4.831 | Mann-Whitney U | 0.5253 | U=541.5 |
| S1J | WT VEH | WT CLNZ | Late RS ensemble | % of cells active in trial | 39, 27 | 48.077 +/- 5.145 | 54.749 +/- 3.246 | Mann-Whitney U | 0.2980 | U=447 |
| S1J | WT VEH | Het VEH | Late RS ensemble | % of cells active in trial | 39, 33 | 48.077 +/- 5.145 | 59.033 +/- 4.011 | Mann-Whitney U | 0.1228 | U=507.5 |
| S2C | Het VEH | Het CLNZ | Early RS Correct activity | % samples classified | 1000, 1000 | 53.301 +/- 1.745 | 65.751 +/- 1.952 | Robust Cohen's d | $8.81 \times 10^{-12}$ | d=6.72 |
| S2C | Het VEH | Het postCLNZ | Early RS Correct activity | % samples classified | 1000, 1000 | 53.301 +/- 1.745 | 77.192 +/- 1.665 | Robust Cohen's d | $< 1 \times 10^{-15}$ | d=14.01 |
| S2C | WT VEH | WT CLNZ | Early RS Correct activity | % samples classified | 1000, 1000 | 54.686 +/- 1.475 | 78.68 +/- 1.198 | Robust Cohen's d | $< 1 \times 10^{-15}$ | d=17.86 |
| S2C | WT VEH | Het VEH | Early RS Correct activity | % samples classified | 1000, 1000 | 54.686 +/- 1.475 | 53.301 +/- 1.745 | Robust Cohen's d | 0.1956 | d=0.86 |
| S2C | Het VEH | Het CLNZ | Early RS Error activity | % samples classified | 1000, 1000 | 18.218 +/- 3.014 | 44.32 +/- 1.329 | Robust Cohen's d | $< 1 \times 10^{-15}$ | d=11.21 |
| S2C | Het VEH | Het postCLNZ | Early RS Error activity | % samples classified | 1000, 1000 | 18.218 +/- 3.014 | 62.795 +/- 1.94 | Robust Cohen's d | $< 1 \times 10^{-15}$ | d=17.59 |
| S2C | WT VEH | WT CLNZ | Early RS Error activity | % samples classified | 1000, 1000 | 96.159 +/- 1.06 | 47.966 +/- 1.19 | Robust Cohen's d | $< 1 \times 10^{-15}$ | d=42.78 |
| S2C | WT VEH | Het VEH | Early RS Error activity | % samples classified | 1000, 1000 | 96.159 +/- 1.06 | 18.218 +/- 3.014 | Robust Cohen's d | $< 1 \times 10^{-15}$ | d=34.5 |
| S2D | Het VEH | Het CLNZ | Early IA Error activity | % samples classified | 1000, 1000 | 45.04 +/- 1.236 | 48.969 +/- 3.213 | Robust Cohen's d | 0.0533 | d=1.61 |
| S2D | Het VEH | Het postCLNZ | Early IA Error activity | % samples classified | 1000, 1000 | 45.04 +/- 1.236 | 56.417 +/- 4.19 | Robust Cohen's d | 0.0001 | d=3.68 |
| S2D | WT VEH | WT CLNZ | Early IA Error activity | % samples classified | 1000, 1000 | 50.411 +/- 2.237 | 24.626 +/- 1.763 | Robust Cohen's d | $< 1 \times 10^{-15}$ | d=12.8 |
| S2D | WT VEH | Het VEH | Early IA Error activity | % samples classified | 1000, 1000 | 50.411 +/- 2.237 | 45.04 +/- 1.236 | Robust Cohen's d | 0.0015 | d=2.97 |
| S2D | Het VEH | Het CLNZ | Early RS Error activity | % samples classified | 1000, 1000 | 51.353 +/- 1.801 | 88.192 +/- 1.915 | Robust Cohen's d | $< 1 \times 10^{-15}$ | d=19.82 |
| S2D | Het VEH | Het postCLNZ | Early RS Error activity | % samples classified | 1000, 1000 | 51.353 +/- 1.801 | 91.201 +/- 1.993 | Robust Cohen's d | $< 1 \times 10^{-15}$ | d=20.98 |
| S2D | WT VEH | WT CLNZ | Early RS Error activity | % samples classified | 1000, 1000 | 86.599 +/- 1.466 | 70.517 +/- 4.155 | Robust Cohen's d | $1.22 \times 10^{-7}$ | d=5.16 |
| S2D | WT VEH | Het VEH | Early RS Error activity | % samples classified | 1000, 1000 | 86.599 +/- 1.466 | 51.353 +/- 1.801 | Robust Cohen's d | $< 1 \times 10^{-15}$ | d=21.47 |
| S2E | Het VEH | Het CLNZ | Early IA Error activity | % samples classified | 1000, 1000 | 78.321 +/- 7.817 | 61.475 +/- 2.035 | Robust Cohen's d | 0.0016 | d=2.95 |
| S2E | Het VEH | Het postCLNZ | Early IA Error activity | % samples classified | 1000, 1000 | 78.321 +/- 7.817 | 50.653 +/- 1.833 | Robust Cohen's d | $5.49 \times 10^{-7}$ | d=4.87 |
| S2E | WT VEH | WT CLNZ | Early IA Error activity | % samples classified | 1000, 1000 | 11.615 +/- 1.265 | 58.882 +/- 1.285 | Robust Cohen's d | $< 1 \times 10^{-15}$ | d=37.08 |
| S2E | WT VEH | Het VEH | Early IA Error activity | % samples classified | 1000, 1000 | 11.615 +/- 1.265 | 78.321 +/- 7.817 | Robust Cohen's d | $< 1 \times 10^{-15}$ | d=11.91 |
| S2E | Het VEH | Het CLNZ | Early RS Error activity | % samples classified | 1000, 1000 | 47.708 +/- 1.71 | 75.112 +/- 2.076 | Robust Cohen's d | $< 1 \times 10^{-15}$ | d=14.41 |
| S2E | Het VEH | Het postCLNZ | Early RS Error activity | % samples classified | 1000, 1000 | 47.708 +/- 1.71 | 39.044 +/- 2.292 | Robust Cohen's d | $9.14 \times 10^{-6}$ | d=4.28 |
| S2E | WT VEH | WT CLNZ | Early RS Error activity | % samples classified | 1000, 1000 | 43.873 +/- 1.851 | 23.5 +/- 1.636 | Robust Cohen's d | $< 1 \times 10^{-15}$ | d=11.66 |
| S2E | WT VEH | Het VEH | Early RS Error activity | % samples classified | 1000, 1000 | 43.873 +/- 1.851 | 47.708 +/- 1.71 | Robust Cohen's d | 0.0157 | d=2.15 |

